# Persistence of Non-industrial Gut Microbiota Discriminates Health and Disease States in Industrial Populations

**DOI:** 10.64898/2026.11.17.767315

**Authors:** Neil G. O. Ibata, Amar Sarkar, Esther S. Brielle, Lasya R. Penumarthi, Andrew H. Moeller, Rachel N. Carmody

## Abstract

Industrialization has sharply reduced gut microbial diversity, possibly affecting human health. To isolate the effects of industrialization, we assembled 18,075 human metagenomes spanning hunter-gatherer, agro-pastoral, and industrial lifestyles. We confirmed individual-level diversity reductions in industrial populations, but also found that nearly all known human-associated microbes exhibit population-level persistence. Previously hypothesized ‘vanishing’ microbes are reduced from high prevalence in non-industrial populations to above-median prevalence in industrial populations. Species-level signatures in industrial case–control cohorts suggest that harboring ‘vanishing’ taxa can be associated with health or pathology, depending on the condition. Controlling for background prevalence, ‘vanishing’ microbes were more likely associated with cases than controls in obesity, both ‘vanishing’ microbes and those now ‘blossoming’ in industrialized populations were associated with glucose intolerance, while ‘blossoming’ taxa were associated with inflammatory bowel disease. We conclude that most microbes have not gone extinct under industrialization, and that the health effects of lifestyle-associated taxa are context-dependent.

---

The gut microbiome plays a critical role in shaping host physiology, development, and health (*1–4*). Humans and our gut microbiomes have co-diversified and co-speciated over time (*5–8*). Indeed, contemporary non-industrial populations on different continents share a large number of microbial taxa, with average nucleotide identities suggesting that common microbiome members co-migrated with human hosts out of Africa and have operated in stable association with these populations for tens of thousands of years (*8*). However, this situation is different for populations that have experienced industrialization, which has elicited numerous significant lifestyle changes that may deplete or alter the microbiota, including dietary ultra-processing (*9*, *10*), antibiotic use (*11*), and caesarean births (*12*). These changes have led to reductions in gut microbiota taxonomic diversity within individual human hosts (*13*). Losses of microbial taxa found at high prevalence and abundance in non-industrial contexts have fueled concerns that we have driven essential health-promoting microbes extinct in industrial populations (*13–17*). This concern has in turn prompted proposals that industrialized gut microbiomes require restoration toward preindustrial profiles to mitigate the negative health consequences of ‘missing’ microbes (*14*, *16*, *17*).

However, the extent to which industrial lifestyles have caused local extinctions of gut microbes remains unclear. Murine experiments suggest that sustained, multigenerational exposure to westernized, fiber-poor diets can drive microbes specializing in the digestion of complex carbohydrates to below-detection-threshold levels (*18*). Recent work in humans has also identified numerous ‘vanishing’ gut microbes (referred to as <u>v</u>olatile and/or <u>a</u>ssociated <u>n</u>egatively with industrialized <u>s</u>ocieties of <u>h</u>umans or ‘VANISH’ taxa) (*19*). These are commonly found in hunter-gatherers, for which the Hadza of Tanzania have served as the primary reference (*20–24*), but are rare or absent in industrial individuals (*20*). These studies have also revealed a smaller number of microbes that are present at low prevalence in hunter-gatherer individuals but show substantial ‘blossoming’ (referred to as <u>blo</u>om or <u>s</u>elected in <u>s</u>ocieties of <u>u</u>rbanization/<u>m</u>odernization or ‘BloSSUM’ taxa) in industrial individuals (*20*).

The health consequences of such microbial losses likewise remain unclear. Microbiomes present in hunter-gatherers are commonly used as models of an ancestral-like state to which human physiology is assumed to be better adapted (*15*). The loss of microbial taxa present in hunter-gatherer microbiomes from industrial microbiomes is therefore theorized to disrupt host-microbiome interactions that promote human fitness (*16*, *17*). Moreover, such loss has been argued to contribute to non-communicable diseases now rising in prevalence in industrial populations, including gastrointestinal and metabolic syndromes (*13*, *25–27*). However, an alternative but not mutually exclusive hypothesis is that the gut microbiome, owing to its extraordinary plasticity, may in some cases help hosts adapt to rapidly changing environments (*15*). Two predictions arise from this alternative hypothesis: 1) industrialized microbiomes may sometimes be associated with health rather than disease in industrialized contexts, and 2), conversely, deviations from the typical industrialized microbiome may be associated with disease rather than health.

Here we evaluate these hypotheses by assembling one of the largest metagenomic datasets to date, capturing studies of the human gut microbiome across hunter-gatherers, agriculturalists, pastoralists, and industrials in 42 countries (*N* = 18,075). We then use this expansive dataset to examine patterns of gut microbiota diversity at the individual and population levels, and test associations between patterns of diversity and health outcomes.

## Assembly of a metagenomic mega-dataset for studies of human lifestyle variation

To investigate how lifestyle impacts gut microbiota diversity and composition, we assembled one of the largest known metagenomic datasets. Our dataset includes 83 studies that sequenced the fecal microbiome of human populations across 42 countries, spanning hunter-gatherer lifestyles, agricultural or pastoral (collectively referred to here as “agro-pastoralist”) lifestyles, and industrial lifestyles (**Fig. 1, figs. S1–S4, Supplementary Text 1**, **Tables S1–S3**, and **Data S1** that was used to generate all figures in this paper). Our database consists of 18,075 deeply sequenced fecal microbiome samples (metagenomes), of which 401 were classified as hunter-gatherer (8.02 trillion bases), 3,697 as agro-pastoralist (35.3 trillion bases), and 13,977 as industrial (60.4 trillion bases) (**Data S2**), all of which we processed from raw reads using a common alignment and quality-control workflow. Our database captures a substantially greater number of samples than used in previous studies. For comparison, one of the most comprehensive databases previously assembled to compare microbiomes across lifestyles (now included as a subset of our dataset) included 267 hunter-gatherer metagenomes (4.9 trillion bases), 583 agro-pastoralist metagenomes (4.8 trillion bases), and 950 industrial metagenomes (4.3 trillion bases) (*20*) (**Data S2**).

**Figure 1.**
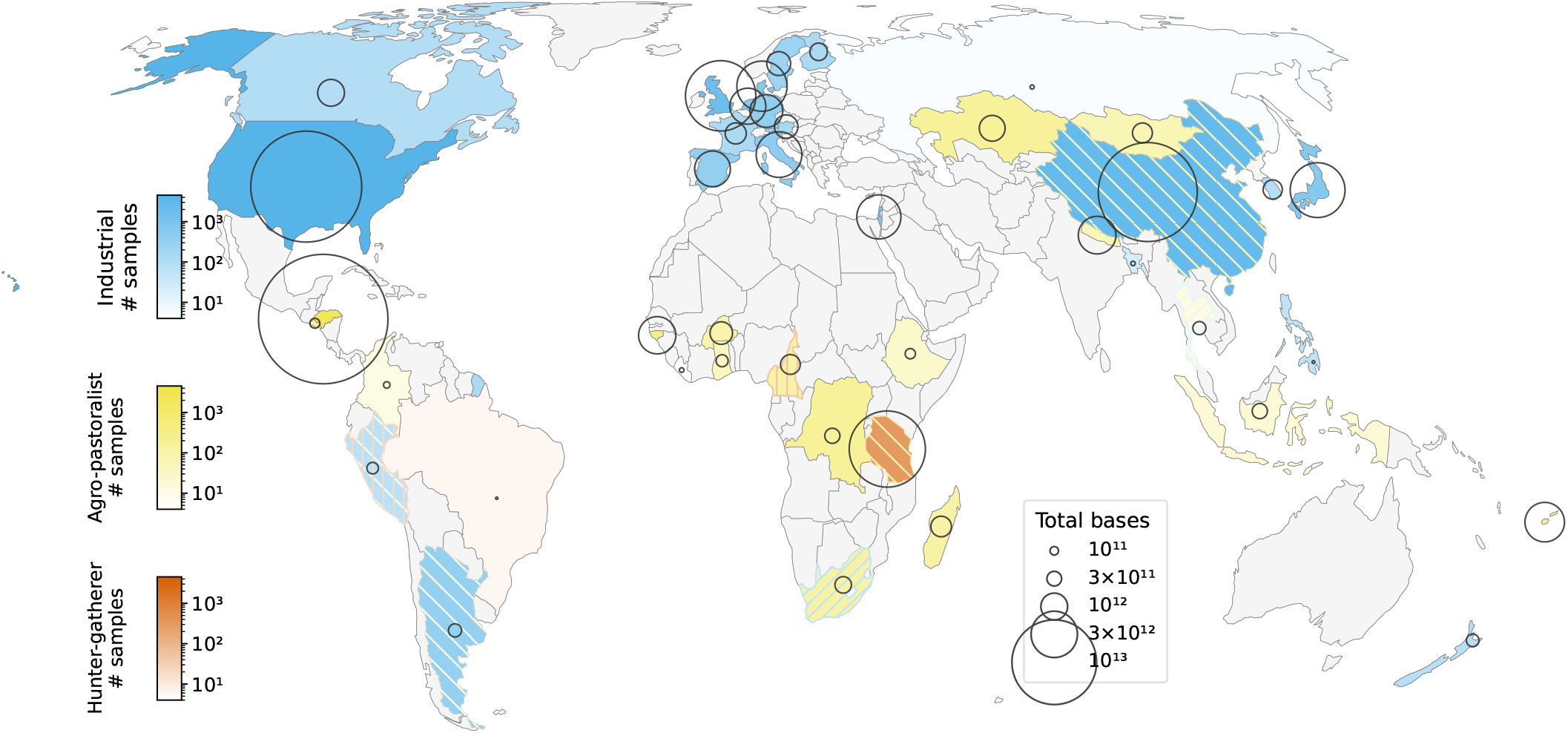
Global distribution of human metagenome studies. Visualization of the global sequencing depth and breadth of gut microbial metagenomes representing populations with different lifestyles (blue: industrial; yellow: agro-pastoralist; red: hunter-gatherer) from 42 countries (*N*=18,075). When two or three lifestyle categories are present in the same country (e.g. Tanzania, Peru), the stripes indicate the less prevalent lifestyle(s). Deeper shades show more broadly sequenced populations; wider circles correspond to populations with greater sequencing effort (depth and breadth). Most of the global microbiome sequencing effort thus far has been concentrated in 5 regions: North America, Honduras, Western Europe, Tanzania (specifically, Hadza hunter-gatherers), and East Asia. We used an equal-area (Equal Earth) projection to better illustrate the global discrepancy in coverage between industrial and non-industrial populations.

Because several studies resampled the same individuals longitudinally, these 18,075 samples represent 14,624 hosts (401 hunter-gatherer samples from 238 hosts, 3,697 agro-pastoralist samples from 3,539 hosts, and 13,977 industrial samples from 10,847 hosts; **Tables S1–S2**). Longitudinal samples were retained in the lifestyle analyses but reduced to one sample per host in the health analyses described below. This is because treating them as independent would inflate the evidence to detect an association (**Supplementary Text 1C**).

We mapped the reads to one of three reference databases: 1) the Carter catalogue, which includes 3,971 species derived primarily from non-industrial Hadza and Nepali individuals plus a subset of the Unified Human Gastrointestinal Genome (UHGG) database (*20*, *28*). 2) The Leviatan catalogue, which includes 3,594 species derived primarily from industrial populations (mainly an Israeli cohort) (*29*). 3) A dereplicated “combined” catalogue that we constructed from these two published catalogues (**Supplementary Text 2, Table S4**). Of these bases, 41% survived trimming, removal of host reads, mapping to the combined catalogue, and alignment at MAPQ ≥30 (an extremely conservative requirement to avoid potential false positive microbial detections). In all three catalogues, industrial samples map a higher share of their bases versus non-industrial samples: 46.0% of industrial bases are retained versus 35.7% of hunter-gatherer and 34.7% of agro-pastoralist bases (**Fig. S1**). All depths quoted subsequently are bases/reads successfully mapped against our combined reference catalogue.

## Correction for reference bias and sequencing effort enables robust quantification of within-sample microbiota diversity

Testing for extinction at the population level presents the proverbial difficulty of finding a needle in a haystack. Because low abundance microbes are more difficult to detect, distinguishing microbes that are rare from those that are truly absent presents a formidable challenge. To effectively identify microbes that are rare but not absent at the population level, we targeted populations that were deeply sequenced, broadly sampled, or both (**Supplementary Text 1).** Because reference-based metagenomics enables the detection of much lower-abundance taxa than reference-free metagenomics, we reprocessed all retained metagenomes through a common pipeline, using a reference-based approach.

We designed our analyses to resolve two issues that we have identified as affecting prior large-scale studies of human gut microbiome diversity. The first of these issues was that the choice of reference database can introduce substantial bias (**Supplementary Text 2**). Therefore, we aligned the reads for each sample to two different high-quality reference databases with opposing ascertainment biases. The first of these is the Carter catalogue with 3,971 taxa inferred from species-level genome bins (SGBs; clusters of genomes sharing ≥95% average nucleotide identity, a working definition of a bacterial species) (**Data S3**), which is enriched in bacterial genomes identified in hunter-gatherers, and is therefore enriched for non-industrial discovery (*20*). The second is the Leviatan catalogue with 3,594 SGBs (**Data S3**), which is enriched in bacterial genomes identified in industrial populations, and is therefore enriched for industrial discovery (*29*) (**Supplementary Text 2**, **Table S4**). As expected, we found that the hunter-gatherer-enriched Carter catalogue recovered more SGBs from hunter-gatherer samples than did the Leviatan catalogue (**fig. S1B**), while the two performed nearly equally on industrial samples. In an effort to compensate for such biases, we created a combined reference with 5,081 unique SGBs by clustering the SGBs into 95% average nucleotide identity (ANI) clusters and finding the highest-coverage SGB per cluster. Combining the catalogues measurably improved the number of bases mapping to a reference (**fig. S1B**).

Second, we identified that previous attempts to quantify the microbial taxa in different human populations (*20*) have been significantly biased by differences in sequencing depth (number of reads per sample) across populations (**fig. S5–S8; Supplementary Text 2**). This bias can affect inferences about lost diversity in industrial populations because hunter-gatherer and agro-pastoralist populations have generally been sequenced more deeply than samples from industrial populations (hunter-gatherer: mean 7.15±0.32 Gb/sample; agro-pastoralist: mean 3.31±0.04 Gb/sample; industrial: mean 1.99±0.01 Gb/sample) (**Fig. S4**). Some studies have accounted for differences in sequencing depth through rarefaction, which is statistically problematic (*30*) and especially inappropriate if we seek to examine the tail of rare microbial taxa that would be most susceptible to extinction. Recent large-scale metagenomic comparisons have not suggested an unambiguous solution to this problem, making it unclear whether increased within-individual diversity in hunter-gatherer and agro-pastoralist populations reflects true biological variation rather than increased sequencing effort.

In our reference-based framework, sequencing depth was strongly associated with recoverable diversity (Spearman’s *ρ*=0.682, *p*<10⁻³⁰⁰; *N*=18,075) (**fig. S9**). We therefore additionally corrected for sequencing depth in all analyses, using a geometric-mean-target weighting (GMW) method (a variant of inverse probability weighting). This method re-weights microbial samples rather than eliminating data by down-sampling (**figs. S5–S13; Supplementary Text 3**). Studies also differ in read length, which affects how many reads map to a reference and therefore the richness that we observe. We tested whether read length could confound the depth bias further, but we found only a small positive association between read length and richness, inconsistent between industrial and non-industrial samples and attributable to batch effects (**fig. S14**). Therefore, we discard read length as a confounder and treat sequencing depth as the dominant technical confound, which we then correct for throughout.

Evidence for microbial detection relies on sequenced DNA reads mapping sufficiently broadly onto the reference bacterial genome. The fraction of the bacterial genome that is mapped to by any DNA read is referred to as breadth of coverage. We used a standard 10% breadth coverage threshold (*31*, *32*), based on the biological rationale that 95% of plasmids, which are not informative for taxonomic identification due to their frequent transfer across species, are typically smaller than this (<300kb or 10% breadth coverage for a typical 3Mb genome (*33*)). Full model specification and robustness analyses (verifying similar conclusions under 30% and 50% thresholds) are provided in **Supplementary Text 4** and **Figs. S15–S17**.

## Individuals in industrial populations have reduced gut microbiota diversity

We used our GMW method (**Supplementary Text 3**) to directly compare microbiota richness among groups with systematically different sequencing depths **(figs. S5–S13)**. Our analysis confirmed that individuals in industrial populations harbor lower-diversity gut microbiota, with industrial guts exhibiting 61.4% of the richness found in non-industrial populations (95% CI 60.1–62.7%, 2,000 bootstrap resamples; Mann–Whitney U on the depth-corrected richness, p<10^−300^) (**Fig. 2**, **figs. S9, S10, S13, Supplementary Text 3**). Interestingly, gut microbiota of agro-pastoralist populations differed fundamentally from those of hunter-gatherer and industrial populations, rather than contributing to a graded spectrum of lifestyles. We projected agro-pastoralist samples onto a coordinate-invariant hunter-gatherer→industrial axis space, dividing all samples into 20 sequencing-depth bins of equal sample count and defining the axis separately within each bin. Of these, 17 bins contained sufficient samples of every lifestyle to be retained (15,363 samples; f**igs. S18–S19**). We found these agro-pastoralist samples and populations to be highly distinct from both hunter-gatherer and industrial samples (combined reference PERMANOVA: F=89.5, *p=*0.001, R²=0.789), lying partway along the main contrast (median fractional position=0.468, Jonckheere–Terpstra, *p<*0.001), but remaining systematically displaced off-axis (median perpendicular displacement=0.41; Wilcoxon signed-rank, *p*<0.001). This signature is consistent with a distinct agro-pastoralist component of change in the microbiome **(Fig. 2F, figs. S18–S19)**.

**Figure 2.**
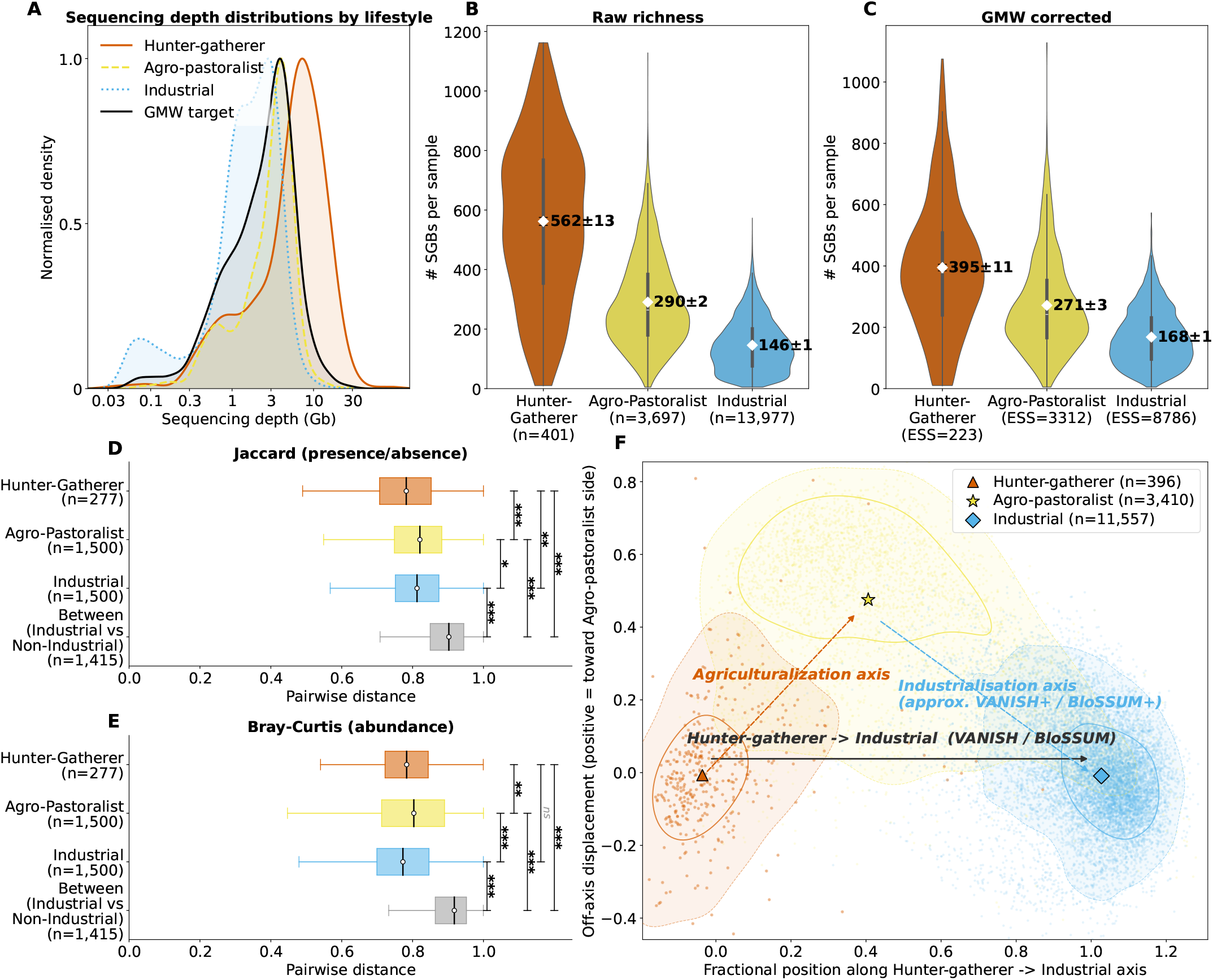
Gut microbiota diversity and composition across human lifestyles. **(A**) Hunter-gatherer and agro-pastoralist populations have generally been sampled with greater sequencing depth than industrial populations. The black line reflects the geometric-mean-target-weighted (GMW) distribution equidistant to hunter-gatherer, agro-pastoralist, and industrial sample density as a function of sequencing depth. **(B)** Due to these lifestyle-based differences in sequencing depth, uncorrected (raw) measures of microbiota richness are much higher in non-industrial than in industrial samples. **(C)** Correcting for differences in sample sequencing depth with the GMW method reduces (but does not eliminate) lifestyle-based differences in microbiota richness; after correction, industrial populations show 64% and 42% of the weighted mean richness of agro-pastoralist and hunter-gatherer populations, respectively. Overall richness counts are low because we are reporting the median of the distributions of SGB counts across samples with very different sequencing depth distributions. The richness values of the most deeply-sequenced samples (tops of each distribution) are consistent with the number of SGBs detected elsewhere (∼300-500 SGBs in industrial samples and >600 in some non-industrial samples). **(D)** Jaccard distance (presence/absence-based difference between two samples), computed on the 1:1 industrial-to-non-industrial depth-matched set (3,727 pairs, matched to within 3% of sequencing depth). For computational tractability each within-group distribution is drawn from at most 1,500 samples (hunter-gatherer *n*=277, agro-pastoralist *n*=1,500, industrial *n*=1,500), and the between-group reference from 1,000 samples per side. Distance is lower between two samples drawn from the same lifestyle than between samples drawn from industrial versus non-industrial lifestyles. Between hunter-gatherer distance is relatively low because most hunter-gatherer samples are from a single group, the Hadza of Tanzania. **(E)** Bray-Curtis distance (abundance-based difference between two samples) shows the same broad pattern. **(F)** Coordinate-invariant projection of samples onto hunter-gatherer→industrial axis space. Samples are separated into 20 depth bins with equal sample count; the 17 bins containing all three lifestyles are shown. Within each bin, each sample is projected onto a normalized axis whose endpoints are the hunter-gatherer and industrial centroids within the corresponding depth bin. The x-axis gives the sample’s fractional position along this hunter-gatherer→industrial axis (0=hunter-gatherer centroid, 1=industrial centroid), while the y-axis gives signed off-axis displacement, oriented so that positive values lie on the agro-pastoralist side. Contours show the pooled sample density for each lifestyle. Agro-pastoralist samples are intermediate along the main hunter-gatherer→industrial axis but remain consistently displaced off that axis, indicating that the shift into agro-pastoralist microbiome composition is not collinear with the direct hunter-gatherer→industrial contrast. The black arrow denotes the original hunter-gatherer→industrial contrast (approximately a VANISH/BloSSUM contrast), whereas the orange and blue arrows indicate the inferred agropastoralist and industrial components of compositional change. All panels use a ≥10% breadth coverage detection threshold.

Crucially, within each lifestyle group, individuals exhibited substantial differences in both gut microbiota richness and composition (**Fig. 2C–F**). Binning samples by sequencing depth and computing the within-bin coefficient of variation (CV: SD/median richness) revealed mean within-bin CV of 0.28 (hunter-gatherer), 0.40 (agro-pastoralist) and 0.42 (industrial) (**fig. S20**), confirming that between-sample richness differences reflect genuine biological heterogeneity and not sampling artifacts. Samples within a given lifestyle group shared only 18–22% of detected species on average (median within-group Jaccard similarity=0.22, 0.18, and 0.19 for hunter-gatherer, agro-pastoralist, and industrial samples, respectively; **Fig. 2D**). This pattern indicates that broad sampling across many individuals is required to characterize population-level microbiome composition.

## Almost all human gut bacterial taxa are present in industrial populations

For both industrial and non-industrial (i.e., hunter-gatherer and agro-pastoralist) lifestyles, we modeled population-level diversity as a function of the number of samples pooled (**Fig. 3, Supplementary Text 4**). Likewise, we considered how diversity within a lifestyle group increases as a function of the number of samples or number of populations included.

**Figure 3.**
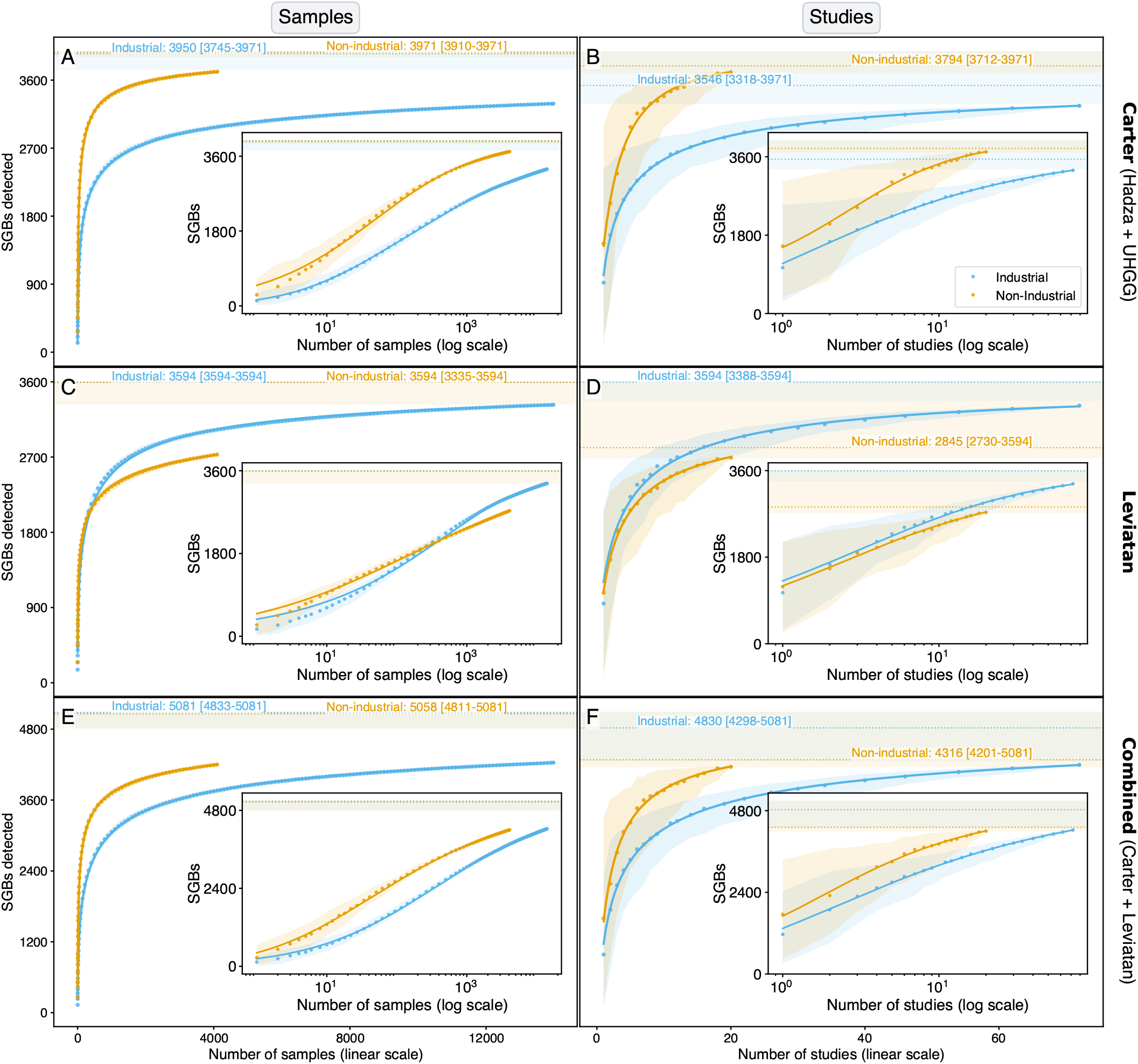
Extensive microbial extinction has not occurred in industrial populations. **Left panels**: Expected number of SGBs detected as a function of number of pooled samples. **Right panels**: Expected number of SGBs detected as a function of number of pooled studies. Outer plots are shown in linear space; inner insets are shown in log-linear space. Curves show the bootstrap median with 95% quantile ribbons; the fitted line [4-parameter Richards sigmoidal curve] is shown within the observed range. Asymptote annotations give the fitted convergence value (median) with 95% bootstrap CI. **(A, B)** Data for SGB accumulation using the Carter reference catalogue [Hadza + Unified Human Gastrointestinal Genome (UHGG)] biased towards non-industrial samples. **(C, D)** Data for SGB accumulation using the Leviatan reference catalogue biased towards industrial samples. **(E, F)** Data for SGB accumulation using the combined Carter + Leviatan reference catalogue (see **Supplementary Text 2D** for further detail). At low degrees of sample pooling both references detect more SGBs in non-industrial populations; at higher degrees of pooling, the pattern diverges according to database composition, with the Carter reference catalogue continuing to favor higher diversity in non-industrial populations and the Leviatan reference catalogue crossing over to favor higher diversity in industrial populations. At very high sample counts (*N*>100), the difference in the number of SGBs detected in the two lifestyles decreases in both the Carter and combined references. This indicates that rare microbes found in non-industrial populations and incorporated into non-industrial-biased reference catalogues are detected in industrial populations with sufficiently broad pooling of samples.

We incrementally added randomly chosen microbiome samples from a lifestyle group and updated the pooled composition, which provides a robust estimate of the change in population-level diversity as a function of the number of samples analyzed. At low levels of sample pooling, because compositional similarity between randomly chosen samples is low, each additional sample causes many species not yet present in the pooled set to appear and therefore contributes to a relatively large increase in population-level diversity. As more samples are added and the pooled diversity increases, each new sample contributes fewer previously undetected species, and thus produces diminishing increases in population-level diversity. Because the total number of microbial species within each lifestyle group is finite (*34*), continued sampling moves the population-level diversity closer to this total and approaches convergence. Empirically, we found two best-fit models for the accumulation of the number of SGBs mapped to a microbial reference metagenome as a function of the number of samples **(Supplementary Text 4)**. The best-fitting 3-parameter model was a Hill-type Michaelis-Menten sigmoidal curve 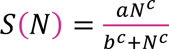 (**fig. S15**; *R*^2^∼0.993-0.999), and the best-fitting 4-parameter model was a Richards sigmoidal curve *S*(*N*) = *a*[1 + *ν* exp{−*b*(log(*n*) − *c*)}]^−1/*ν*^ **(Fig. 3A–F**; *R*^2^∼0.989-1.000). In both, *S* is the expected number of SGBs detected, *N* is the number of pooled samples, *a* is the fitted asymptote, *b* and *c* are the half-saturation and shape parameters, and *v* is the Richards asymmetry parameter. Full definitions are given **Supplementary Text 4**. These curves can be interpreted as compact empirical fits to the accumulation of SGBs found in at least one sample.

Reflecting the quality of the fit, the asymptotes of these empirical curve families were similar to the total number of SGBs in our combined (Carter + Leviatan) reference database with 5,081 distinct SGBs, suggesting that sufficiently broad sampling within a lifestyle group would recover the full diversity of previously characterized human-associated microbes. To verify that these asymptotes were not an artifact of these empirical curve families, we repeated the asymptote analysis using finite-catalogue *β*-binomial and *β*-mixture models, which explicitly model heterogeneous SGB detection probabilities and the probability that an SGB is detected in at least *k* samples **(Supplementary Text 4, fig. S16)**. We also examined the *k* = 1 curves using stricter per-sample detection thresholds requiring 30% or 50% SGB recovery (**fig. S17**). Across both stricter detection thresholds and repeated-detection thresholds of *k* = 1,2,3, and 5, the asymptotes remained close to the total number of SGBs in the combined reference catalogue.

We also examined the accumulation of the number of SGBs detected as a function of the number of studies (many of which examined different populations) included within a lifestyle group (right column in **Fig. 3** and **Fig. f15**). This validation produced similar accumulation curves and asymptotes, consistent with the total number of SGBs in the reference being recovered but with a larger uncertainty because there are far fewer studies than samples. While an individual study may not sample sufficiently broadly to recover most of the SGBs in the reference catalogue, increasing sample size by pooling studies and compositional differences between populations sharing a lifestyle rapidly capture most SGBs thereafter.

There is substantial reference bias in the characterization of rare microbial species: SGBs accumulate faster in non-industrial samples in the Carter reference and, beyond roughly 25 pooled samples, faster in industrial samples in the Leviatan reference (**Fig. 3** and **fig. S15**). However, for both industrial and non-industrial groups, we found that population-level diversity converges at or near the same total value, which is the total number of microbial taxa present in the reference catalogue. This pattern is replicated when using both the non-industrial-biased Carter reference (**Fig. 3A,B**), the industrial-biased Leviatan reference (**Fig. 3C,D**), and the combined reference (**Fig. 3E,F**). Crucially, for both non-industrial and industrial lifestyle groups, the fitted asymptotes of the fits to the median bootstrap replicates derived from pooling individual samples (**Fig. 3A,C,E**) corresponds to at least 99.5% of the total number of SGBs for each of the references. Collectively, these analyses suggest that almost all SGBs found in human guts can be recovered from both non-industrial and industrial microbiomes given sufficiently broad pooling. As a negative control, the same accumulation model compared against a chimpanzee-derived catalogue (*6*) recovered only ∼45% of SGBs in that catalogue (see **Supplementary Text 4.K** and **Table S5** in the Supplement). This number is close to previous reporting (*6*) and confirms that saturation reflects genuine microbial presence rather than permissive detection of microbes.

## Microbial species that are differentially prevalent between industrial and non-industrial populations are neither missing nor particularly rare

Next, we used the same pooled-detection framework to examine whether the taxa most differentially prevalent between industrial and non-industrial microbiomes are genuinely near an extinction threshold. Current research distinguishes VANISH taxa, which are relatively less prevalent in industrial samples, and BloSSUM taxa, which are less prevalent in non-industrial samples (*17*, *20*). In prior work, however, these lists were defined in a Hadza-versus-industrial comparison rather than the broader non-industrial-versus-industrial comparison used here. Crucially, neither VANISH nor BloSSUM taxa were especially difficult to recover with pooling (**Fig. 4A–F**, **Table S7**, **Supplementary Text 5**). We found that a median of 192 industrial samples was required to achieve a 50% probability of detecting a given VANISH taxon at ≥10% breadth coverage, compared with 485 samples for a typical SGB. Conversely, a median of only 8 non-industrial samples was required to detect a given BloSSUM taxon, compared with 95 for a typical SGB. In other words, contrary to *ipso facto* expectation, VANISH and BloSSUM taxa exhibit above-median prevalence in industrial and non-industrial populations, respectively.

**Figure 4.**
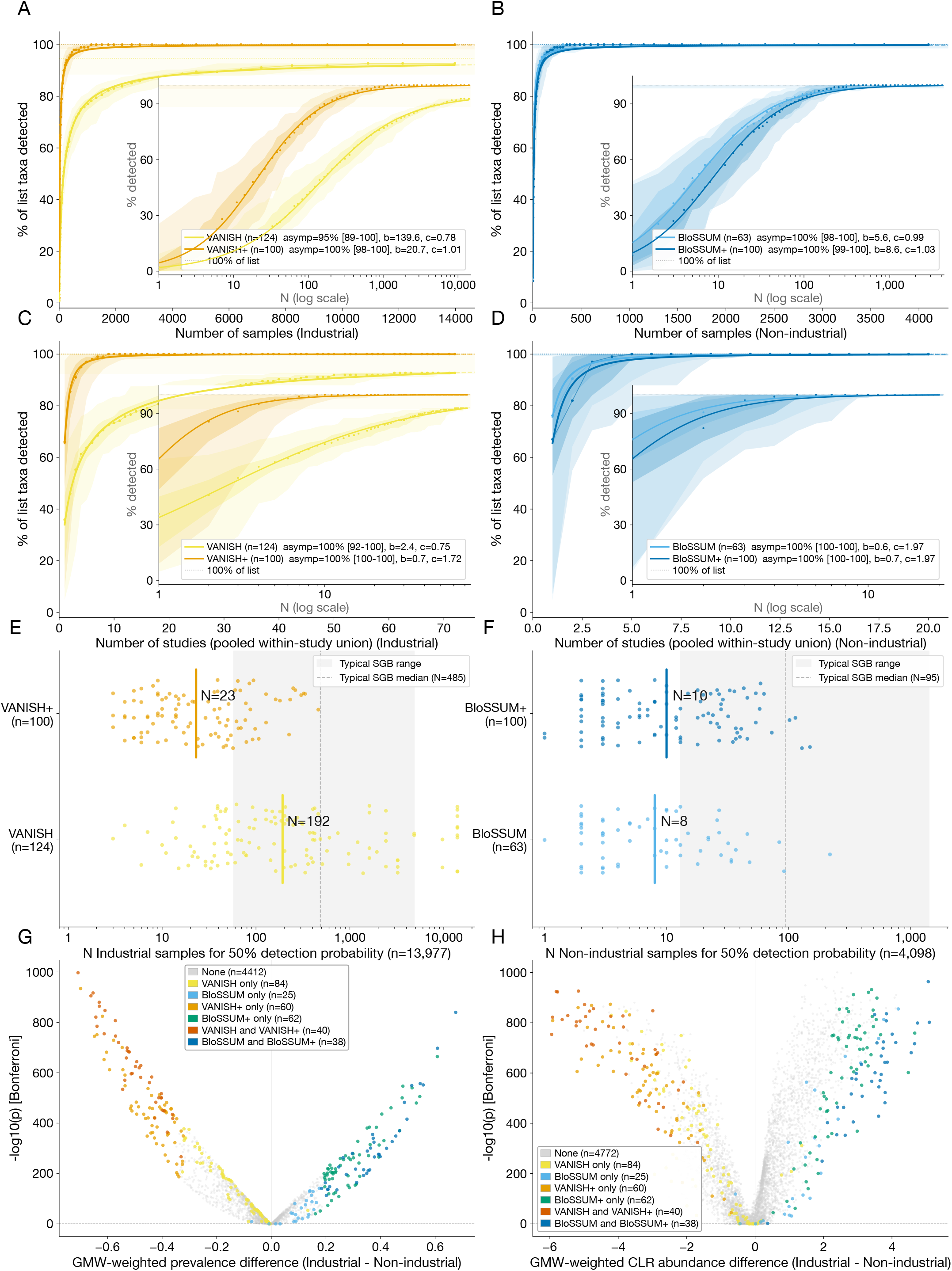
Taxa that are most differentially prevalent between industrial and non-industrial populations are not especially rare, exhibiting above-median prevalence across lifestyle groups. Accumulation of **(A,C)** VANISH taxa in industrial populations and **(B,D)** BloSSUM taxa in non-industrial populations as a function of **(A,B**) increasing number of samples pooled or **(C,D)** studies pooled. Outer plots are shown in linear space, and insets are shown in log-linear space. The Hill-type Michaelis-Menten (Hill-MM) function is fit to the data, with fitted coefficients reported in the panel legends. **(E,F)** Number of samples required for 50% detection probability of VANISH/VANISH^+^ taxa in industrial populations **(E)** or BloSSUM/BloSSUM^+^ taxa in non-industrial populations **(F),** based on the combined reference. This analysis shows that most taxa included in lists of lifestyle-discriminatory taxa (VANISH, VANISH^+^, BloSSUM, BloSSUM^+^) are not particularly rare in the populations where they are depleted. The original VANISH taxa are the rarest list, requiring a median of 192 samples to be detected in industrial populations (versus 485 for the median of all taxa in the reference), but are still more prevalent than the median-prevalence taxon in the reference. All of the BloSSUM taxa are quickly detected in non-industrial populations, requiring a median of just 8 non-industrial samples for the BloSSUM list and 10 non-industrial samples for the BloSSUM^+^ list. **(G,H)** Volcano plots showing differences in **(G)** GMW-weighted prevalence and **(H)** GMW-weighted CLR abundance between the original VANISH/BloSSUM taxa (lighter shades) and the most differentially prevalent taxa identified empirically in our analysis, i.e., the VANISH^+^/BloSSUM^+^ taxa (darker shades). Taxa labelled VANISH only are not found in the VANISH^+^ list; taxa labelled VANISH^+^ are not found in the VANISH list; those labelled VANISH and VANISH^+^ are found in both. Likewise, taxa labelled BloSSUM only are not found in the BloSSUM^+^ list; taxa labelled BloSSUM ^+^ are not found in the BloSSUM list; those labelled BloSSUM and BloSSUM ^+^ are found in both.

When pooling across the full industrial cohort defined in **Table S3** (*n*=13,977), 115 of the 124 original VANISH taxa were detected at the 10% breadth coverage threshold, while 63 of the 63 BloSSUM taxa were detected across the full non-industrial cohort (*n*=4,098). Thus, the original VANISH and BloSSUM lists are better understood as lists of taxa with strong differential prevalence across lifestyle contexts rather than lists of taxa lying near a practical extinction threshold. The nine remaining taxa are best treated as individual exceptions requiring case-by-case interpretation rather than as evidence for local extinction (**Tables S8–S9)**. Some or all of these may simply require broader sampling: the 95% interval of the fitted asymptote for the VANISH accumulation curve reaches 100% of the list (**Fig. 4A,C**).

The original VANISH and BloSSUM lists, however, do not isolate the industrial transition from pre-industrial changes in the gut microbiome because they compare the ends of the lifestyle spectrum in contrasting Hadza versus industrial microbiota. We therefore defined novel industrialization-specific lists, VANISH^+^ and BloSSUM^+^, as the 100 taxa most differentially prevalent between industrial and the collective set of non-industrial samples in the present dataset (**Data S4, Table S6, figs. S21–S22)**. These lists overlap to some extent with the original lists defined by Carter and colleagues (*20*) but are not identical (40/124 VANISH taxa overlap with VANISH^+^; 38/63 BloSSUM taxa overlap with BloSSUM^+^; **Fig. 4E–F**). Importantly, VANISH^+^ and BloSSUM^+^ are prevalence-defined lists, not rarity-defined lists: they capture the taxa most shifted in prevalence across the industrial–non-industrial contrast after correcting for sequencing-depth imbalance, not the taxa closest to extinction. The 9 taxa from the original VANISH list not detectable with pooling in industrial populations are not broadly prevalent across populations other than the Hadza. Because of this, all 100 VANISH^+^ taxa remained detectable (at 10% breadth coverage, almost all with higher breadth coverage thresholds) in industrial populations and all 100 BloSSUM^+^ taxa remained detectable in non-industrial populations (**Table S7**).

## Persistence of non-industrial-associated taxa, and prevalence of industrial-associated taxa, exhibit condition-dependent associations with health in industrial contexts

Having established that original VANISH/BloSSUM taxa persist under deep pooling and that VANISH^+^/BloSSUM^+^ capture a partially overlapping but distinct signal of industrialization, we next asked whether these focal taxa are associated with health outcomes within industrial populations. Because it is instructive to compare non-industrial samples with both all industrial samples and a subset of samples from healthy industrial individuals, we further introduce lists of taxa labelled VANISH^H+^ and BloSSUM^H+^, each capturing the 100 taxa most differentially prevalent between non-industrial and healthy industrial samples in the present dataset (**figs. S21–S22**, **Table S6**, **Data S4)**. Health and disease metadata were extracted from existing meta-analyses of cases/controls, specifically MicrobiomeDB (*35*) as well as very recent work (*36*). Each case–control comparison retained one sample per host and only studies contributing ≥5 cases and ≥5 controls were included. We analyzed four medical conditions separately: obesity, glucose intolerance, inflammatory bowel disease (IBD), and colorectal cancer(CRC)/adenoma. These were chosen because each combines prior evidence for a gut microbial contribution to disease risk or severity (*37–42*) with sufficient independent case–control metagenomes in this dataset to support study-stratified testing (at least three studies with 5 cases and 5 controls each; **Fig. 5D–E, Supplementary Text 6A–B, Tables S3**, **S10**).

**Figure 5.**
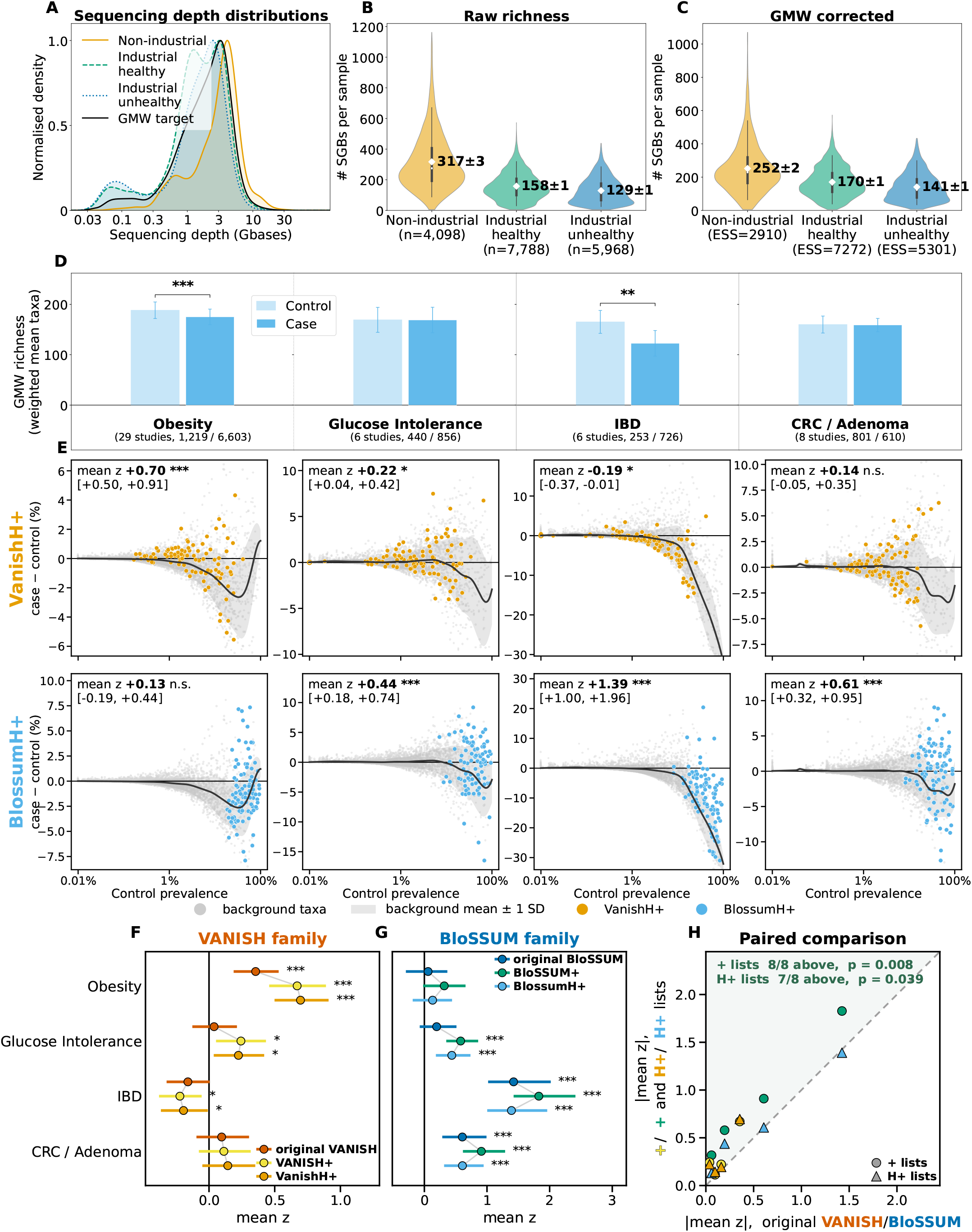
Host health status is associated with microbial richness and differential prevalence of lifestyle-discriminatory taxa. (**A**) Sequencing depth distributions for studies of non-industrial, healthy industrial, and unhealthy industrial individuals, as determined from case-control studies of metabolic and gastrointestinal diseases as well as other conditions listed in **Supplementary Text 6B**. The black line reflects the geometric-mean-target-weighted (GMW) distribution equidistant to non-industrial, healthy industrial, and unhealthy industrial sample density as a function of sequencing depth. **(B,C)** Microbiota richness expressed as the (**B**) raw number of SGBs per sample and (**C**) GMW-corrected number of SGBs per sample. GMW-corrected richness of healthy industrial individuals is intermediate between that of unhealthy industrial individuals and non-industrial (hunter-gatherer plus agro-pastoralist) individuals. **(D)** GMW-corrected richness in healthy industrial individuals (controls, pale) and unhealthy industrial individuals (cases, solid), stratified by condition; error bars are standard errors and the bracket above each pair carries the study-stratified case–control test (*p < 0.05, **p < 0.01, ***p < 0.001). The four conditions shown are those supported by at least three studies contributing both cases and controls. Significant associations with microbial richness vary by disease, after controlling for study batch effects and differences in sample sequencing depth (**Supplementary Text 6**). Each comparison retains one sample per host and only studies with at least five cases and five controls; the number of contributing studies and of cases / controls is printed beneath each condition. (**E**) For each condition, every background taxon’s study-stratified case−control prevalence difference is plotted against its control prevalence (grey), with the fitted expectation and its ±1 SD band, and the focal list taxa colored (VANISH^H+^ in the upper row, BloSSUM^H+^ in the lower). Each taxon on the two focal lists is compared with background taxa at the same control prevalence (see main text). Statistics printed in bold reflect the focal list’s mean distance from the background trend, in units of the local scatter of those background taxa (mean z), with a bootstrap 95% interval; asterisks are the two-sided bootstrap p-value, with detection at 10% genome breadth (**Supplementary Text 6G**). A positive value for this statistic indicates that the focal list taxa are more prevalent in cases than are the background taxa. VANISH^H+^ lies above it in obesity and glucose intolerance, below it in IBD, and is indistinguishable from it for CRC / adenoma. BloSSUM^H+^ lies above its background in glucose intolerance, IBD, and CRC / adenoma, but not in obesity. **Fig. S25** shows the same comparison for the original VANISH and BloSSUM taxa. **(F, G)** The same statistic for all six focal lists (VANISH, VANISH^+^, VANISH^H+^, BloSSUM, BloSSUM^+^, BloSSUM^H+^): mean z with its bootstrap 95% interval, by condition, for the three VANISH-family lists **(F)** and the three BloSSUM-family lists **(G)**, with the original VANISH or BloSSUM list at the top of each band and our + and H+ lists below it. Asterisks represent the significance of the departure of the focal list from the condition-specific background taxa (calculated as a two-sided bootstrap p-value). **(H)** For each condition, we compare |mean z| between the original VANISH/BloSSUM lists (*x*-axis) and our +/H+ lists (*y*-axis). A point above the dashed line is a condition in which our +/H+ lists outperform the original VANISH/BloSSUM lists. Because almost all points lie above the identity line, our +/H+ lists generally discriminate health from disease better than the original lists.

Additionally, we analyzed a pooled “unhealthy” group representing 26 non-communicable conditions for overall richness comparisons (**Fig. 5A–C),** comprising 13,756 industrial samples (5,968 cases, 7,788 controls) from 71 studies. After adjusting for sequencing depth using the GMW method (**Supplementary Text 3**), bacterial richness in healthy industrial individuals was intermediate between unhealthy industrial individuals and non-industrial individuals, who exhibited low and high richness, respectively (**Fig. 5A–C; figs. S23–S24**). Richness effects were heterogeneous across disease-specific comparisons, ranging from no detectable difference for glucose intolerance and CRC, to lower richness in cases of obesity (13 fewer SGBs; *p<*0.001) and a large reduction in IBD (43 fewer SGBs, *p=*0.007) (**Fig. 5D; Table S10**).

We examined whether the focal lifestyle-associated taxa (i.e., those in the VANISH, VANISH^+^, VANISH^H+^, BloSSUM, BloSSUM^+^, BloSSUM^H+^ lists) showed associations with human health status. To do so, we first began with a naïve association between a binary health predictor (case versus control) and binary change in prevalence (enrichment versus depletion) of these focal taxa (**Supplementary Text 6)**. To correct for study-level fixed effects, this approach used a GMW variant of the Cochran-Mantel-Haenszel (CMH) test that reduces the complexity of the microbial prevalence distribution to one parameter: the number of significantly enhanced or depleted focal taxa in disease cases versus controls (**Supplementary Text 6D-G**). This naïve association finds broad prevalence depletion across several disease conditions for several of the focal lists, and especially the higher-prevalence BloSSUM family lists (**Table S11).**

Because many common microbial taxa show lower prevalence in cases versus controls, depletion of specific taxa may simply reflect broad associations between health and microbial prevalence rather than associations between health and specific lifestyle-based taxa. To correct for this, among healthy individuals, we first matched each microbe on the focal lists (VANISH, VANISH^+^, VANISH^H+^, BloSSUM, BloSSUM^+^, or BloSSUM^H+^ lists) to the set of “background” taxa (those not on the focal lists) that exhibited comparable prevalence. We then scored shifts in prevalence between cases and controls for these focal and prevalence-matched background taxa, quantifying whether the focal taxa showed smaller or larger shifts than the average among background taxa (**Fig. 5E**, **Table S12**). These residuals allowed us to isolate patterns of enrichment and depletion in specific taxa from patterns due to underlying disease-associated changes in richness.

Using this prevalence-matching method, we found that harboring lifestyle-associated taxa could be associated with either health or disease, depending on the condition. In obesity, residuals derived from the distribution of prevalence-matched background taxa revealed that ‘vanishing’ taxa associated with non-industrial lifestyles (VANISH^H+^) were more likely to be associated with cases than controls, whereas ‘blossoming’ taxa differentially prevalent in industrial lifestyles (BloSSUM^H+^) were equally associated with cases and controls (**Fig. 5E**, **Table S12**). The VANISH^H+^ taxa with the largest positive residuals in obesity were *Duodenibacillus* sp900544255 (z = +3.9, q = 0.033) and *Fimenecus* sp000432435 (z = +2.8, q = 0.020), both significant after Benjamini-Hochberg correction. By contrast, among individuals with glucose intolerance, both VANISH^H+^ and BloSSUM^H+^ taxa were positively associated with cases. The largest positive residuals were carried by *Prevotella* sp003447235 (z = +3.9) and *Escherichia coli* (z = +2.2) among VANISH^H+^ taxa, and by *Bacteroides fragilis* (z = +2.3) and UMGS1623 sp934647945 (z = +2.3) among BloSSUM^H+^ taxa, though none of these four reached per-condition significance (all q ≥ 0.32). Note, the only individually significant VANISH^H+^ taxon in this condition, *Roseburia* sp900756925, was depleted in cases (z = −2.2, q = 0.008). Among individuals with CRC/adenoma, the industrial-associated BloSSUM^H+^ taxa were positively associated with cases, but the VANISH^H+^ taxa did not discriminate between cases and controls The BloSSUM^H+^ taxa most strongly associated with CRC/adenoma cases were *Bilophila wadsworthia* (z = +3.3, q = 0.033) and *Bacteroides fragilis* (z = +3.2, q = 0.033). Finally, among individuals with IBD, industrial signatures (i.e. low VANISH ^H+^ and high BloSSUM^H+^ residuals) were associated with cases. Among VANISH^H+^ taxa the most depleted in IBD cases were *Oscillibacter* sp900550685 (z = −1.0, q = 0.002) and *Prevotella copri*_M (z = −0.6, q = 0.002), while among BloSSUM^H+^ taxa the most positively associated with cases were *Phocaeicola dorei* (z = +7.3, q = 0.043), *Phocaeicola vulgatus* (z = +6.9, q = 0.043) and *Mediterraneibacter gnavus* (z = +5.6, q = 0.005). *Escherichia coli* runs against the VANISH^H+^ trend, carrying the largest positive residual on that list in IBD (z = +3.2, q = 0.002). The lists derived from our own industrialization contrast (VANISH^+^/VANISH^H+^ or BloSSUM^+^/BloSSUM^H+^) discriminated health status more effectively than the original VANISH and BloSSUM lists (**Fig. 5F-H**, **fig. S25**, **Table S12**).

Finally, to achieve greater resolution into the microbes driving the lifestyle-associated signatures of health and disease, we examined whether list-bound members of a bacterial clade more consistently discriminate health and disease compared with non-list-bound (i.e., “background”) members of the same clade (**fig. S26**, **Supplementary Text 6G.9**). Of 340 clade-based contrasts (85 clades and 4 health conditions), list-bound microbes were more discriminatory in 48 contrasts spanning 12 clades after Benjamini-Hochberg correction. For example, in obesity, the VANISH^H+^ members of *Prevotella* lay 1.1 SDs above the rest of their congeners (q = 0.003). By contrast, in IBD, VANISH^H+^ members of Oscillospiraceae lay 0.7 SDs below their non-list congeners (q = 0.001). Reflecting the relatively consistent association that we observed between BloSSUM^+^ taxa and disease across conditions, BloSSUM^+^ members of Oscillospiraceae lay above their relatives in contrasts for all four conditions (+0.8 to +1.3 SDs, q ≤ 0.013), whereas the same was true for Lachnospiraceae across three conditions (obesity, glucose intolerance, IBD, q ≤ 0.002).

Consistent with the idea that a small number of human lifestyle-associated microbes may be generally beneficial or detrimental for health (*16*, *17*), we found 8 VANISH^H+^ taxa and 21 BloSSUM^H+^ taxa that were consistently either positively or negatively associated with cases across each of our four tested conditions, and significantly associated with health or disease (at q < 0.05, after Benjamini-Hochberg correction) when data from these four conditions were pooled (**figs. S27–S30**). The direction of these signatures was strikingly one-sided on both lists. Of the 23 VANISH^H+^ taxa with a consistent pooled effect (q < 0.20), 19 were enriched in cases and 4 were depleted. Of the 33 such BloSSUM^H+^ taxa, 30 were enriched and 3 were depleted.

On the VANISH^H+^ list, the disease-associated signal was carried disproportionately by a single genus: nine of the 19 enriched taxa were *Prevotella* species, with all of them enriched in cases (mean z from +0.78 to +1.66, four at q < 0.05), including both *Prevotella copri* clades (*copri*_M, q = 0.031; *copri*_N, q = 0.079) (**fig. S27–S30**). The strongest individual effect on the list was *Escherichia coli* (mean z = +2.16, q = 0.031), enriched in cases across all four conditions. The four VANISH^H+^ taxa depleted in cases are, by contrast, organisms of significant interest as biomarkers of a healthy gut, including *Faecalibacterium prausnitzii_E* (−1.13, q = 0.162), *Roseburia* sp900756925 (−1.07, q = 0.051) and *Oscillibacter* sp900550685 (−0.63, q = 0.179) (*43–46*).

On the BloSSUM^H+^ list, the enriched taxa are, for the most part, organisms implicated in gastrointestinal or inflammatory disease, including: *Bacteroides fragilis* (+2.16), *Eggerthella lenta* (+1.94), *Bilophila wadsworthia* (+1.63) and *Bacteroides thetaiotaomicron* (+1.58) (all at q ≤ 0.02), together with two *Alistipes* species (*A. indistinctus*, +0.90; *A. ihumii*, +0.86; both q = 0.167) (*47–51*). Of the three BloSSUM^H+^ taxa depleted in cases, only *Hominimerdicola aceti* survived correction at q < 0.05 (−0.89, q = 0.037), with *Borkfalkia ceftriaxoniphila* (−0.61, q = 0.118) and *Limivivens* sp905214955 (−0.64, q = 0.175) reaching only q < 0.20.

## Discussion

Understanding how recent, dramatic transformations in human lifestyles drive gut microbiome changes is crucial for our understanding of human evolution, the etiology of microbiome-mediated diseases, and for future microbiota-directed efforts to enhance human health. Our analyses demonstrate three broad patterns that clarify the connections between industrialization, reduced gut microbiota diversity, and human health. First, individuals in industrial populations do indeed harbor lower gut microbiome diversity. However, this does not mean that industrial populations have suffered collective microbial extinctions, only that many microbes are less prevalent than in non-industrial populations. Second, the VANISH taxa are not particularly rare. Indeed, we find that they are *more* prevalent than the median taxon recovered from the industrialized gut. The reason for this is that the VANISH taxa are identified by differential prevalence – not by lifestyle- or population-wide “missingness”. Thus, it is both the case that depleted microbes may be relatively common and enriched microbes may be relatively rare. Third, we show that VANISH and BloSSUM taxa, and to an even greater extent our empirically determined sets of VANISH^+^/VANISH^H+^ and BloSSUM^+^/BloSSUM^H+^ taxa, exhibit condition-specific patterns of association with health and disease in industrial populations. Here, we discuss these three findings and their implications for human health and microbial ecology in turn.

First, our data support the idea that lifestyle-driven differences in microbial prevalence may be variably associated with health or disease in industrial contexts, depending on the condition. Using our prevalence-matching approach, we found VANISH^H+^ taxa to be positively associated with obesity rather than health. Although our current analysis is correlational and cannot establish causation, this finding resonates with prior empirical and theoretical evidence that non-industrial microbes may exacerbate metabolic disease if enriched in industrial contexts (*15*, *52*, *53*). Similarly, individuals with glucose intolerance were more likely to carry VANISH^H+^ taxa compared to controls, but were also more likely than controls to harbor BloSSUM^H+^ taxa. With respect to CRC/adenoma, we observed a positive association between BloSSUM^H+^ taxa and cases, but did not detect a consistent generalized signature between VANISH^H+^ taxa and health or disease. Across all three conditions, we found a positive association between disease and 9 *Prevotella* species on the VANISH^H+^ list. These *Prevotella* taxa were more strongly associated with disease than *Prevotella* congeners not on the VANISH^H+^ list, underscoring the power of lifestyle-associated taxa to discriminate health and disease states. Critically, this pattern is consistent with prior findings that some members of this non-industrial-associated genus may promote disease in industrial individuals (*46*). Finally, with respect to IBD, VANISH^H+^ taxa were more likely to be depleted among cases, while BloSSUM^H+^ taxa were strongly positively associated with cases. This signature is consistent with the idea that industrialization has exerted detrimental health effects in certain disease states (*54*). Overall, despite substantial heterogeneity in these signatures across the four conditions, enrichments of BloSSUM^H+^ taxa were more strongly associated with disease across health conditions than were depletions of VANISH^H+^ taxa. Taken together, this pattern suggests that pathology may be more closely linked to the acquisition of new microbes rather than the loss of old ones.

The strength of such inferences will improve as more uniformly designed case-control metagenomic studies become available. Notably, the absence of case-control studies coupled to metagenomic data in non-industrial populations meant that we could not examine how the prevalence of lifestyle-associated taxa associated with health among non-industrial populations. Performing this reverse analysis would help distinguish whether health is associated with a restriction of microbiota membership to taxa associated with non-industrial lifestyles or simply by increasing taxonomic diversity *per se*. Additionally, the small number of case/control studies reporting whether participants were treated versus treatment-naïve restricts our ability to discriminate the extent to which differential patterns of VANISH/VANISH^+^/VANISH^H+^ and BloSSUM/BloSSUM^+^/BloSSUM^H+^ prevalence reflect microbial associations with disease versus treatment of disease. Finally, our analysis detects only correlational patterns, encouraging future experimental investigation of the causal role of gut microbes in these lifestyle-health associations.

These results advance our understanding of bacterial extinction in the microbiome. One possibility proposed in the field is that industrialization has driven certain evolutionarily important microbial taxa to local extinction (*8*, *16*, *17*, *20*, *38*). The present work demonstrates that microbes feared to be extinct in industrial populations instead persist at low prevalences at the population level, and have likely gone undetected in prior analyses owing to limited depth and breadth of sequencing. This conclusion applies only to SGBs present in the tested reference catalogues, which recover a smaller share of non-industrial than of industrial sequencing effort (**fig. S1**), and does not rule out the potential loss of unrepresented taxa, strain-level diversity, or indeed of microbial functions. Although our analysis suggests a possible role of depleted taxa in health and disease in industrial populations, it remains unclear whether taxa present at such low prevalence retain functional significance for the host or instead represent transient colonizers that lack stable ecological niches in the industrial gut environment. Further work will be necessary to delineate the functions and physiological effects of these depleted taxa. Relevant approaches might include, for instance, spike-in experiments and studies of synthetic communities of bacterial isolates in which the presence and abundance of these taxa can be manipulated. Such efforts will be important for discriminating between these alternatives.

The persistence of non-industrial lifestyle-associated taxa in industrial populations suggests that, despite widespread influence of ecology on the gut microbiome, at least some hosts might be able to retain critical microbes in novel ecologies, thereby benefiting from some of the ecosystem services that those microbes provide. For example, seasonal variation in the Hadza drives some microbes to below-detection levels, only for these taxa to rebound when ecological conditions favorable to their expansion return (*24*). The capacity of the human gut microbiome to retain access to ecosystem services by maintaining a persistent reservoir of low-abundance or low-prevalence microbes at the population level has been proposed to be an important force in shaping both individual health outcomes and the benefits and costs of social interaction (*55*, *56*). Future studies that isolate the genetic, physiological, and social conditions under which low-abundance or low-prevalence microbes are acquired and transmitted will help us better understand the functions of this microbial reservoir and the mechanisms through which taxonomic persistence is maintained.

Finally, our findings highlight two possibilities for microbiota-directed therapies for diseases associated with industrial lifestyles. First, under a model of widespread microbial extinctions (*15*, *17*), restoration ultimately depends on reacquiring missing microbes, which may include ethically challenging solutions such as sourcing microbes from vulnerable hunter-gatherer or other non-industrial populations (*8*, *21*). In contrast, our finding of the persistence of almost all human-associated taxa in both industrial and non-industrial contexts suggests that restoration or therapeutic efforts might instead focus on extracting and enriching still-existing bacterial diversity from within the populations that stand to benefit. Second, our finding that carriage of VANISH/VANISH^+^/VANISH^H+^ taxa can be associated with disease as well as health in industrial populations highlights the possibility that industrial microbiomes may be a health asset in industrial contexts. Under this model, a therapeutic approach might entail reducing deviations from a healthy industrial microbiome rather than attempting to reconstruct a non-industrial profile. Ultimately, any microbiome-directed therapy will necessitate understanding the ecological context of both the microbiome and the host.

## Data, code, and materials availability

All sequencing data analyzed in this study are publicly available. Raw reads are deposited in the NCBI Sequence Read Archive (SRA; https://www.ncbi.nlm.nih.gov/sra) and the European Nucleotide Archive (ENA; https://www.ebi.ac.uk/ena); the accession for every sample is listed in **Data S2**. The analysis reads eight per-sample breadth and depth matrices and the curated per-sample metadata table archived at Zenodo under DOI 10.5281/zenodo.23437518 (*57*). The download_data.sh script in the analysis repository retrieves them and verifies every checksum before use. The figures, result tables, per-figure source data and interactive cohort visualizations are archived in a companion Zenodo record, DOI 10.5281/zenodo.23444503 (*58*). The source values behind every figure panel are provided as **Data S1**, and the curated metadata, reference catalogues and taxon lists as **Data S2–S4**; all four are deposited as individual files (Data_S1_figure_data.xlsx, Data_S2_metadata_M1.xlsx, Data_S3_metadata_M2.xlsx and Data_S4_metadata_M3.xlsx) in the companion Zenodo record (*58*). Both analysis stages are in version-controlled repositories: preprocessing, from raw reads to coverage matrices, at https://github.com/neilibata/Bacterial_persistence_preprocessing, and the analysis that consumes those matrices at https://github.com/neilibata/Bacterial_persistence_analysis_pipeline. The exact state used here is tagged v1.0-as-published and v1.0.0 respectively. Full descriptions of both workflows are provided in **Figure S31** and **Supplementary Text 2**. All other data are in the manuscript or the supplementary materials.

## List of Supplementary Materials

## Materials and Methods

Figs. S1–S31

Supplementary Text

Tables S1–S13

References (1–115)

Data S1–S4

## Acknowledgments

We thank Cary Allen-Blevins, Joseph Henrich, Daniel Lieberman, Grace Rubin, Emily Venable, Laura Schell for feedback and advice. **Funding**: A.S., E.B., and L.R.P report no relevant funding. N.G.O.I acknowledges funding from the William F. Milton Fund and the Dean’s Competitive Fund for Promising Scholarship of Harvard University. A.H.M acknowledges funding from the National Institutes of Health (R35 GM138284 and R01 DK139214). R.N.C. acknowledges funding from the National Institute of General Medical Sciences (1R35GM160067), and the William F. Milton Fund and the Dean’s Competitive Fund for Promising Scholarship of Harvard University. The content of this article is solely the responsibility of the authors and does not necessarily represent the official views of the National Institutes of Health. **Author contributions:** Conceptualization: N.G.O.I., A.S., A.H.M., and R.N.C.; Methodology: N.G.O.I., E.B., L.R.P., A.H.M., and R.N.C.; Software: N.G.O.I.; Formal analysis: N.G.O.I., A.H.M., and R.N.C.; Data curation: N.G.O.I.; Writing – original draft: N.G.O.I., A.S., R.N.C.; Writing – review and editing: all co-authors reviewed and edited the manuscript. **Competing interests:** The authors declare that they have no competing interests.

