## Supplementary material for "Persistence of Non-industrial Gut Microbiota Discriminates Health and Disease States in Industrial Populations": Materials and Methods, Supp figures, tables, text

### Supplementary Materials

#### Materials and Methods

##### *Dataset assembly and analytic cohorts*

We assembled a large metagenomic dataset containing public-domain human gut shotgun metagenomes deposited in the NCBI Sequence Read Archive (SRA) or European Nucleotide Archive (ENA), seeded from existing curated resources and then extended with manual literature review (**Supplementary Text 1A**). Studies were eligible for inclusion if they profiled the human gut microbiome using shotgun metagenomics with sufficient metadata for lifestyle and/or health annotation. Each retained sample was required to have ≥100 million raw bases prior to preprocessing and ≥30 million bacterial bases mapped at MAPQ >= 30 after removal of host or PhiX reads. We worked with bases rather than reads throughout to avoid read-length ascertainment bias of microbial species (see also QC in **Fig. S14**). Samples from any bodily site other than the gut and samples from children younger than three years were excluded (**Supplementary Text 1B**).

Sequencing runs were consolidated at the BioSample level, so that each sample is one BioSample. Repeat samples from the same host were retained in the lifestyle analyses (with a one-sample-per-host sensitivity analysis for richness), but health analyses were restricted to one sample per host; technical replicates were used only in robustness checks (**Figs. S2–S3**, **Supplementary Text 1C**). This pipeline yielded a final analytic cohort of $N = 18{,}075$ samples from 14,624 hosts sampled in 83 studies across 42 countries. These comprised an industrial lifestyle group ($n = 13{,}977$) and a non-industrial lifestyle group ($n = 4{,}098$), with the non-industrial lifestyle group comprised of samples from hunter-gatherers ($n = 401$) and agro-pastoralists ($n = 3{,}697$). Canonical and figure-specific cohort definitions are given in **Supplementary Text 1D–1E** and **Tables S1–S3**. Metadata from individual studies are listed in **Data S2**.

The curated per-sample metadata table is archived at Zenodo under DOI 10.5281/zenodo.23437518 (*1*) and is provided as **Data S2**. Details of the curation process, including selection criteria and quality filters, are given in **Supplementary Text 1**.

##### *Read processing, reference catalogues, microbial detection, and relevant software tools*

Because the central question of taxonomic extinction concerns the low-abundance tail of the microbiome, we used reference-based alignment rather than *de novo* assembly, which preferentially recovers abundant taxa and inflates apparent "missingness" in samples with shallow sequencing (**Supplementary Text 2A**). All metagenomes were reprocessed from raw reads through a single Snakemake workflow (*2*): downloading from SRA/ENA, quality and adapter trimming with Trim Galore, filtering against PhiX and the human hg38 genome, repairing reads with bbtools repair.sh, and aligning to the reference catalogues (*3*). Coverage was summarized with samtools coverage (*4*) and aggregated from contig to species genome bin (SGB) (*5*). To suppress cross-genome multimapping, reads with MAPQ < 30 were discarded (**Supplementary Text 2B**).

To address reference ascertainment bias, all samples were aligned independently to two complementary catalogues with opposing discovery biases: the Carter catalogue [Hadza + Unified Human Gastrointestinal Genome (UHGG) (*6*)], enriched for non-industrial discovery (*7*), and the Leviatan catalogue, enriched for industrial discovery (*8*) (**Supplementary Text 2C**, **Table S4**, **Data S3**). The most credible biological signals are those robust to both biases. Furthermore, we constructed a combined non-redundant reference by sketching all SGBs with sourmash (*9*), computing symmetric containment-derived ANI, and clustering at ≥95% ANI by complete linkage (*10*), retaining the higher-mapping representative per shared cluster (5,081 combined SGBs; **Supplementary Text 2D**). A taxon was scored as detected when sequenced reads mapped to ≥10% of its reference genome length. This threshold excludes spurious detection from common plasmids or contaminant DNA, and key results were confirmed at 30% and 50% thresholds (**Figs. S15–S17**, **Supplementary Text 4**).

Preprocessing was automated with Snakemake, which also resubmitted failed samples at higher memory and time allocations. The preprocessing workflow is available at https://github.com/neilibata/Bacterial_persistence_preprocessing (tag v1.0-as-published) and the analysis code that consumes its output at https://github.com/neilibata/Bacterial_persistence_analysis_pipeline (tag v1.0.0). Both workflows are described in full in **Figure S31** and **Supplementary Text 2**.

***Correction for sequencing-depth bias***

Sequencing depth was strongly associated with recoverable richness (Spearman's $\rho = 0.682$, $p < 10^{-300}$, $N = 18{,}075$), and sequencing depth differed systematically across lifestyles (**Fig. S9**). Thus, raw richness comparisons conflated biology with sequencing effort. Rather than rarefying the data, which discards data and is ill-suited to probing the rare tail of the taxonomic abundance and prevalence distributions, we instead applied geometric-mean-target weighting (GMW). GMW is an inverse-probability-weighting (*11*) variant that reweights samples toward a target depth distribution defined as the geometric mean of the groups' kernel-density depth distributions, equidistant in Kullback–Leibler divergence from each group and therefore favoring neither (*12*) (**Supplementary Text 3**). GMW retains all samples and was used to compare sampled distributions of SGB counts per sample for every between-group richness, composition, and health analysis (**Figs. 2A–C**, **5A–C**, **Figs. S5–S13**). Crucially, this method is not useful to ascertain within-group richness/composition differences, as it only accounts for systematic between-group differences in sequencing depth distributions and does not remove within-group biases in sequencing depth.

***Per-sample richness and compositional analyses***

We used bootstrap permutation (*13*) to compare depth-corrected per-sample richness across lifestyles (**Fig. 2C**). Because that bootstrap resamples samples rather than hosts, and 3,451 samples are repeat samples of a host already in the cohort, we repeated the comparison resampling hosts with their samples kept together, and again retaining one sample per host (**Table S13, Supplementary Text 7C**). Compositional structure was assessed using Jaccard (presence/absence) and Bray–Curtis (abundance) dissimilarities on 1:1 depth-matched samples (**Fig. 2D–E**). To test whether agro-pastoral microbiomes were intermediate in composition between hunter-gatherer and industrial states, samples were binned into 17 equal-count depth bins and projected onto a coordinate-invariant hunter-gatherer→industrial axis. Position along, and perpendicular displacement from, this axis were evaluated by PERMANOVA, Jonckheere–Terpstra, and Wilcoxon signed-rank tests (**Fig. 2F**, **Figs. S18–S20**).

***Population-level accumulation and persistence modelling***

Population-level diversity was modelled as a finite-catalogue discovery problem: the expected number of referenced SGBs detected as samples or studies were pooled, with the asymptote (convergence value) as the inferential target (**Supplementary Text 4**). Samples (or studies) were incrementally pooled in bootstrap replicates; first-detection ($k = 1$) accumulation curves were fit with three-parameter Hill-type Michaelis–Menten and four-parameter Richards sigmoidal functions (**Figs. 3**, **4A–D, Fig. S15**). Uncertainty in the convergence value was obtained by likelihood profiling over bootstrap replicates. We stress-tested the robustness of these results by repeating the analysis with finite-catalogue beta-binomial and beta-mixture models for detection in at least $k = 1,2,3,$ and 5 samples, and at stricter 30% and 50% detection thresholds (**Figs. S16–S17**, **Supplementary Text 4**). Accumulation was computed separately for the Carter, Leviatan, and combined reference catalogues, and, as a negative control, for a chimpanzee-derived catalogue over the same reads (**Supplementary Text 4K, Table S5**).

##### *Lifestyle-discriminatory taxon lists*

We evaluated published lists of VANISH (depleted in industrial populations) and BloSSUM (enriched in industrial populations) taxa (*7*). We also used our empirical data to define industrialization-specific VANISH$^{+}$ and BloSSUM$^{+}$ lists, representing the 200 taxa (100 per list) most differentially prevalent between industrial and non-industrial samples, and health-targeted VANISH$^{H+}$ and BloSSUM$^{H+}$ lists, each representing the 100 taxa most differentially prevalent between healthy-industrial and non-industrial samples, in both cases after GMW depth correction (**Figs. S21–S22**, **Supplementary Text 5**, **Table S6**, **Data S4**). The number of samples required to reach a 50% detection probability was computed for each focal taxon and compared with the catalogue median (**Fig. 4E–F**, **Tables S7–S9**). Rationale for the multiple list families and guidance on which list is primary for which question are given in **Supplementary Text 5A–5I**.

##### *Within-industrial health-association analyses*

Due to a paucity of case–control studies for non-industrial populations, microbiome–health associations were tested within industrial cohorts only. We tested for associations between microbial presence/absence and binary "unhealthy" (case) / "healthy" (control) designation across four metabolic and gastrointestinal conditions supported by at least three independent studies [obesity, glucose intolerance, inflammatory bowel disease (IBD), and colorectal cancer (CRC)], and a pooled "unhealthy" group representing cases across the following 26 recorded conditions: obesity (BMI ≥ 30 or study-designated), type 1 diabetes, type 2 diabetes, impaired glucose tolerance, metabolic syndrome, metabolic dysfunction-associated steatotic liver disease (MASLD), hypertension, atherosclerotic cardiovascular disease, colorectal cancer, colorectal adenoma, melanoma, inflammatory bowel disease (Crohn's disease, ulcerative colitis, or unspecified), treatment-associated colitis, *Clostridioides difficile* infection, Shiga toxin-producing Escherichia coli infection, acute diarrhoea, soil-transmitted helminth infection, intestinal colonization by multidrug-resistant bacteria, liver cirrhosis (with hepatitis, ascites, schistosomiasis, or Wilson's disease), multiple sclerosis, Parkinson's disease, rheumatoid arthritis, ankylosing spondylitis, Behçet's disease, myalgic encephalomyelitis/chronic fatigue syndrome, and schizophrenia. Each study contributed an independent stratum, each host contributed one sample, and only strata with ≥5 cases and ≥5 controls were retained (**Supplementary Text 6B**, **Tables S3**, **S10**). The primary taxon-level test was a geometric-mean-target weighted Cochran–Mantel–Haenszel test (GMW-CMH) that combines study stratification with within-stratum depth equalization (**Fig. S24**, **Supplementary Text 6D**, **6G**) (*14*, *15*).

Because baseline prevalence constrains how much a taxon can shift between cases and controls, each focal list (VANISH, BloSSUM, VANISH$^{+}$, BloSSUM$^{+}$, VANISH$^{H+}$, BloSSUM$^{H+}$) was compared not with the raw background pool but with a fitted expectation for taxa of the same prevalence. For each condition we regressed the study-stratified case-control prevalence difference of every non-list taxon on its within-study control prevalence, and scored each list taxon by its distance from that fitted curve in units of the local spread. The list-level statistic is the mean of those standardised distances, with a confidence interval from a bootstrap that refits the curve on resampled background taxa (**Fig. 5E**, **Figs. S25–S30**, **Supplementary Text 6E–6F**). Raw and matched case–control results across all list families, together with driver-taxon analyses, are reported in **Tables S10–S12** and **Figs. S25–S30**.

***Robustness analyses and limitations***

The principal robustness analyses, and the limits of what the data can support, are overviewed and examined further in **Supplementary Text 7**. These comprise dependence on the reference catalogue and the resulting bounds on any inference about taxa it does not contain (**Supplementary Text 7A**); read-length and other sequencing-design effects, which motivated our use of mapped bases rather than reads as the measure of sequencing effort (**Supplementary Text 7B, Fig. S14**); the independence of samples, given that 23 of the 83 studies sampled some participants more than once (**Supplementary Text 7C, Table S13, Fig. S13**); and sensitivity to the genome-breadth detection threshold, which key results were reproduced at 30% and 50% as well as 10% (**Supplementary Text 7D**). **Supplementary Text 7E–7G** sets out three interpretive distinctions that bear on how these results should be read: detectability of a taxon at the population level does not imply that it is functionally important (**7E**), prevalence is not abundance (**7F**), and our central claim concerns attenuation of prevalence rather than extinction (**7G**).

### Figures S1–S31

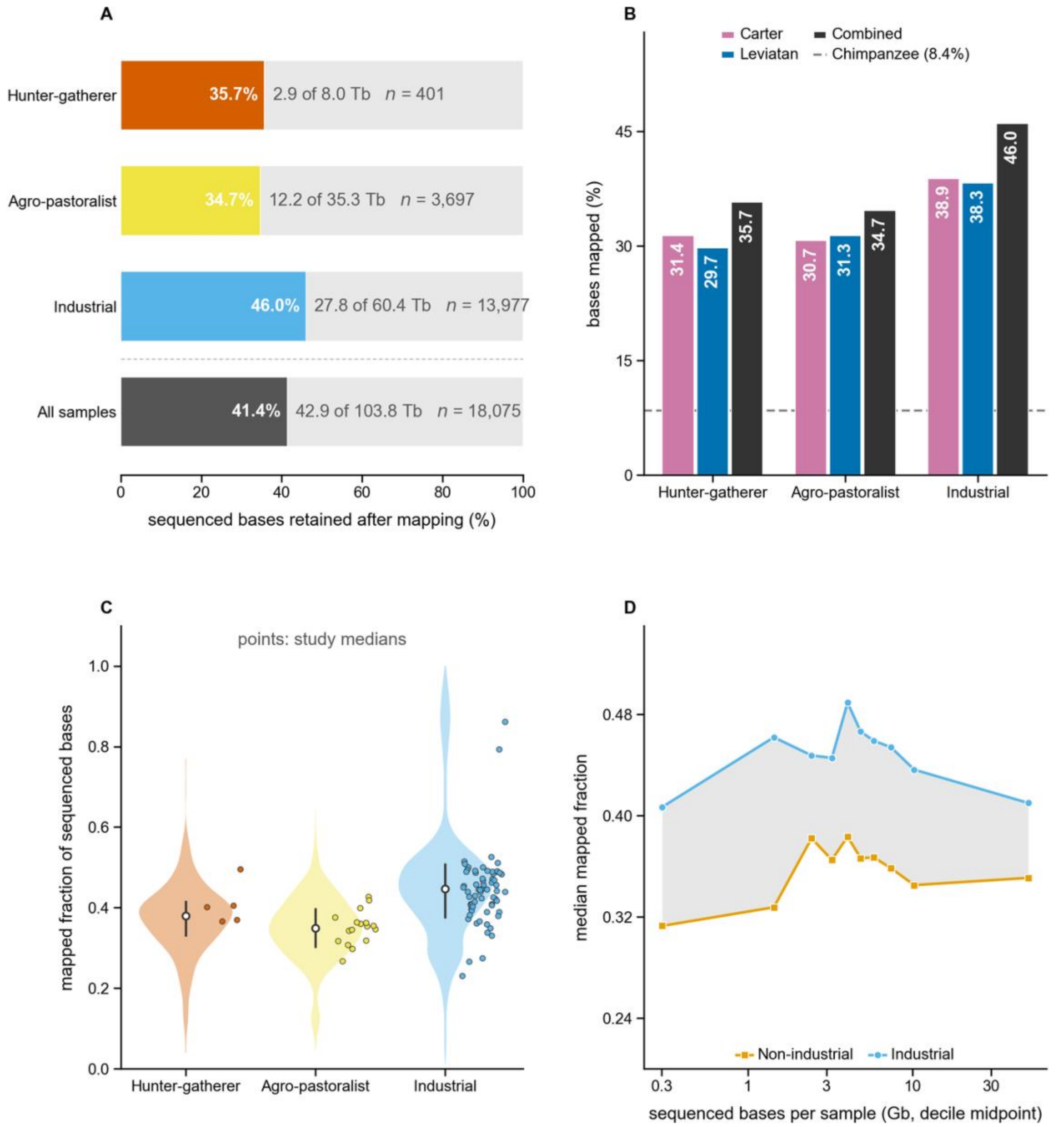

**Figure S1. Read accounting and reference recovery across the dataset.** Read accounting and reference recovery across the dataset. **(A)** For each lifestyle, the share of sequenced bases retained after trimming, host-read removal and alignment at MAPQ ≥30, against the combined reference; absolute totals and sample counts are printed beside each bar, and the pooled figure quoted in the main text is the bottom row. **(B)** The same quantity for each reference catalogue separately (Carter, 3,971 SGBs, 8.92 Gb; Leviatan, 3,594 SGBs, 8.75 Gb; combined, 5,081 SGBs, 11.62 Gb). The dashed line is the chimpanzee catalogue over the 3,799 industrial samples processed against it that carry an archived sequencing total (Supplementary Text 4K), the floor a non-human catalogue reaches on these reads. The combined reference recovers more than either single catalogue in every lifestyle, and the Carter catalogue exceeds the Leviatan catalogue on hunter-gatherer samples (31.4% against 29.7%) while the two are level on industrial samples, the asymmetry in ascertainment their provenance predicts. **(C)** The per-sample distribution behind (A). The white marker is the group median and the bar its interquartile range; each point beside a violin is one study's median, for the 89 studies contributing at least ten samples. The difference is a property of studies rather than of samples: the median across 67 industrial studies is 0.444 against 0.361 across 22 non-industrial studies (Mann-Whitney on study medians, $p = 4 \times 10^{-6}$). **(D)** The same contrast within deciles of sequencing depth, equal in sample count. Industrial samples map a higher median fraction in all ten deciles, with a median gap of 9.3 percentage points, and mapped fraction is close to flat in depth within each group (Spearman $\rho = +0.08$ industrial, $-0.01$ non-industrial), so the difference is not the depth imbalance of **Fig. S4** in another form. Mapped bases are Σ(mean depth × genome length) per sample and sequenced bases are the archived totals, so the unmapped remainder in (A) comprises trimmed, host-derived and unalignable reads together and is not attributable to catalogue coverage alone. The comparison in (B) holds reads, trimming and host removal fixed, and there the asymmetry is the catalogue's: non-industrial samples recover a smaller share against every human catalogue tested, and because an uncatalogued taxon contributes no mapped bases, the richness comparisons of Fig. 2 are made against a reference ascertainment bias that runs counter to them.

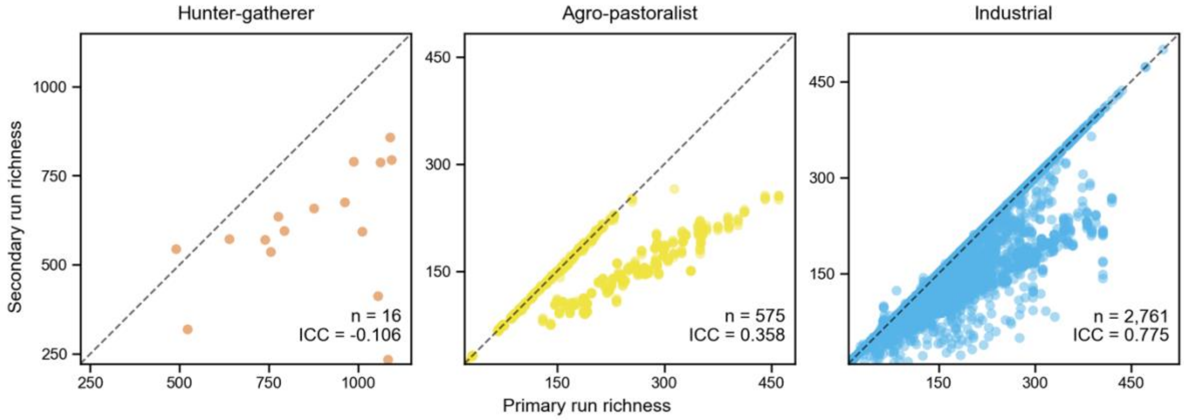

**Figure S2. Technical replicates confirm the high reproducibility of richness estimates, particularly in industrial samples.** Scatter plots comparing SGB richness (≥10% breadth coverage) between primary and secondary sequencing runs from the same biosample (technical replicates), for Hunter-gatherer (orange, $n$=11), Agro-pastoralist (yellow, $n$=574), and Industrial (blue, $n$=2,761) samples. The dashed diagonal represents perfect concordance. Intraclass correlation coefficients (ICC) are shown per group. Both groups appear to show instances of perfect sample replicates along the diagonal, and a more meaningful line showing secondary runs with lower richness (because we define the primary run as the run with the highest sequencing depth) that is strongly correlated to that of the primary run.

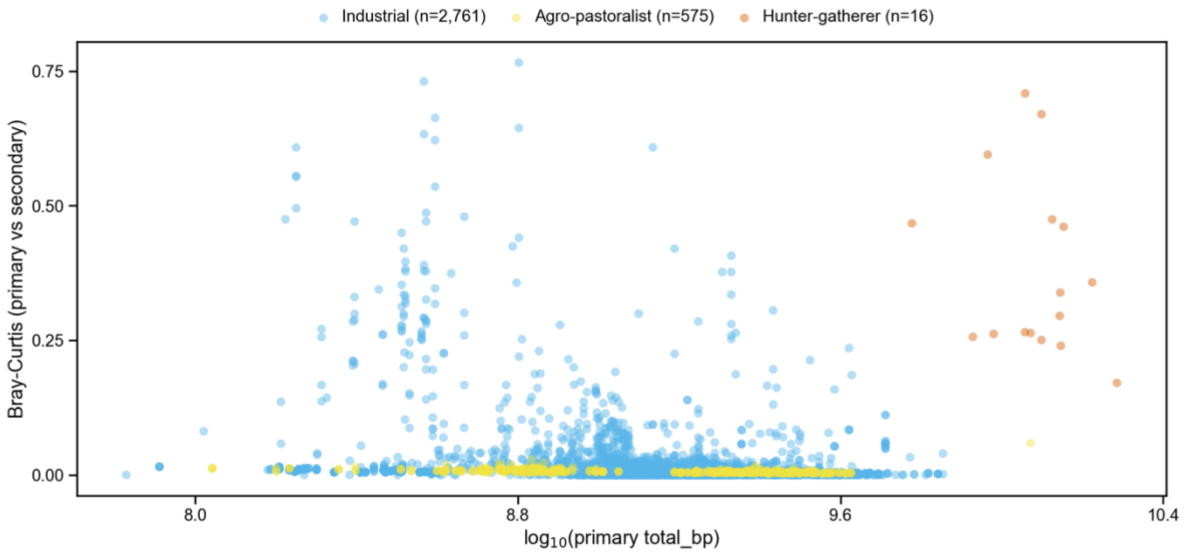

**Figure S3. Within-sample Bray-Curtis dissimilarity between technical replicates is low and largely independent of sequencing depth.** Each point represents a pair of technical replicates (primary vs. secondary run, where primary runs were defined to have the higher sequencing depth) from the same individual, plotting the Bray-Curtis dissimilarity between the two runs (y-axis; 0 = identical composition, 1 = completely different) against the primary run's sequencing depth ($\log_{10}$ total mapped bp; x-axis), for Hunter-gatherer (orange, *n*=11), Agro-pastoralist (yellow, *n*=574) and Industrial (blue, *n*=2,761) samples. The large majority of pairs cluster at near-zero dissimilarity, indicating that the compositional profiles of technical replicates are highly reproducible. A subset of pairs – particularly at intermediate sequencing depths (~$10^{8.5}$–$10^9$ bp) – show elevated dissimilarity (up to ~0.10).

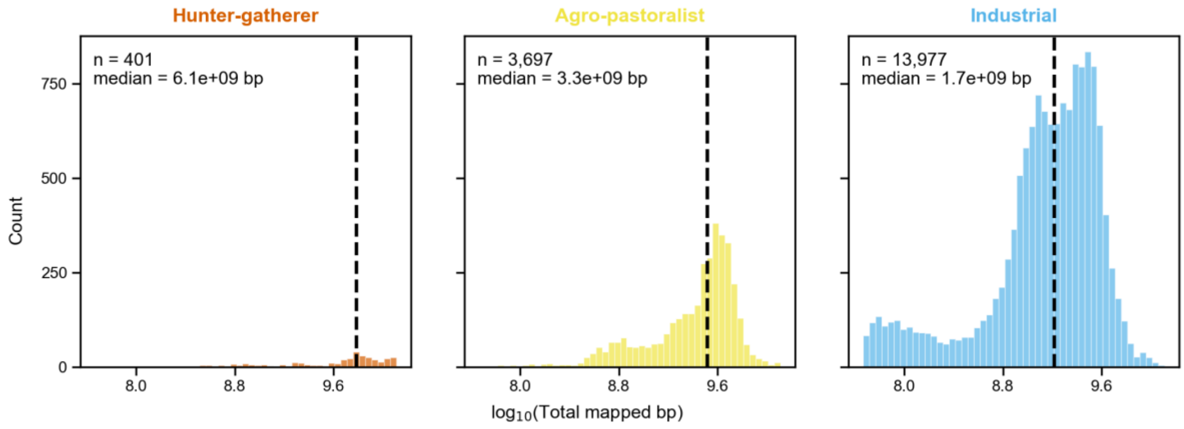

**Figure S4. Sequencing depth distributions by lifestyle group, before depth-matching.** Histograms of total bacterial bases mapped per sample ($\log_{10}$ scale) for the three lifestyle groups present in the main dataset prior to any depth-matching: Hunter-gatherer (orange, *n*=401), Agro-pastoralist (yellow, *n*=3,697), and Industrial (blue, *n*=13,977). Dashed lines indicate group medians. Industrial samples are sequenced at substantially lower depth (median $1.7\times10^9$ bp) than Agro-pastoralist (median $3.3\times10^9$ bp) or Hunter-gatherer samples (median $6.1\times10^9$ bp), reflecting systematic differences in sequencing effort across studies rather than biological differences. This confound motivates the depth-correction methods applied throughout the analysis.

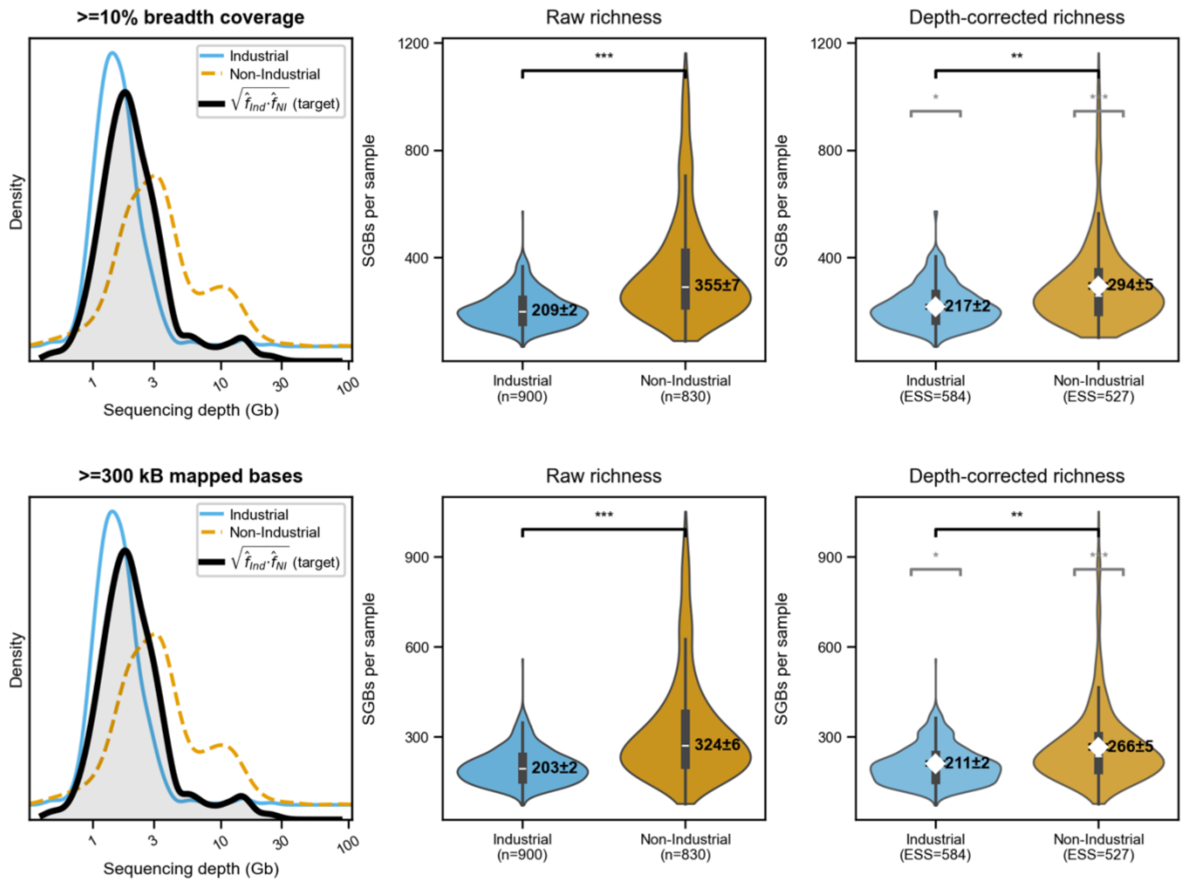

**Figure S5. Focus on samples from the Carter study: Geometric-mean-target weighting (GMW) corrects for sequencing depth bias in richness comparisons.** Each row corresponds to a different taxon detection threshold (top: ≥10% breadth coverage; bottom: ≥200 kb mapped bases). Left panels show the sequencing depth density distributions for Industrial (blue) and Non-Industrial (agro-pastoralist + hunter-gatherer; gold) samples, with the GMW target distribution (black curve) overlaid. The GMW method reweights each sample in proportion to how well its depth matches this shared target, so that neither group is systematically favored. Middle panels show violin plots of raw (uncorrected) per-sample SGB richness; right panels show the GMW-corrected richness, with weighted means (white dots ± weighted SEM) and 95% bootstrap confidence intervals. The pattern here is similar to what we show in **fig. S9** below, albeit with a smaller effect size. GMW reweighting significantly reduces the apparent richness of non-industrial samples (by ~42 taxa).

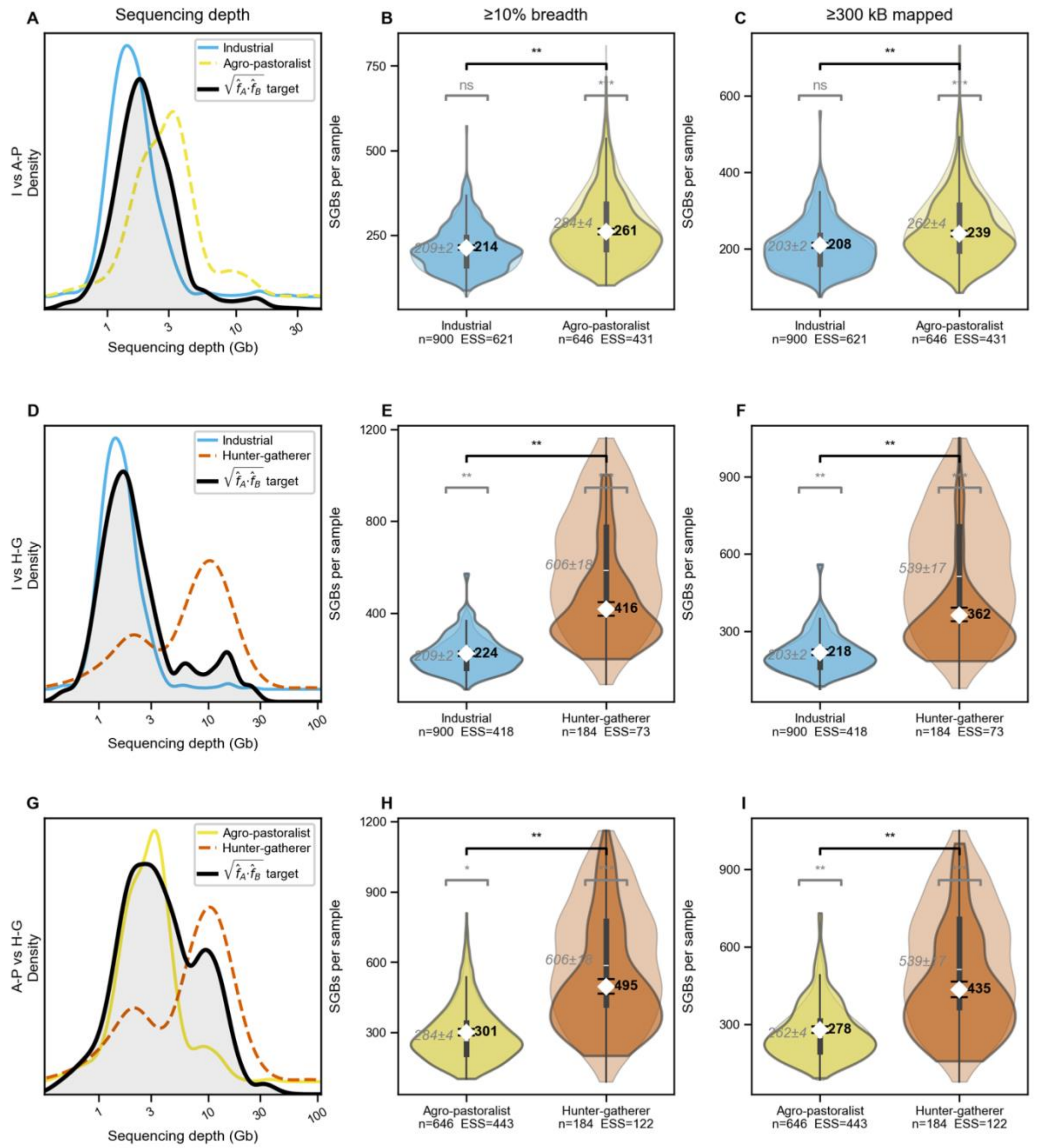

**Figure S6. Focus on samples from the Carter study: Geometric-mean-target weighting (GMW) corrects for sequencing depth bias in richness comparisons between lifestyles.** A 3-way comparison of GMW-corrected richness between Industrial, Agro-pastoralist, and Hunter-Gatherer samples in the Carter et al. sample set finds significantly smaller richness differences between Hunter-Gatherer and the other groups than without correction, and does not have sufficient power to detect any differences in richness between Industrial and Agro-pastoralist samples. The top row panels show the sequencing depth density distributions for Industrial (blue) and Non-Industrial (agro-pastoralist + hunter-gatherer; gold) samples, with the smoothed GMW target distribution (black curve) overlaid. The middle and bottom row show the results of applying this method to two different richness metrics: a coverage threshold where finding 10% or more of the SGB reference genome counts as a detection, and a flat threshold where finding 200kb or more of the SGB reference genome counts as a detection.

The shadowed distribution is the raw distribution. Reweighting it substantially changes it for hunter-gatherers and somewhat shifts it for agro-pastoralist populations.

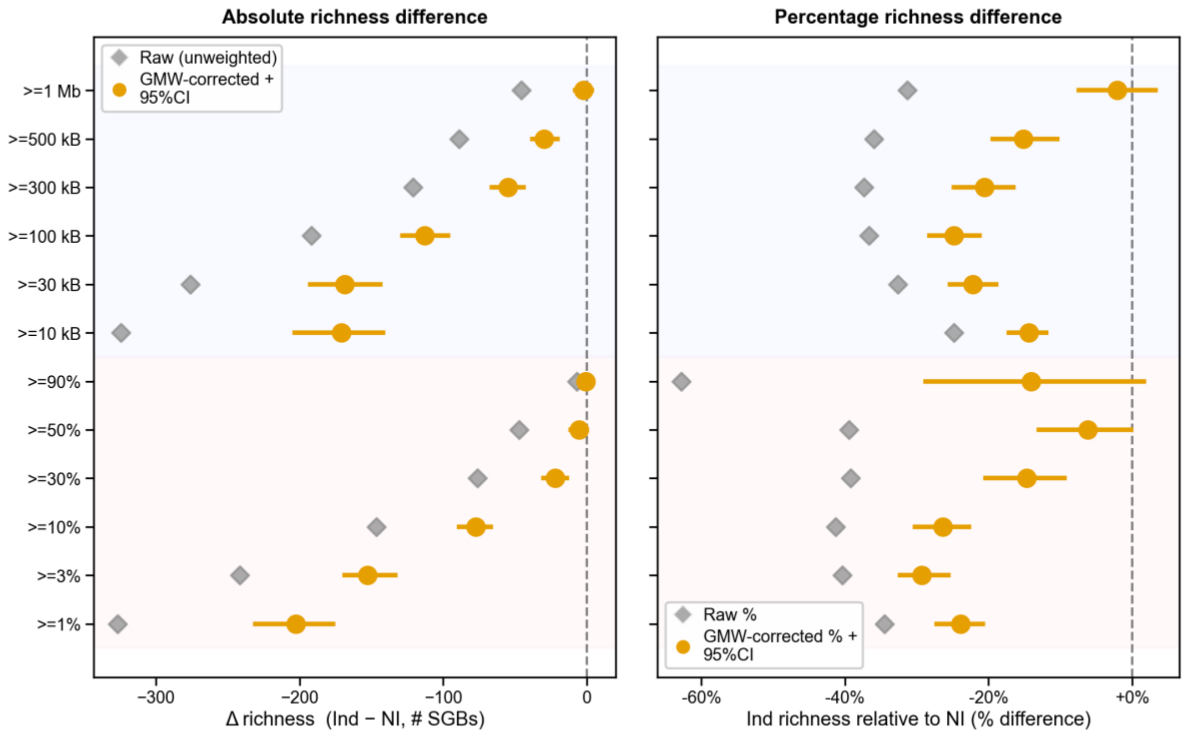

**Figure S7. Focus on samples from the Carter study: GMW-corrected richness gap between Industrial and non-Industrial samples is present at most detection thresholds.** A 3-way comparison of GMW-corrected richness between Industrial, Agro-pastoralist, and Hunter-Gatherer samples in the Carter et al. sample set finds lower Industrial richness (75-95% of non-Industrial richness). This is systematically higher than the numbers reported in **fig. S12** (~55-80%) and may result from difference in sample selection in either the Carter study or our study. There might also be more instability in depth distribution smoothing (see **fig. S8**) due to a sparser distribution of sample sequencing depth density (i.e., right-most panels show incomplete overlap between weighted distributions).

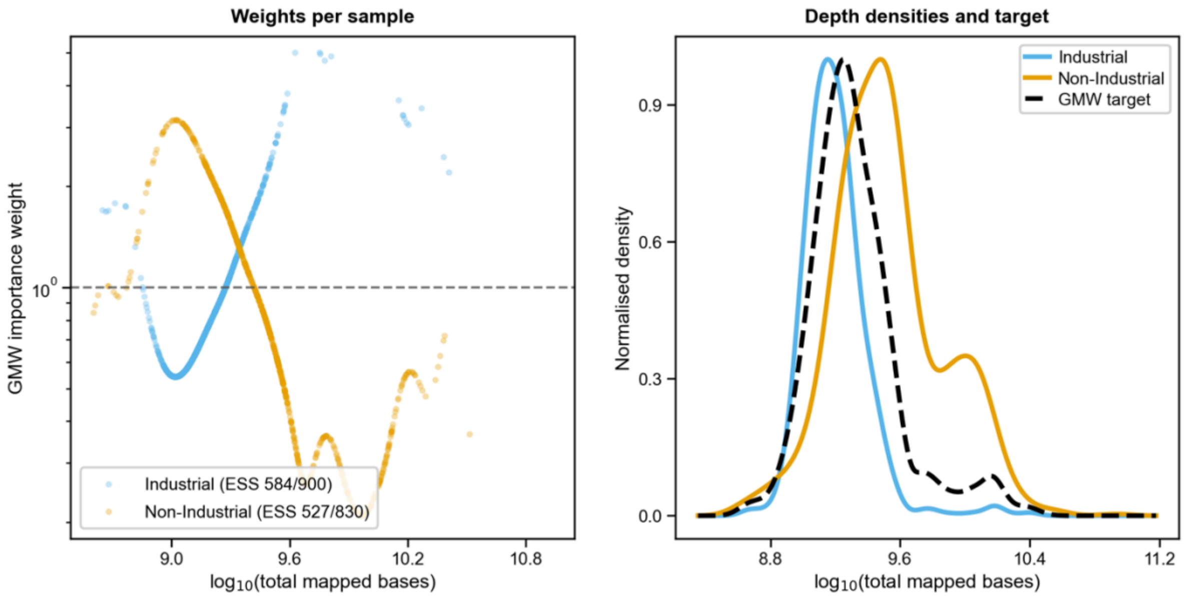

**Figure S8. Focus on samples from the Carter study: GMW weights as a function of sequencing depth.** Curves show the Industrial and Non-Industrial depth distributions in the Carter-only subset together with the corresponding GMW weights across $\log_{10}$(mapped bp). Because the Carter-only subset is smaller and has sparser coverage of the depth range than the full dataset, the weights derived as a function of sequencing depth are noisier, especially in the tails where overlap is limited. This explains the slightly less stable Carter-only robustness estimates in **figs. S5–S7** without changing their qualitative interpretation: depth correction still reduces the apparent industrial versus non-industrial richness gap.

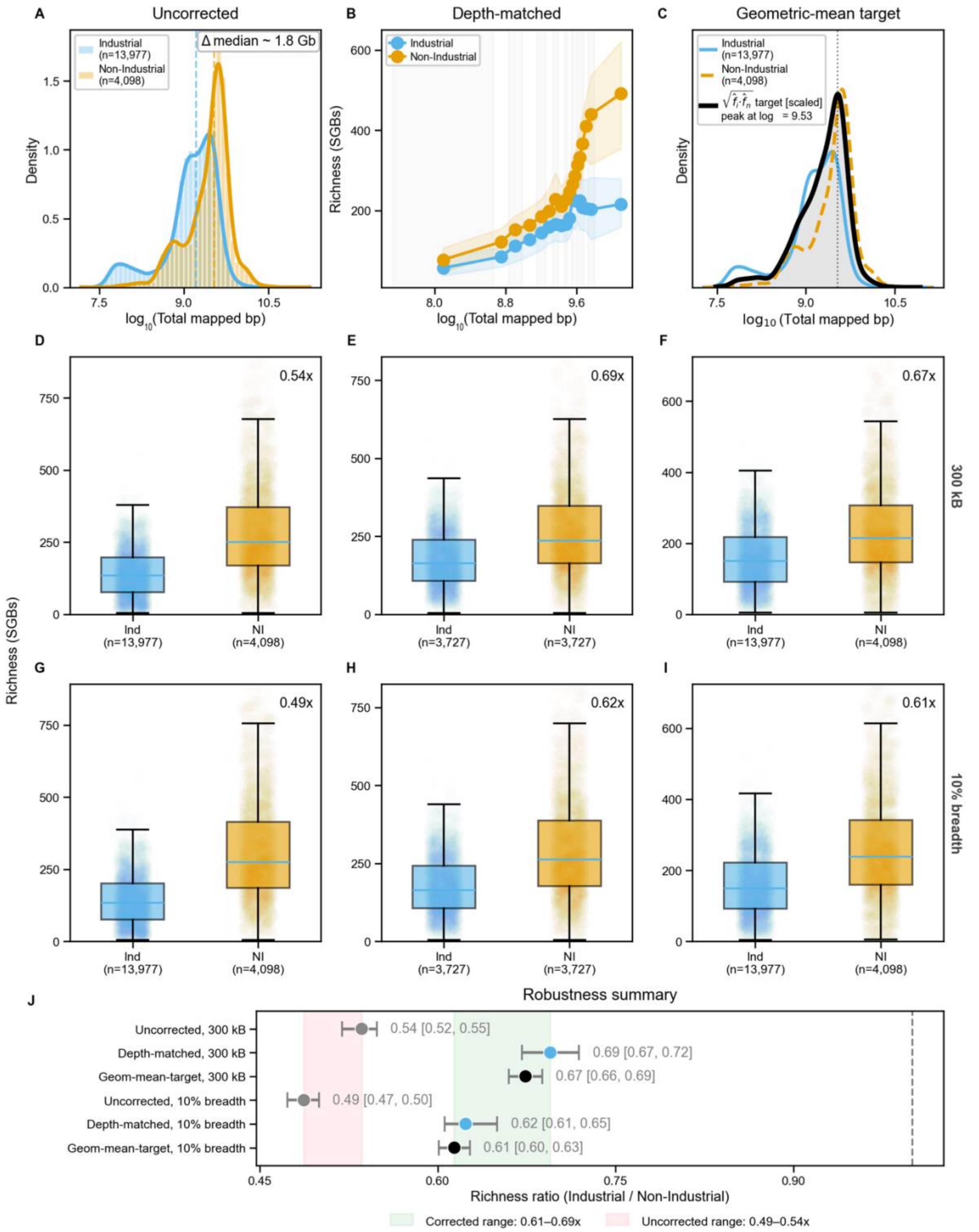

**Figure S9. Robustness of the industrial gut microbiota richness deficit across depth-correction methods and detection thresholds, based on SGBs mapped to the Carter reference database.** **(A)** Sequencing depth distributions for Industrial ($n$ = 13,977) and Non-Industrial (agro-pastoralist + hunter-gatherer; $n$ = 4,098) samples, illustrating the ~1.7 Gb median depth imbalance that motivates depth correction. **(B)** Depth-stratified median richness (±IQR ribbon) for 1:1 depth-matched pairs (≥200 kb mapped bases; 3,282 pairs), showing that the richness gap between Industrial and Non-Industrial samples persists across the full range of sequencing depths. **(C)** Depth kernel density estimates and the geometric-mean-target distribution ($\sqrt{\hat{f}_{\mathrm{Ind}} \cdot \hat{f}_{\mathrm{NI}}}$, scaled) used to derive importance weights for our geometric-mean-target weighted (GMW) correction; effective sample sizes after weighting were ESS = 11,228/14,167 (Industrial) and 2,835/3,560 (Non-Industrial). **(D–F)** Per-sample richness distributions under uncorrected, 1:1 depth-matched, and GMW-corrected comparisons, respectively, using the ≥200 kb mapped-bases detection threshold. Richness ratios (Industrial/Non-Industrial median) were 0.53 (uncorrected, 95% CI [0.51, 0.54]), 0.75 ([0.73, 0.78]), and 0.73 ([0.71, 0.74]). **(G–I)** Equivalent comparisons using the ≥10% breadth coverage threshold, yielding ratios of 0.50 ([0.48, 0.51]), 0.73 ([0.70, 0.75]), and 0.70 ([0.68, 0.71]). **(J)** Forest plot summarizing all six method × threshold combinations. After depth correction, richness ratios converged to a consistent range of 0.70–0.75 regardless of correction method or detection threshold, in contrast to the uncorrected ratios of 0.50–0.53, which reflect differences in sequencing depth rather than true biological difference. The robustness of the corrected estimates across two independent methods (1:1 matching and GMW reweighting) and two detection thresholds supports the reliability of the reported richness deficit. See **Supplementary Text 3** for GMW method details.

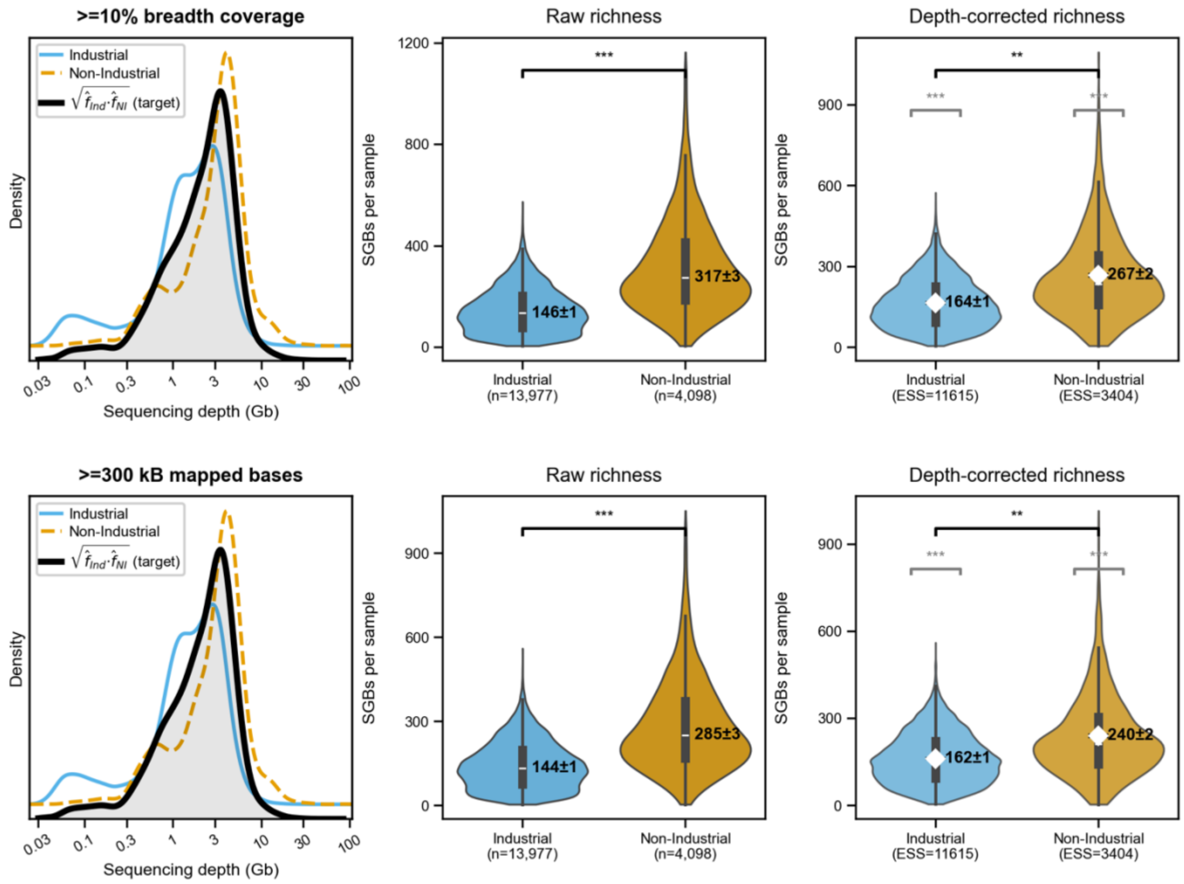

**Figure S10. Geometric-mean-target weighting (GMW) corrects for sequencing depth bias in richness comparisons.** Each row corresponds to a different taxon detection threshold (top: ≥10% breadth coverage; bottom: ≥200 kb mapped bases). Left panels show the sequencing depth density distributions for Industrial (blue) and Non-Industrial (agro-pastoralist + hunter-gatherer; gold) samples, with the smoothed GMW target distribution (black curve) overlaid. The GMW method reweights each sample in proportion to how well its depth matches this shared target, so that neither group is systematically favoured. Middle panels show violin plots of raw (uncorrected) per-sample SGB richness; right panels show the GMW-corrected richness, with weighted means (white dots ± weighted SEM) and 95% bootstrap confidence intervals. Before correction, Industrial samples appear to harbour only ~44–47% of Non-Industrial richness; after depth correction, this narrows to ~63–66%, indicating that a substantial but not complete portion of the apparent richness gap was driven by the lower sequencing depth of industrial samples. ESS = effective sample size after reweighting.

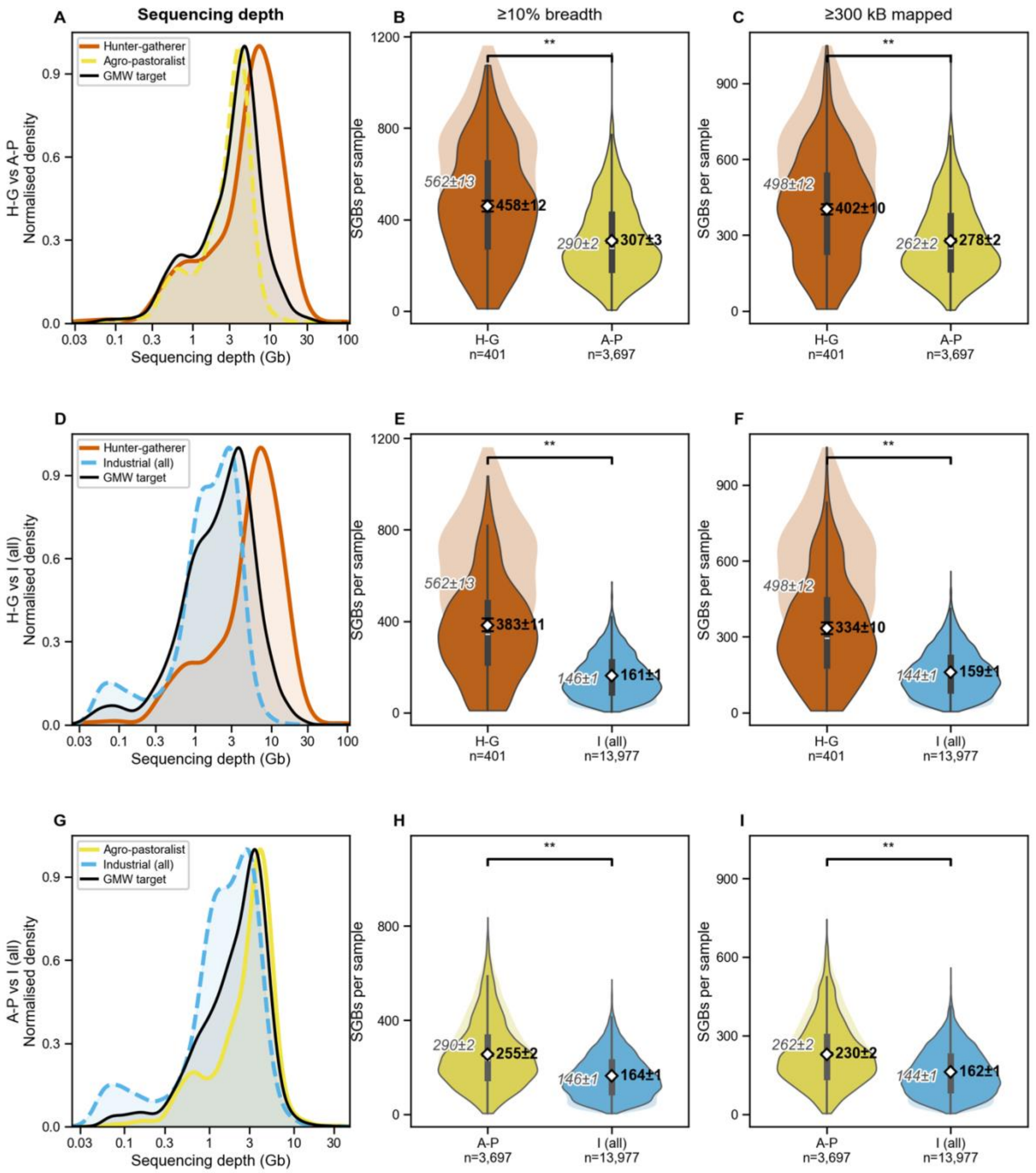

**Figure S11. 3x2-way comparison of geometric-mean-target weighting (GMW) corrects for sequencing depth bias in richness comparisons among Hunter-Gatherers (H-G), Agro-pastoralist, and Industrial (Ind) populations.** Each row corresponds to a comparison within a pair of lifestyle groups. Left panels show the sequencing depth density distributions for Industrial (blue), Agro-pastoralist (yellow) samples, with the smoothed GMW target distribution (black curve) overlaid. The GMW method reweights each sample in proportion to how well its depth matches this shared target, so that neither group is systematically favoured.

Middle panels show violin plots of raw (uncorrected) per-sample SGB richness; right panels show the GMW-corrected richness, with weighted means (white dots ± weighted SEM) and 95% bootstrap confidence intervals. The results are slightly different to those displayed in **Fig. 2A-C in the main text**, where the reweighting target is chosen as the geometric mean of all 3 lifestyle groups simultaneously, rather than for each of the pairwise comparisons shown here.

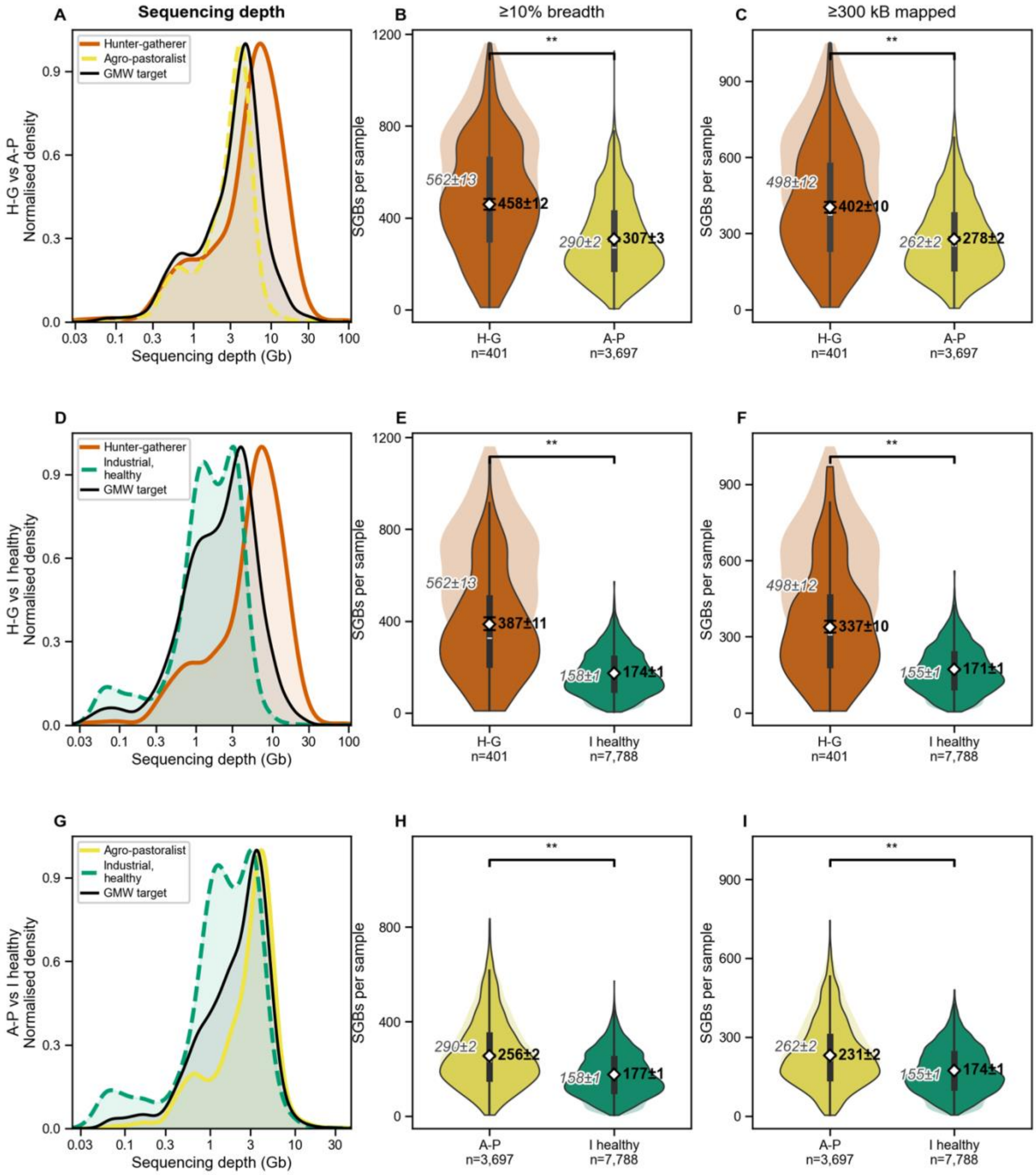

**Figure S12. 3-way comparison of geometric-mean-target weighting (GMW) corrects for sequencing depth bias in richness comparisons between Hunter-Gatherers (H-G), Agro-pastoralist, and Healthy Industrial (IndH) populations.** Each row corresponds to a comparison within a pair of lifestyle groups. Here we subset the Industrial populations into a subset of healthy individuals, and find that the GMW-corrected healthy industrial distributions are somewhat closer to the agro-pastoralist and hunter-gatherer richness distributions than the overall industrial distribution was found to be in the previous figure (**fig. S11**).

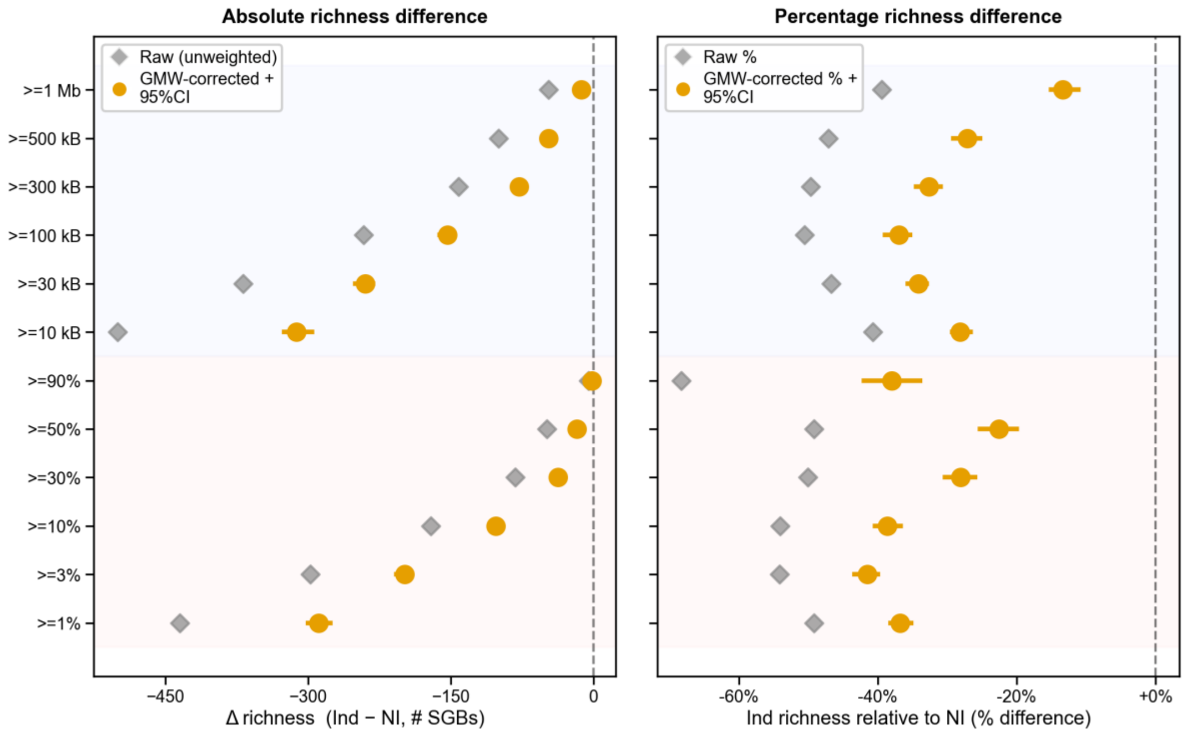

**Figure S13. GMW-corrected richness gap is robust across all detection thresholds.** Forest plot showing the absolute (left) and relative (right) richness difference between Industrial and Non-Industrial samples across twelve taxon-detection thresholds: six breadth-coverage cutoffs (1–90% of the genome covered) and six flat minimum-mapped-base cutoffs (10 kb–1 Mb). Grey diamonds show the raw (uncorrected) difference; orange and blue circles with 95% bootstrap confidence intervals show the GMW depth-corrected estimates. Across all thresholds, the corrected estimates are consistently less negative than the raw values. This confirms that lower sequencing depth in industrial samples inflates the apparent richness gap. Even so, all corrected estimates remain substantially negative, with richness in Industrial populations being 20–45% below richness in Non-Industrial populations, depending on threshold).

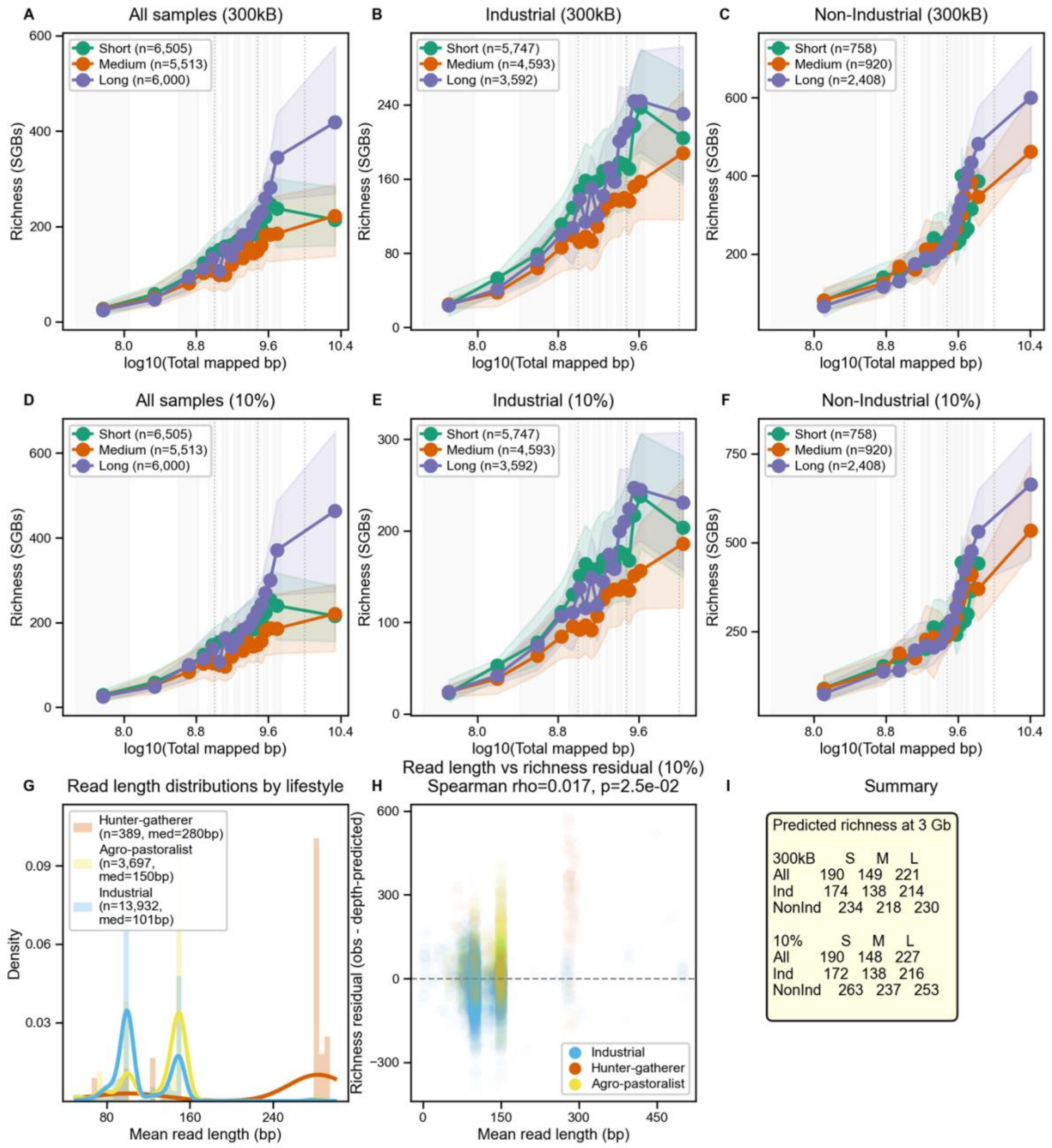

**Figure S14. Read length has a minor but detectable effect on observed richness.** (**A–F**) Per-sample richness (SGBs detected) as a function of sequencing depth ($\log_{10}$ total mapped bp), with samples split into three read-length terciles: Short (≤99 bp, green), Medium (99–111 bp, orange), and Long (>111 bp, purple). The top row (**A–C**) applies the alternative flat detection rule of 300 kb mapped to an SGB; the bottom row (**D–F**) applies the primary rule of ≥10% genome-relative breadth required for a detection. Each rule is shown for all samples combined (**A, D**), industrial samples only (**B, E**), and non-industrial samples only (**C, F**). Longer reads detect modestly more taxa at equivalent sequencing depth. (**G**) Industrial and Agro-pastoralist samples are overwhelmingly sequenced with shorter reads (~100 bp) while H-G samples have longer reads (~150 bp median), creating a potential confound. (**H**) After controlling for sequencing depth, the residual association between read length and richness is small and positive (Spearman $\rho$ = +0.017, $p$ = 0.025). However, because many samples sequenced with the longest reads were also sequenced with the greatest depth (e.g., Hadza and Nepali samples from Carter et al.), correction for sequencing depth cannot fully eliminate confounding due to study batch effects. (**I**) Predicted richness at 3 Gb sequencing depth by read-length tercile and lifestyle. Differences across terciles are modest relative to the between-lifestyle gap, indicating that read length is not a primary driver of the richness differences reported in the main text.

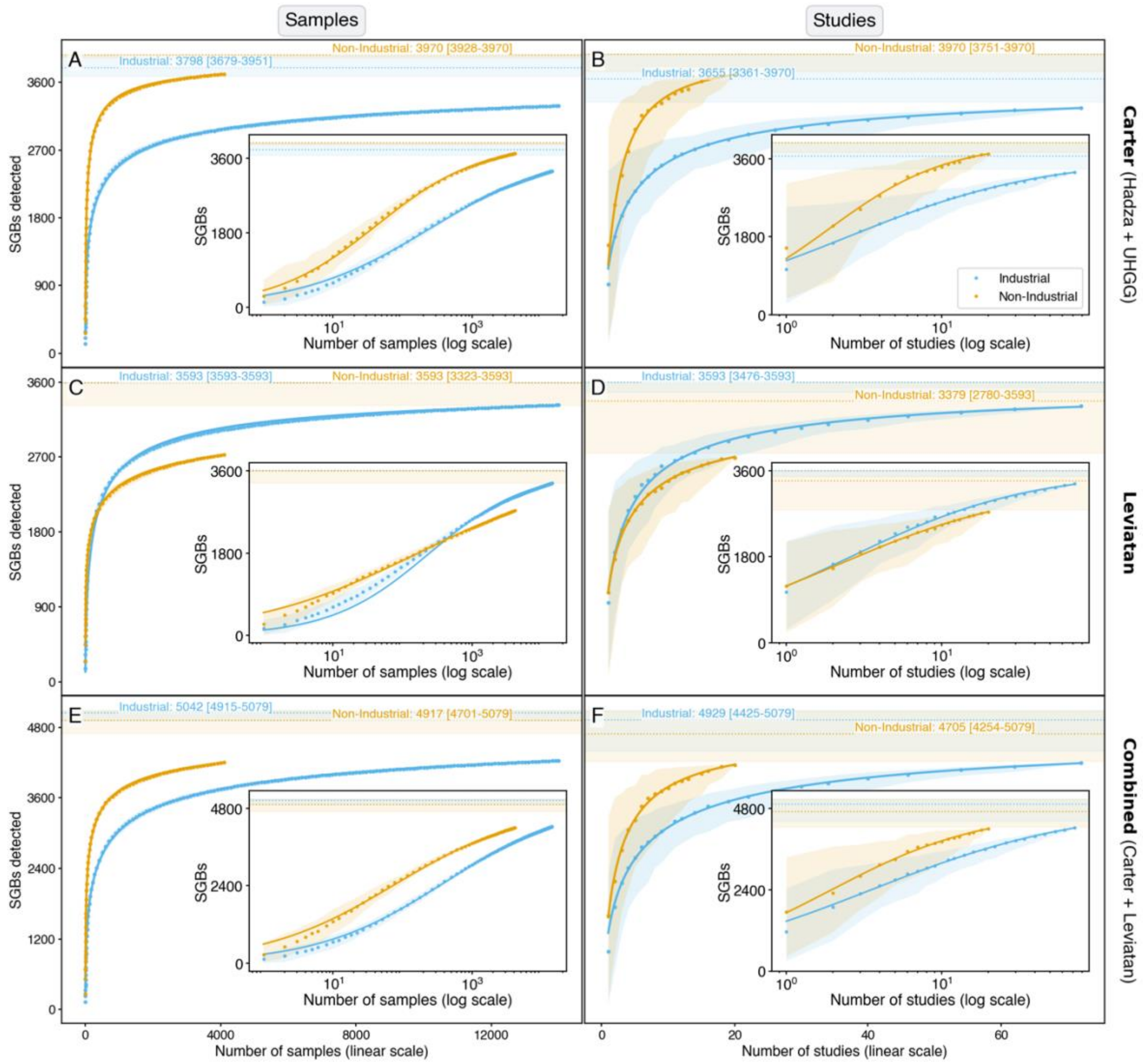

**Figure S15. Extensive microbial extinction has not occurred in industrial populations (empirical Hill-type Michaelis-Menten fit). Left panels**: Expected number of SGBs detected as a function of number of pooled **samples**. **Right panels**: Expected number of SGBs detected as a function of number of pooled **studies**. Outer plots are shown in linear space; inner insets are shown in log-linear space. Curves show the bootstrap mean with 95% quantile ribbons; the fitted line **[3-parameter Hill-type Michaelis-Menten]** is shown within the observed range. Asymptote annotations give the fitted convergence value (median) with 95% bootstrap CI. **(A, B)** Data for SGB accumulation using the Carter reference (Hadza + UHGG) biased towards non-industrial samples. **(C, D)** Data for SGB accumulation using the Leviatan reference biased towards industrial samples. **(E, F)** Data for SGB accumulation using the combined Carter + Leviatan reference (see **Supplementary Text 2D** for details). At low degrees of sample pooling both references detect more SGBs in non-industrial populations; at higher degrees of pooling, the pattern diverges according to database composition, with the Carter reference continuing to favor higher diversity in non-industrial populations and the Leviatan reference crossing over to favor higher diversity in industrial populations. At the very high sample count (*N*>100), the absolute difference between the two references diminishes in the Carter and combined references, indicating that rare microbes found in non-industrial populations and incorporated into non-industrial biased references are detected in sufficiently broad pooling of industrial samples.

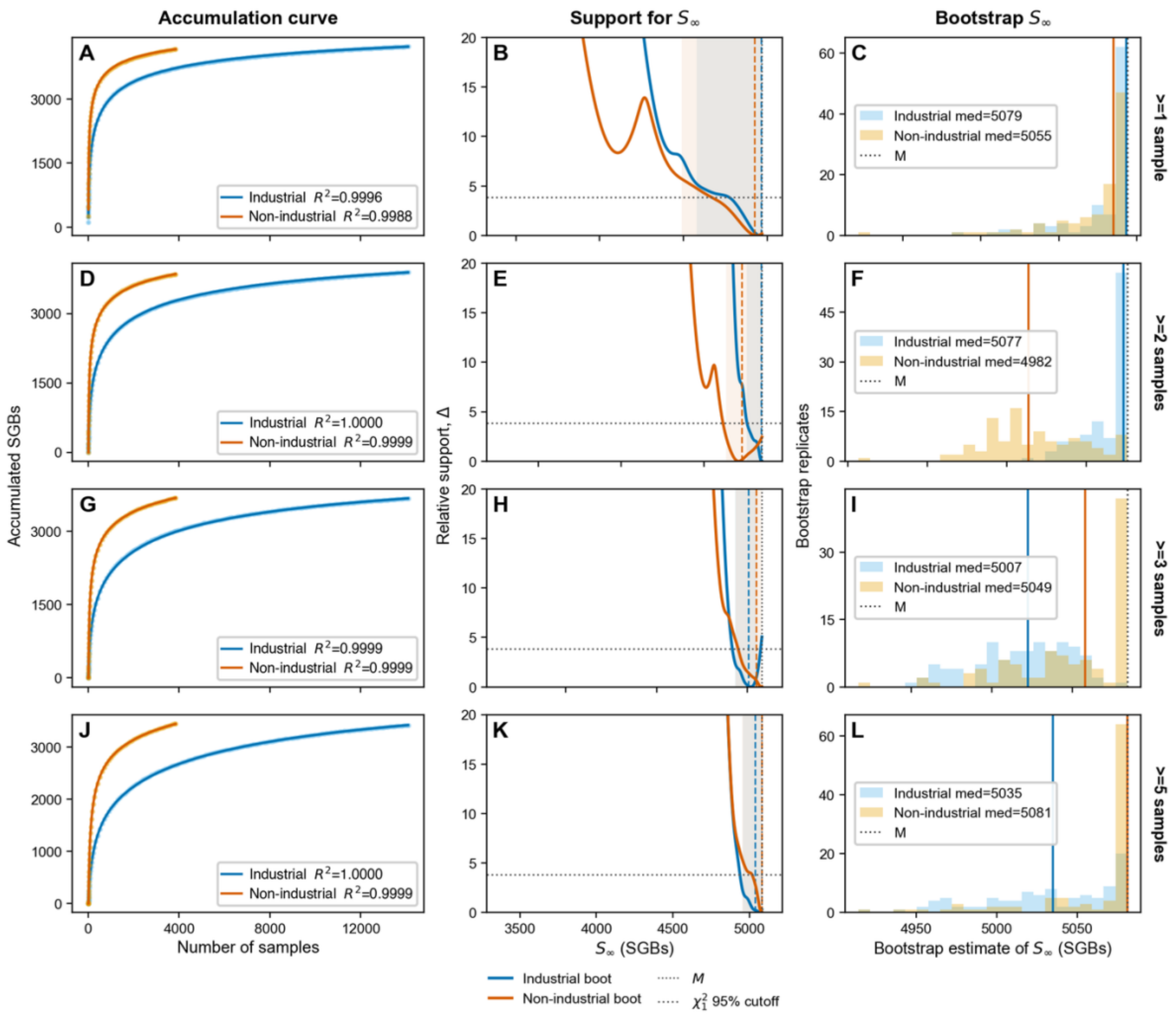

**Figure S16. Accumulation of reference SGBs when SGBs were required to be detected in at least $k$ samples.** Reads mapped to the Combined reference. Rows show $k = 1,2,3,$ and 5. For $k = 1$, accumulation is a first-detection problem and was fitted with the best empirical first-detection curve among Michaelis-Menten, generalized Michaelis-Menten, Hill-type Michaelis-Menten, Richards 4-parameter sigmoid, and 1, 2, or 3-population beta-mixture models. Best fit was either 2-population beta-mixture model or Richards 4-parameter sigmoidal fit. For $k \geq 2$, accumulation was fitted using finite-catalogue beta-mixture curves, which explicitly model the probability that each SGB is detected repeatedly across samples. Left panels (**A,D,G,J**) show bootstrap-median accumulation curves and fitted curves; shaded intervals indicate bootstrap spread. Middle panels (**B,E,H,K**) show support for possible asymptotes $S_\infty$, with the dotted horizontal line marking $\Delta = 3.84$. Right panels (**C,F,I,L**) show bootstrap distributions of inferred $S_\infty$, with colored vertical lines marking medians and the dotted black line marking the full catalogue size $M$ . The inferred asymptotes remain close to $M$ across $k = 1,2,3,$ and 5, supporting the conclusion that most combined-catalogue SGBs are expected to be eventually detectable.

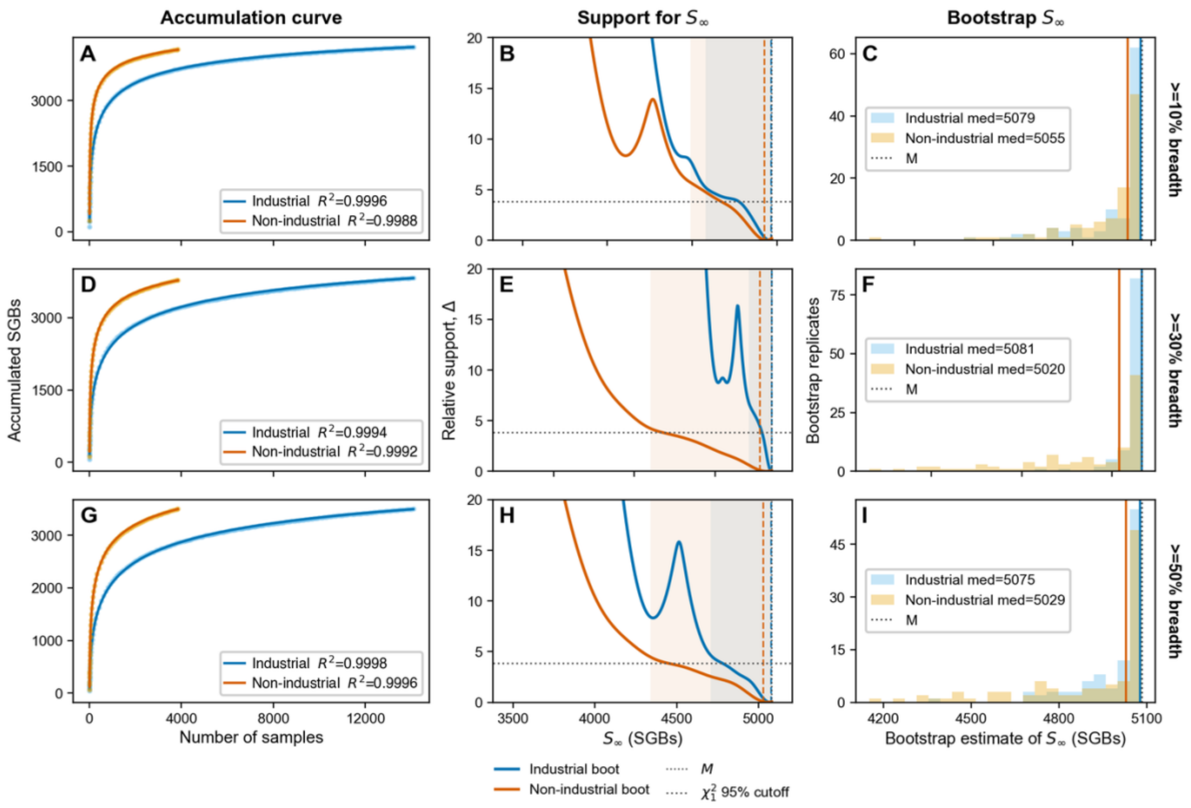

**Figure S17. Accumulation of reference SGBs in Industrial and Non-Industrial samples using increasingly stringent per-sample detection thresholds.** Reads mapped to the Combined reference. Rows show SGB accumulation when an SGB was counted as detected only if at least **10%**, **30%**, or **50%** of the SGB was recovered in a sample. Left panels (**A,D,G**) show the bootstrap-median accumulation curve, with shaded intervals indicating the bootstrap spread, and fitted empirical first-detection curves. This was fitted with the best empirical first-detection curve among Michaelis-Menten, generalized Michaelis-Menten, Hill-type Michaelis-Menten, and Richards models, and 1, 2, or 3-population beta-mixture models. Best fit was either 2-population beta-mixture model or Richards 4-parameter sigmoidal fit. $R^2$ values quantify agreement between the fitted curve and the bootstrap-median accumulation curve. Middle panels (**B,E,H**) show support for alternative finite-catalogue asymptotes $S_\infty$, where lower $\Delta(S_\infty)$ indicates greater support and the dotted horizontal line marks the approximate 95% profile threshold, $\Delta = 3.84$. Right panels **(C,F,I)** show the distribution of bootstrap estimates of $S_\infty$; vertical colored lines mark bootstrap medians and the dotted black line marks the full combined reference catalogue size $M$. Across detection thresholds, inferred asymptotes remain close to the full catalogue size, indicating that the near-catalogue asymptote is not driven by permissive SGB detection criteria.

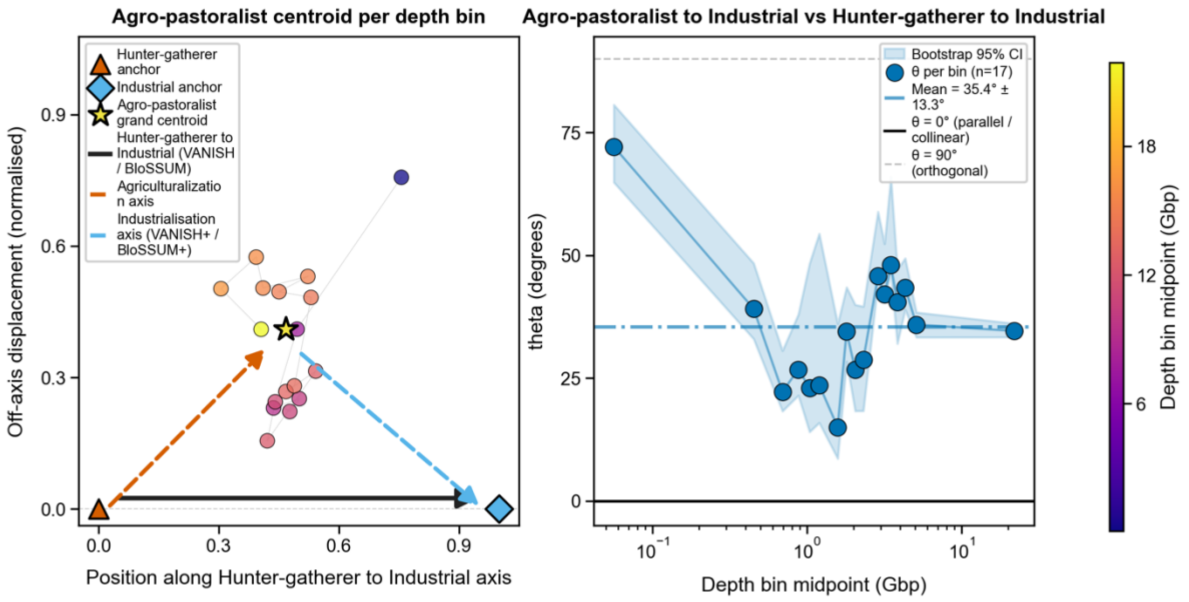

**Figure S18. Coordinate-invariant projection of Agro-pastoralist populations into Hunter-Gatherer→Industrial axis space, aggregated by lifestyle and reported by sequencing depth bin.** The study centroid is shown for all matched samples projected onto a normalized axis whose endpoints are the Hunter-Gatherer and Industrial centroids within the corresponding depth bin. The x-axis gives the sample's fractional position along this Hunter-Gatherer→Industrial axis (0 = Hunter-Gatherer centroid, 1 = Industrial centroid), while the y-axis gives signed off-axis displacement, oriented so that positive values lie on the Agro-pastoralist side. Agro-pastoralist samples are intermediate along the main Hunter-Gatherer→Industrial axis but remain consistently displaced off that axis regardless of bin considered, indicating that the shift into agro-pastoralist microbiome composition is not collinear with the direct Hunter-Gatherer→Industrial contrast. The displacement to the Hunter-Gatherer→Industrial axis is consistent with sequencing depth.

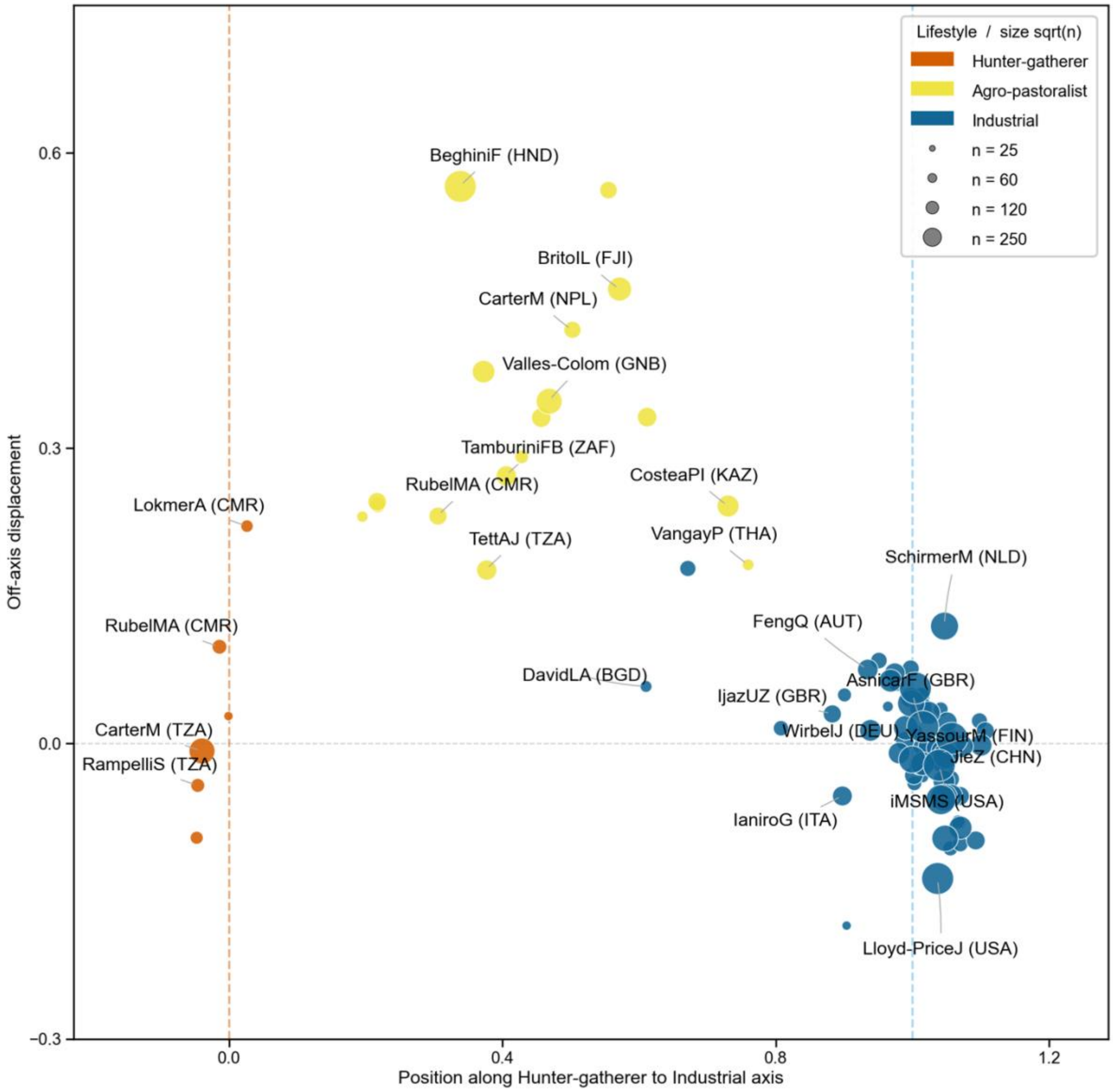

**Figure S19. Coordinate-invariant projection of samples matched in 17 separate depth bins (see fig. S18) into Hunter-Gatherer→Industrial axis space, aggregated by study.** The study centroid is shown for all matched samples projected onto a normalized axis whose endpoints are the Hunter-Gatherer and Industrial centroids within the corresponding depth bin. The x-axis gives the sample's fractional position along this Hunter-Gatherer→Industrial axis (0 = H-G centroid, 1 = Industrial centroid), while the y-axis gives signed off-axis displacement, oriented so that positive values lie on the Agro-pastoralist side. Agro-pastoralist samples are intermediate along the main Hunter-Gatherer→Industrial axis but remain consistently displaced off that axis, indicating that the shift into agro-pastoralist microbiome composition is not collinear with the direct Hunter-Gatherer→Industrial contrast. A few populations are clearly borderline between the two categories (e.g. PehrrsonEC (El Salvador, Peru), VangayP (Thailand), and TamburiniFB (South Africa). These populations consist of peri-urban populations in developing countries that are likely not fully "industrialized". We note that they are not sequenced particularly deeply and broadly and we do not expect them to lead to substantial misindentification of SGBs in our accumulation analysis.

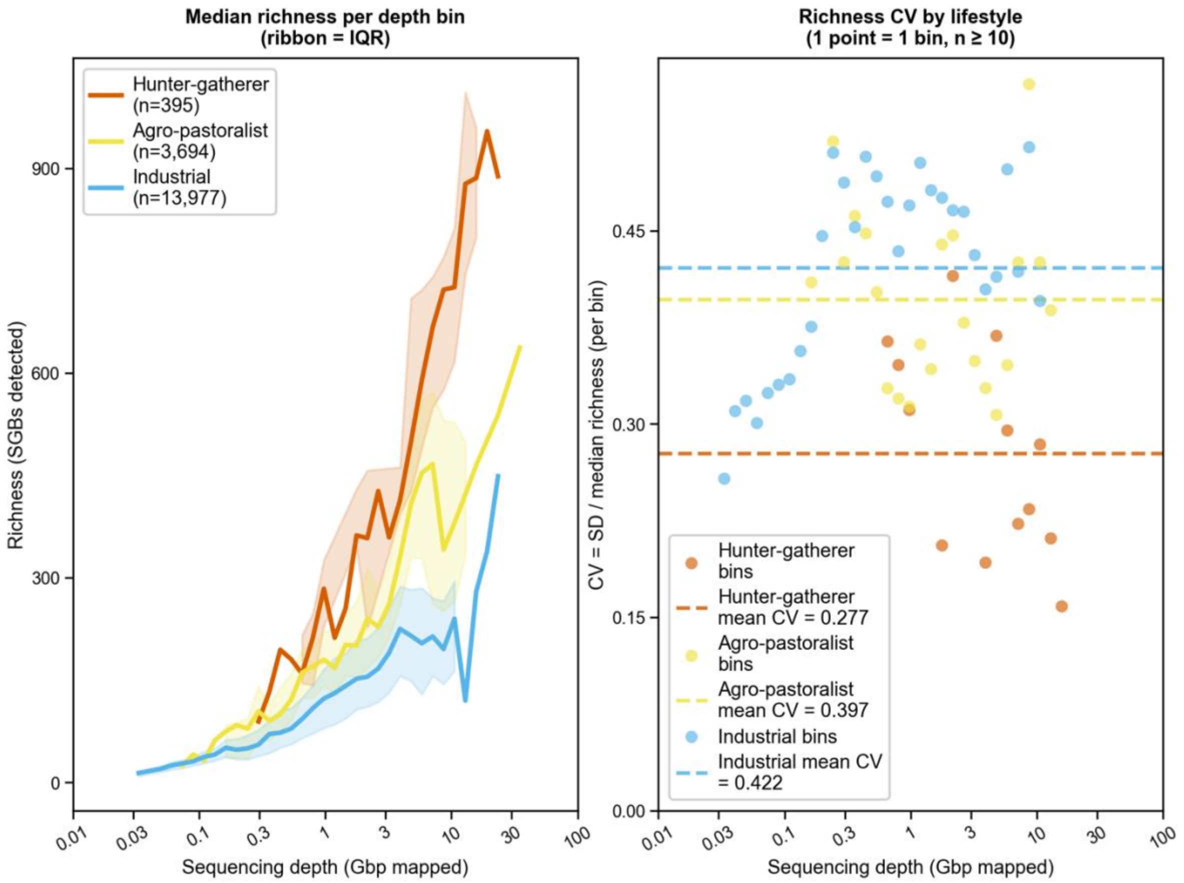

**Figure S20. Depth-controlled richness variability within lifestyle groups.** Samples were assigned species presence using the ≥10% breadth coverage threshold and binned into 40 equally log-spaced sequencing depth bins (x-axis; Gbp mapped). Left panel: median richness per bin by lifestyle, with ribbon showing the interquartile range; the positive depth–richness relationship is consistent across all three lifestyles (Spearman $\rho$ = 0.80, 0.68, and 0.65 for H-G, Agro-pastoralist, and Industrial, respectively). Right panel: coefficient of variation (CV = SD/median richness) per occupied bin (minimum 10 samples per bin), colored separately by lifestyle. Grand mean CV differed significantly across lifestyles (H-G = 0.27 across 13 qualifying bins, Agro-pastoralist = 0.40 across 26 bins, Industrial = 0.45 across 33 bins; Kruskal–Wallis $H = 29.66$, $p = 3.6 \times 10^{-7}$; permutation test on maximum pairwise mean-CV difference: observed $\Delta = 0.18$, $p = 0.0002$), with all pairwise comparisons significant after Benjamini-Hochberg (BH) correction (H-G vs. Agro-pastoralist: $p_{BH} = 2.8 \times 10^{-4}$, $r = 0.75$; H-G vs. Industrial: $p_{BH} = 3.6 \times 10^{-6}$, $r = 0.93$; Agro-pastoralist vs. Industrial: $p_{BH} = 0.003$, $r = 0.45$). The lower CV in H-G samples likely reflects the smaller number of contributing studies and the narrower geographic range of sampled hunter-gatherer populations, rather than lower true biological variability. Crucially, CV values of 0.27–0.45 persist across the full depth range in all three lifestyle groups, confirming that substantial richness variation remains after controlling for sequencing depth and is not a depth artefact.

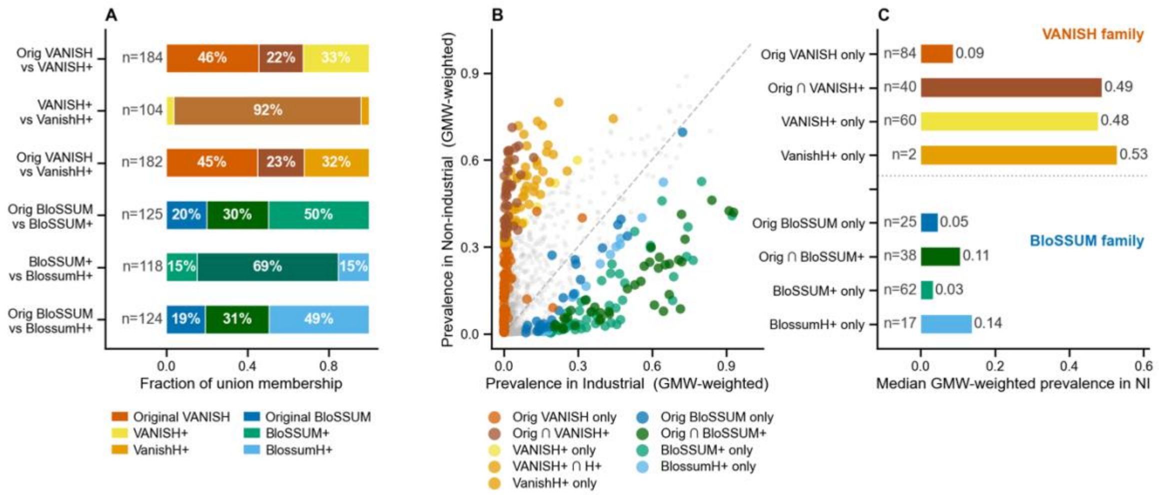

**Figure S21. Pairwise overlap among original (VANISH/BloSSUM), industrialization-defined (VANISH$^{+}$, BloSSUM$^{+}$), and H+-restricted (VANISH$^{H+}$, BloSSUM$^{H+}$) taxon lists.** The figure summarizes overlap within each family (VANISH on the left; BloSSUM on the right) between the original Carter lists, our "+" lists defined from the industrial versus non-industrial comparison in the present dataset, and our "H+" lists defined from the non-industrial versus healthy-industrial comparison in the present dataset. Labels report list sizes and pairwise overlap, and the Jaccard index quantifies similarity as $|A \cap B|/|A \cup B|$. High overlap between the + and H+ lists indicates continuity within each family, whereas the more modest overlap between the original and +/H+ lists shows that these new lists capture related but non-identical aspects of lifestyle-associated prevalence structure.

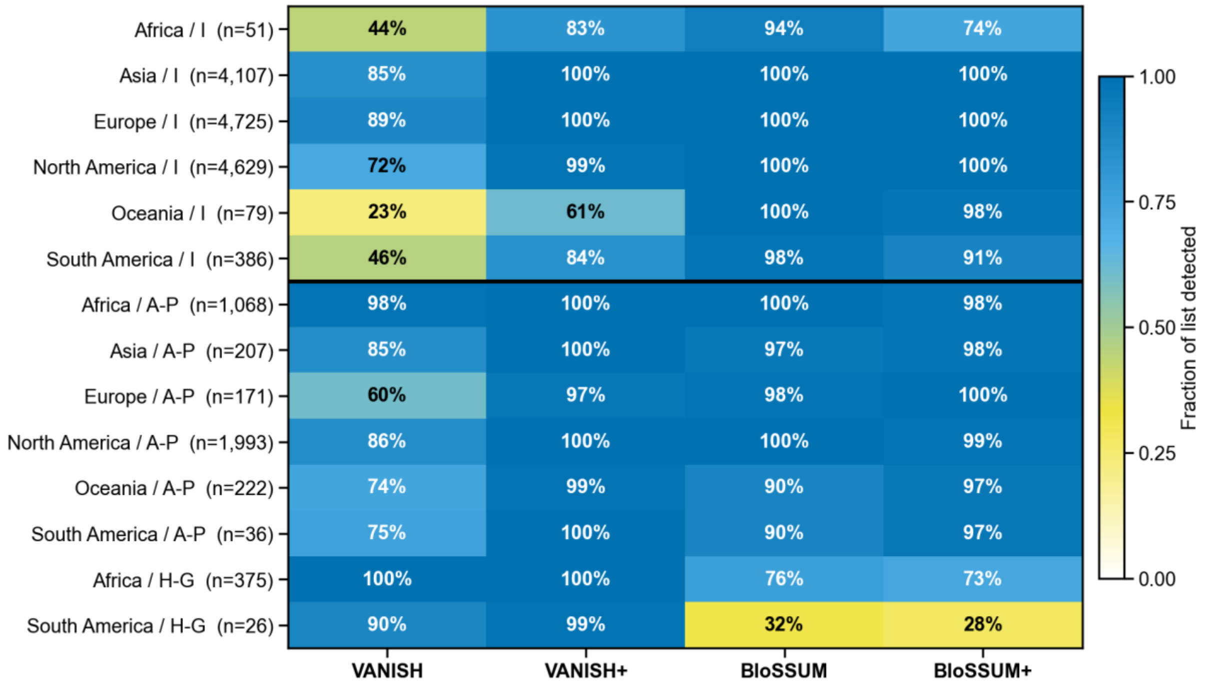

**Figure S22. Breakdown of VANISH/BloSSUM and VANISH$^{+}$/BloSSUM$^{+}$ taxa detection by continent and lifestyle.** Most categories recover most of the taxa. The exceptions are the categories with low sequencing depth (e.g. hunter-gatherers in South America). Because these cohorts are not balanced or corrected for sequencing depth, so cross-interpretation is not intended between cohorts, only within cohorts. So, we can read that hunter-gatherers recover fewer of the BloSSUM than the VANISH group taxa, and industrial populations fewer of the VANISH than the BloSSUM, at the same sequencing depth.

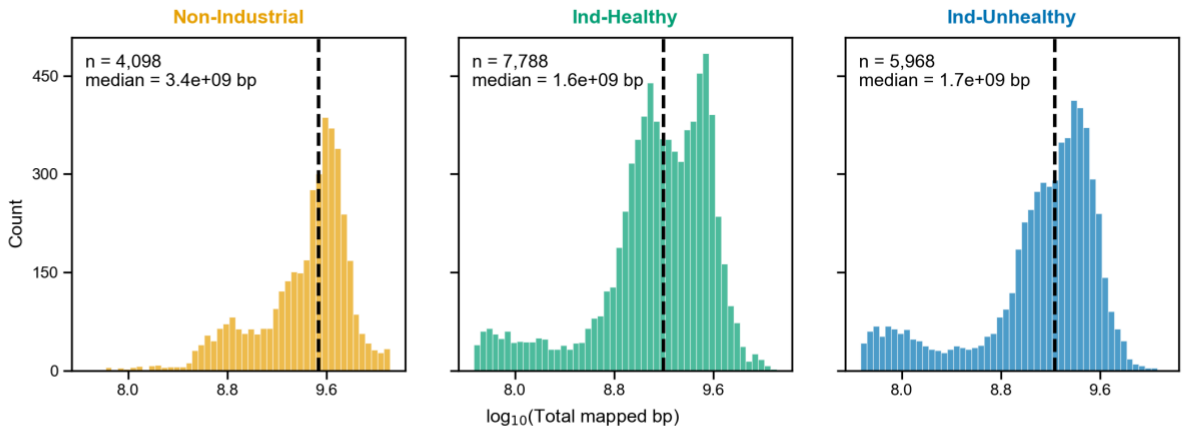

**Figure S23. Sequencing depth distributions by health group, before depth-matching.** Histograms of total bacterial bases mapped per sample ($\log_{10}$ scale) stratified by health status among industrial and non-industrial populations: Non-Industrial (orange, *n*=4,098), Industrial-Healthy (green, *n*=7,788), Industrial-Unhealthy (blue, *n*=5,968). A small number of samples with unclear health status Industrial-Unknown (*n*=221) were not plotted. Dashed lines show group medians. Non-Industrial samples are sequenced more deeply on average (median ~$3.4\times10^9$ bp) than Industrial samples in either health category (~$1.7\times10^9$ bp), underscoring the importance of depth-matching in the health association analyses.

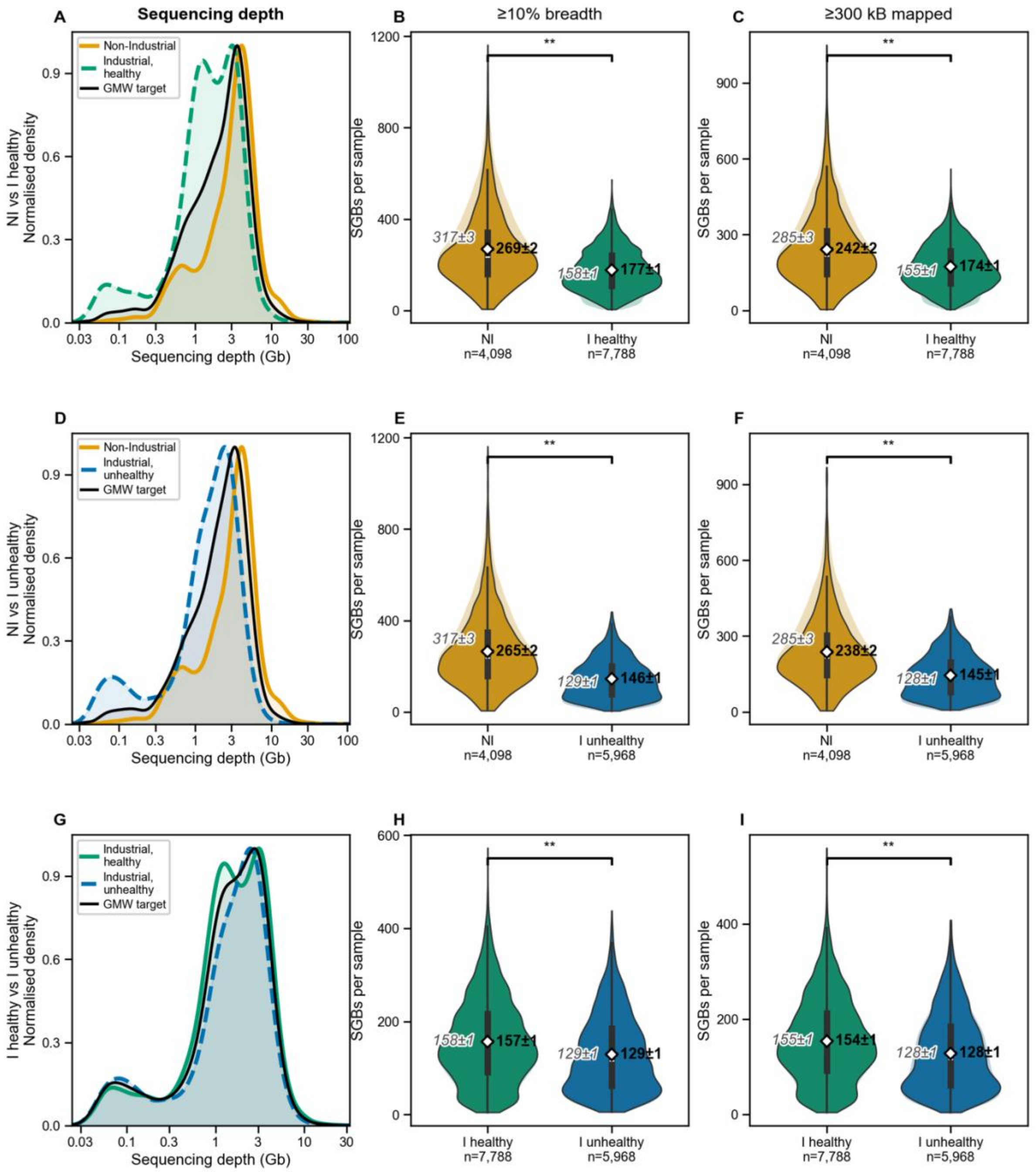

**Figure S24. 3x2-way comparison of geometric-mean-target weighting (GMW) corrects for sequencing depth bias in richness comparisons between Non-Industrial, Healthy Industrial, and Unhealthy Industrial populations.** Each row corresponds to a comparison within a pair of lifestyle groups. Left panels show the sequencing depth density distributions for Industrial Healthy (dark green), Industrial Unhealthy (dark blue) and non-Industrial (gold) samples, with the smoothed GMW target distribution (black curve) overlaid. The GMW method reweights each sample in proportion to how well its depth matches this shared target, so that neither group is systematically favoured. Middle panels show violin plots of raw (uncorrected) per-sample SGB richness; right panels show the GMW-corrected richness, with weighted means (white dots ± weighted SEM) and 95% bootstrap confidence intervals. The results are slightly different to those displayed in **Fig. 5A-C in the main text**, where the reweighting target is chosen as the geometric mean of all 3 lifestyle groups simultaneously, rather than for each of the pairwise comparisons shown here.

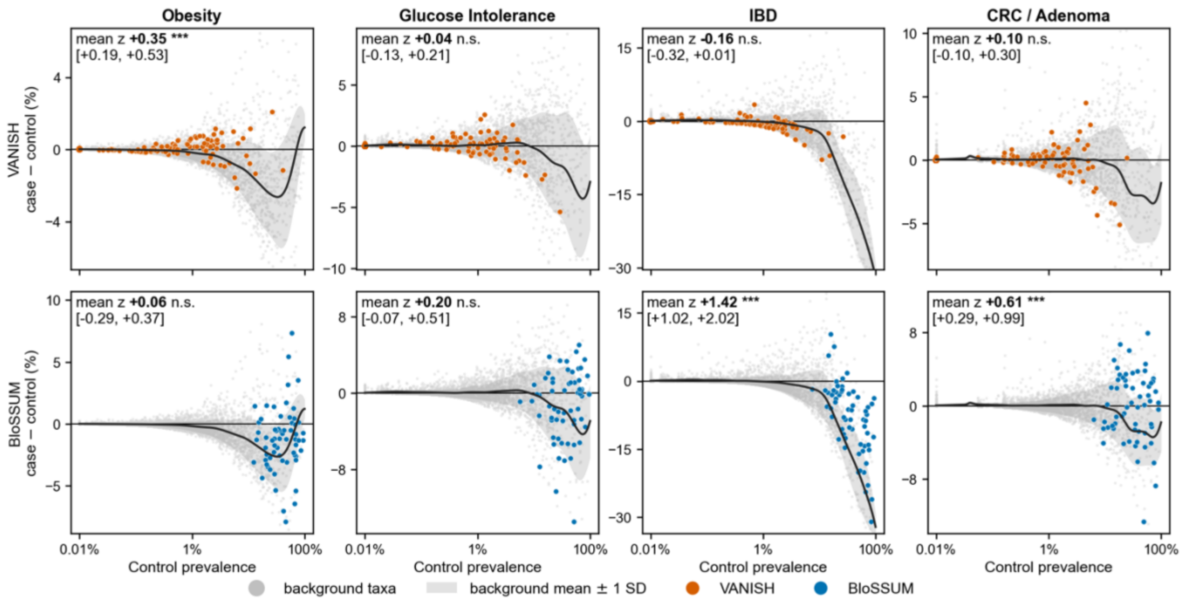

**Figure S25. The prevalence background, original VANISH and BloSSUM lists.** For each of the four reported conditions, every background taxon's study-stratified case−control prevalence difference is plotted against its control prevalence (grey points), with the fitted background curve and its ±1 SD band. List taxa are drawn on top: VANISH in orange (upper row) and BloSSUM in blue (lower row). Each panel prints the list's mean distance from the curve in units of the local spread, with a bootstrap 95% interval that refits the curve on resampled background taxa, and stars from the two-sided bootstrap p-value under the 10% breadth rule. Taxa with fewer than 15 comparable background taxa within 0.25 $\log_{10}$ of their control prevalence are plotted but excluded from the mean. The curve turns down at high prevalence because a taxon present in almost every control has room to fall and none to rise; the question each panel asks is whether the list sits above or below that expectation rather than above or below zero. Main-text **Fig. 5E** is the same figure for the VANISH$^{H+}$ and BloSSUM$^{H+}$ lists.

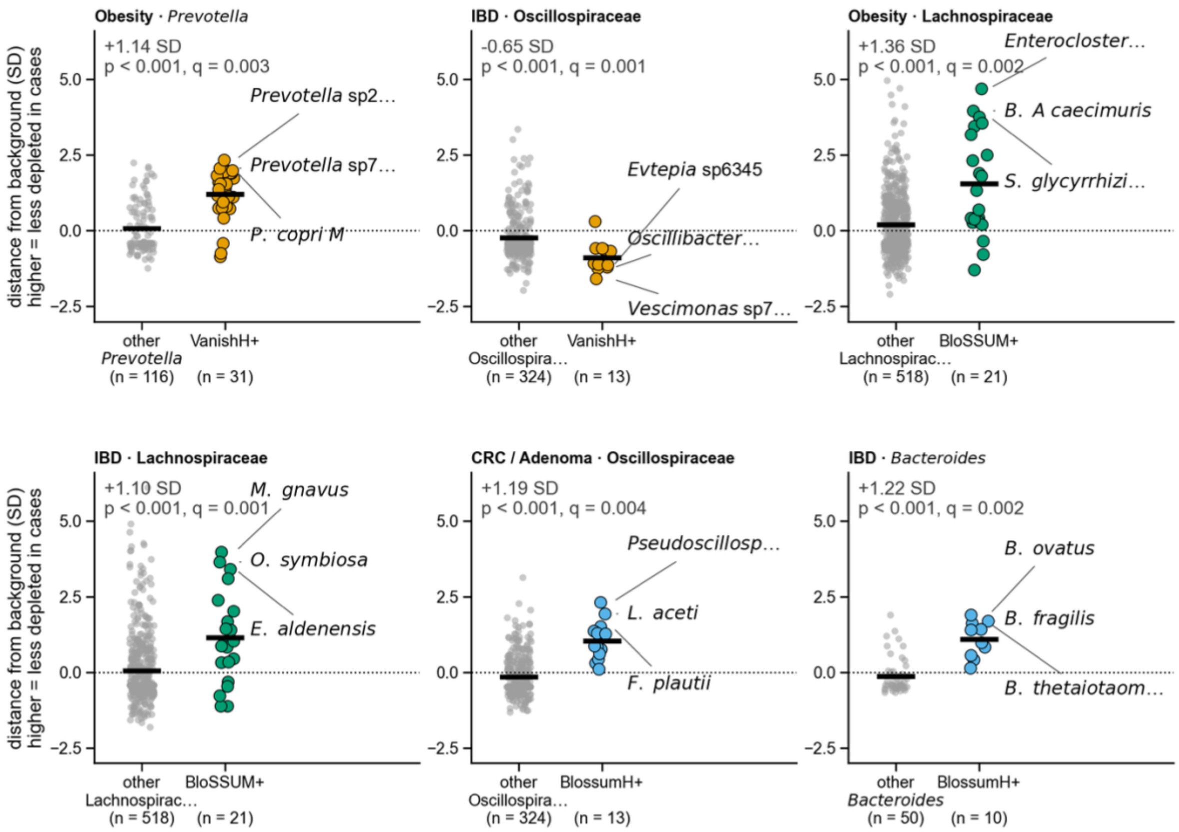

**Figure S26. Within one genus or family, list members behave differently from their relatives.** Each panel takes one disease condition and one clade, and compares the distance from the prevalence background of the clade's list members (coloured) against every other catalogue taxon in the same clade (grey); higher values mean a taxon is less depleted in cases than its baseline prevalence predicts. The background curve is refitted with the clade excluded before scoring, so a large clade is not partly its own control, and only clades with at least five catalogue taxa are tested. Horizontal bars mark group means, and each panel prints the difference between them in SD units with its permutation p and Benjamini–Hochberg q across clades; the three most extreme list members are named. Fifty-seven of the 340 clade-by-condition contrasts clear $q < 0.05$, so the six panels are a selection: each is among the six most significant contrasts for its own list, and together they span both list families and three conditions. They are *Prevotella* under obesity and Oscillospiraceae under IBD for VANISH$^{H+}$, Lachnospiraceae under obesity and under IBD for BloSSUM$^{+}$, and Oscillospiraceae under CRC / adenoma and *Bacteroides* under IBD for BloSSUM$^{H+}$; clades with a larger difference but only a few list members, such as the three BloSSUM$^{+}$ Coprobacillaceae in IBD (+2.5 SD), are not drawn.

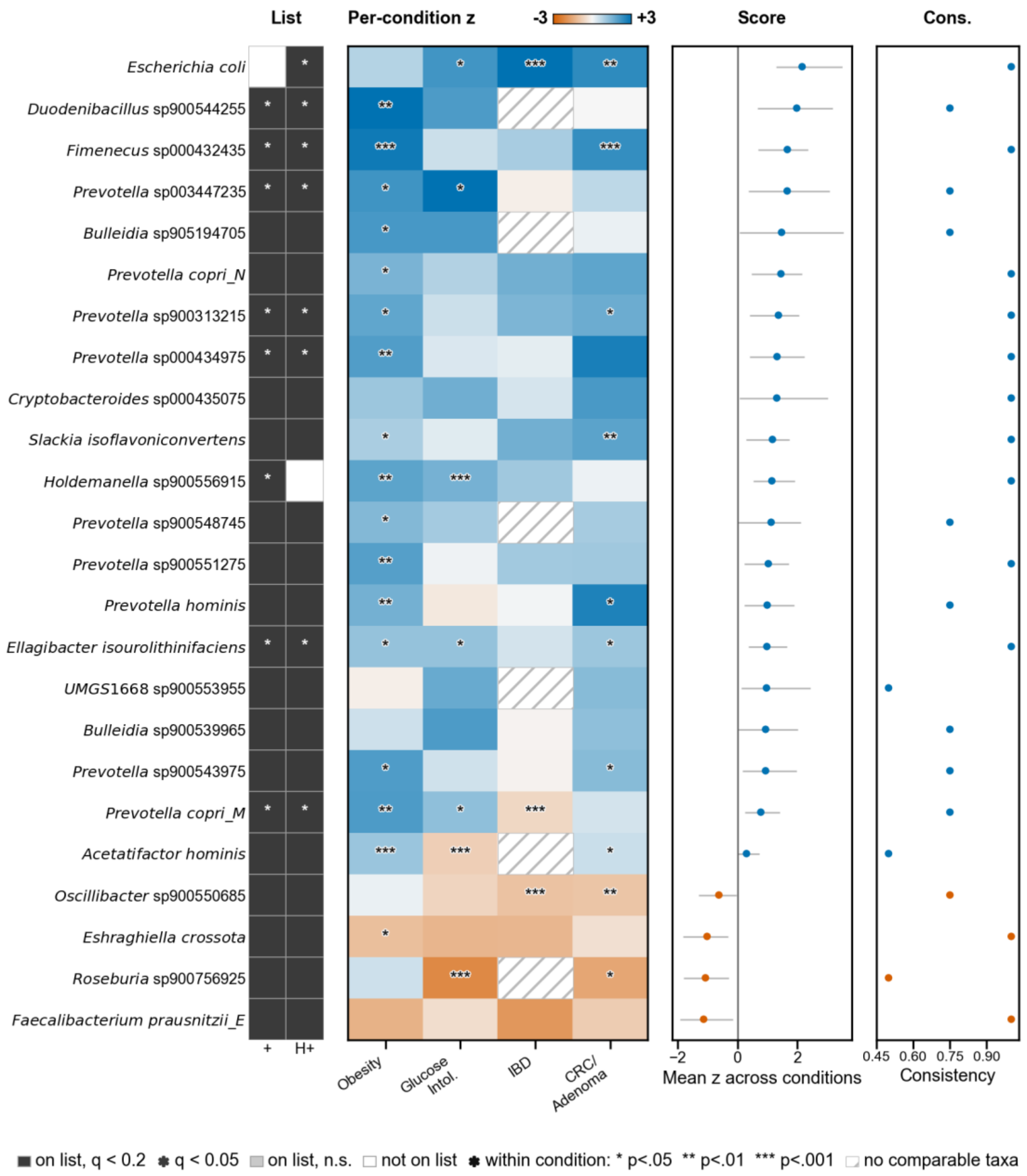

**Figure S27.** Taxa carrying the VANISH-family health associations. The VANISH$^{+}$ and VANISH$^{H+}$ lists share 96 of their 100 taxa, so they are shown together; the two leftmost columns mark which list each taxon belongs to and whether it clears correction on that list. Each taxon is scored, separately in each of the four reported conditions, by its distance from the fitted prevalence background in units of the local spread ($z$; **Supplementary Text 6E–6F**) — the same quantity whose mean over a list gives the list-level statistic in **Fig. 5E**, so this figure decomposes that number. A taxon is scored in a condition when the background curve can reach it (at least 15 comparable background taxa, and detection in at least three of that condition's study cohorts), and enters the cross-condition test when at least three of the four conditions qualify; 92 of the 104 taxa in the union of the two lists qualify. Significance is a cluster bootstrap over the 34 contributing study cohorts, which resamples whole cohorts rather than treating the four condition values as independent observations, with Benjamini–Hochberg correction within each list. Twenty-four taxa reach $q < 0.20$, twenty above the background and four below, and nine reach $q < 0.05$. Left: list membership, shaded where that list's $q < 0.20$ and starred where $q < 0.05$. Centre: $z$ per taxon and condition — blue where a taxon is less depleted in cases than taxa of the same baseline prevalence, orange where it is more depleted, hatched where no comparable taxa exist to score against; stars give the uncorrected within-condition bootstrap p (*$p < 0.05$, **$p < 0.01$, ***$p < 0.001$). With four conditions per row these cell marks are descriptive rather than the FDR-controlled claim, which is the row. Right: the cross-condition mean $z$ with its 95% bootstrap interval, and directional consistency. The enriched side is led by Escherichia coli (+2.16), *Duodenibacillus* sp900544255 (+1.98), *Fimenecus* sp000432435 (+1.66) and *Prevotella* sp003447235 (+1.66), and includes the polyphenol-converting *Eggerthellaceae Slackia isoflavoniconvertens* and *Ellagibacter isourolithinifaciens*; the depleted side by the butyrate producers *Faecalibacterium prausnitzii_E* (−1.13), *Roseburia* sp900756925 (−1.07) and *Eshraghiella crossota* (−1.02). *Prevotella* appears only on the enriched side here. Just one taxon is unique to each list's significant set, so the $VANISH^{+}$ and $VANISH^{H+}$ lists are almost the same at the taxon level.

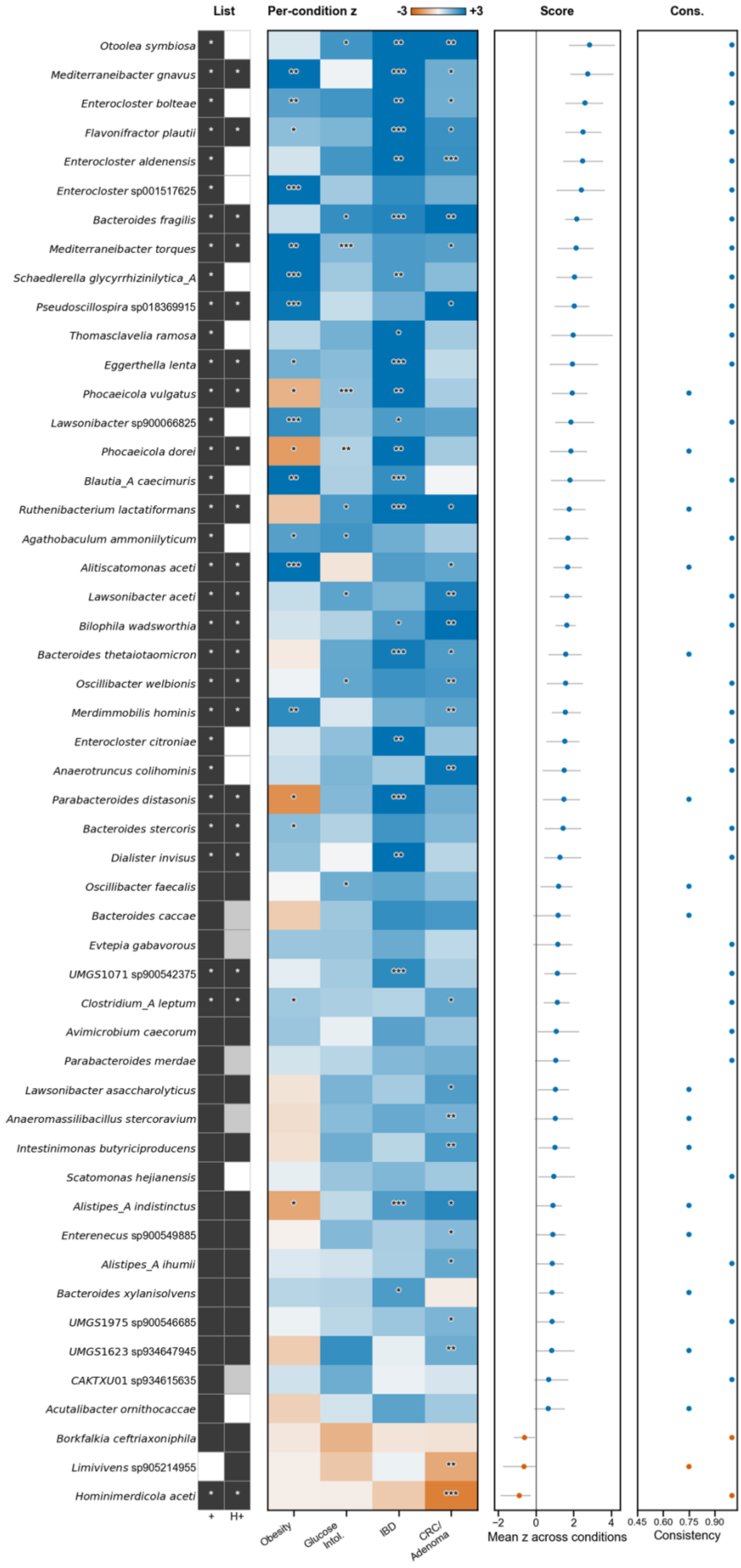

■ on list, q < 0.2 ✱ q < 0.05 ■ on list, n.s. □ not on list ✱ within condition: * p<.05 ** p<.01 *** p<.001 ▨ no comparable taxa

**Figure S28. Taxa carrying the BloSSUM-family health associations.** Layout, scoring and selection are as in **fig. S27**, applied to the BloSSUM$^{+}$ and BloSSUM$^{H+}$ lists, which share 82 of their 100 taxa; all 118 taxa in their union are scored. Fifty-one reach $q < 0.20$, forty-eight above the fitted prevalence background and three below, and thirty-two reach $q < 0.05$. The strongest drivers are *Otoolea symbiosa* (+2.84), *Mediterraneibacter gnavus* (+2.75), *Enterocloster bolteae* (+2.60) and *Flavonifractor plautii* (+2.50); the set runs through *Bacteroides*, *Phocaeicola*, *Parabacteroides*, *Lawsonibacter* and *Oscillibacter* species, with *Hominimerdicola aceti* (−0.89), *Limivivens* sp905214955 (−0.64) and *Borkfalkia ceftriaxoniphila* (−0.61) below the background. Taxa enriched by industrialization therefore move with disease status in one direction at the taxon level as well as in aggregate, matching the list-level result in which BloSSUM$^{H+}$ stands above its background in three of the four tested conditions (**Fig. 5E**). The membership strip carries the comparison between the two lists: of the 18 taxa unique to BloSSUM$^{+}$, 13 clear correction, whereas of the 18 unique to BloSSUM$^{H+}$, one does. Three high-prevalence taxa reverse sign between conditions (*Phocaeicola dorei* and *P. vulgatus*, *Parabacteroides distasonis*: below the background in obesity, far above it in IBD), which the per-condition columns show directly and the cross-condition mean necessarily averages over.

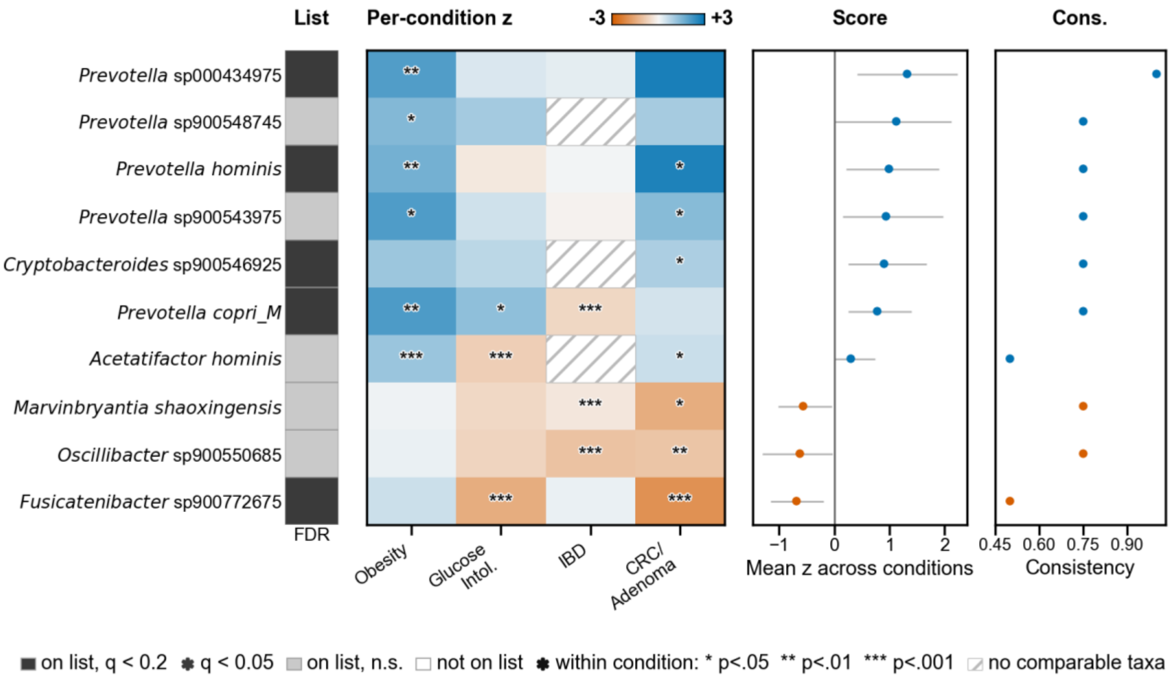

**Figure S29. Taxa contributing most to the original VANISH health association.** Layout, scoring and selection are as in **fig. S27**, applied to the original VANISH list; the single membership column marks which taxa clear correction. Sixty-five of the 124 VANISH taxa can be scored in at least three of the four conditions, 47 lying above the fitted prevalence background and 18 below. This is the lowest proportion of any list, and it is a property of the list rather than of the method: many of its members are too rare for the background curve to reach, or go undetected altogether in a given condition's cohorts. Five taxa reach $q < 0.20$, four above the background and one below, and none reaches $q < 0.05$, so the drivers of this list should be read as candidates rather than as established associations; five further taxa are drawn to meet the figure's ten-row minimum and are not significant. Above the background: *Prevotella* sp000434975 (+1.32), *P. hominis* (+0.99), *Cryptobacteroides* sp900546925 (+0.90) and *P. copri_M* (+0.78). Below it: *Fusicatenibacter* sp900772675 (−0.69). Taxa detected in no sample of either arm are excluded throughout: their case–control shift is zero because nothing was observed rather than because nothing moved, and all of them would otherwise receive the same score at the prevalence floor.

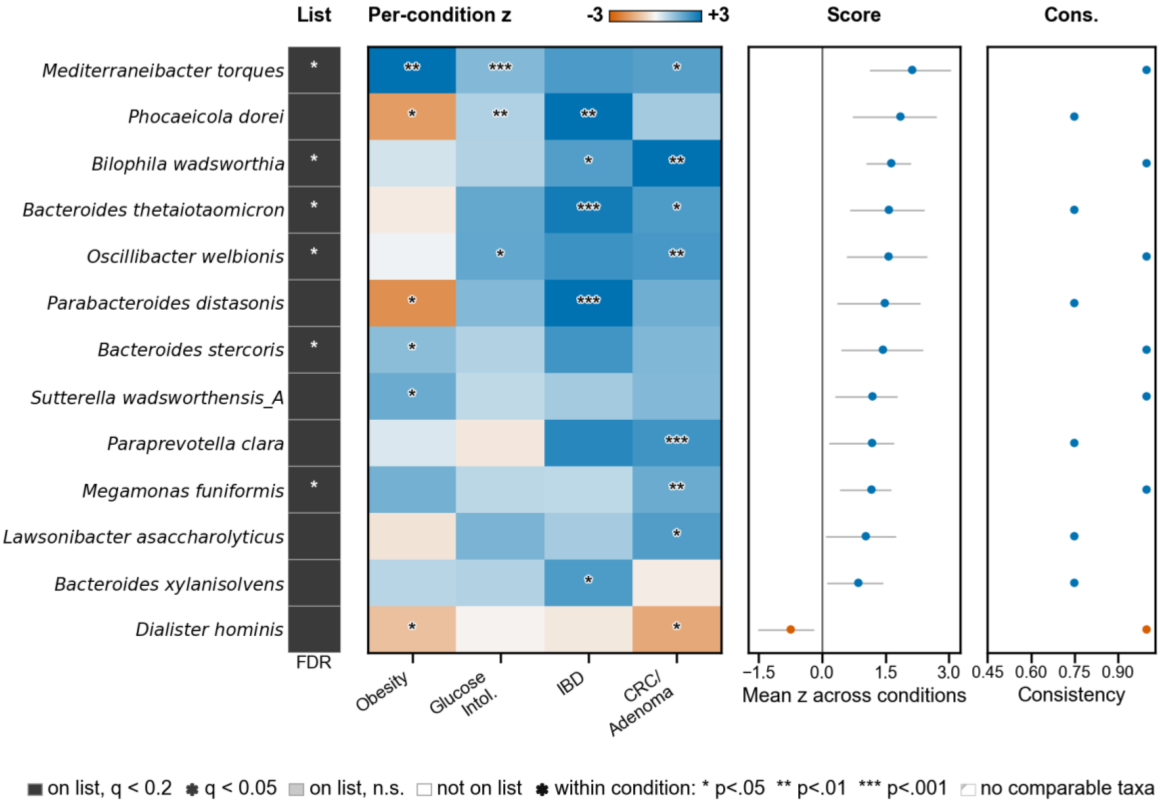

**Figure S30. Taxa contributing most to the original BloSSUM health association.** Layout, scoring and selection are as in **fig. S27**, applied to the original BloSSUM list. All 63 BloSSUM taxa can be scored, 51 lying above the fitted prevalence background and 12 below; 13 reach $q < 0.20$, 12 above the background and one below, and 6 reach $q < 0.05$. The enriched side is led by *Mediterraneibacter torques* (+2.13), *Phocaeicola dorei* (+1.85), *Bilophila wadsworthia* (+1.63) and *Bacteroides thetaiotaomicron* (+1.58), and includes *Parabacteroides distasonis*, *Sutterella wadsworthensis_A*, *Paraprevotella clara* and *Megamonas funiformis*; *Dialister hominis* (−0.74) is the one taxon below the background. Like BloSSUM$^{H+}$ (**fig. S28**), the original BloSSUM list leans strongly toward taxa that are less depleted in cases than their prevalence-matched comparators, so the industrial-enriched signal does not depend on the health-targeted redefinition of the list.

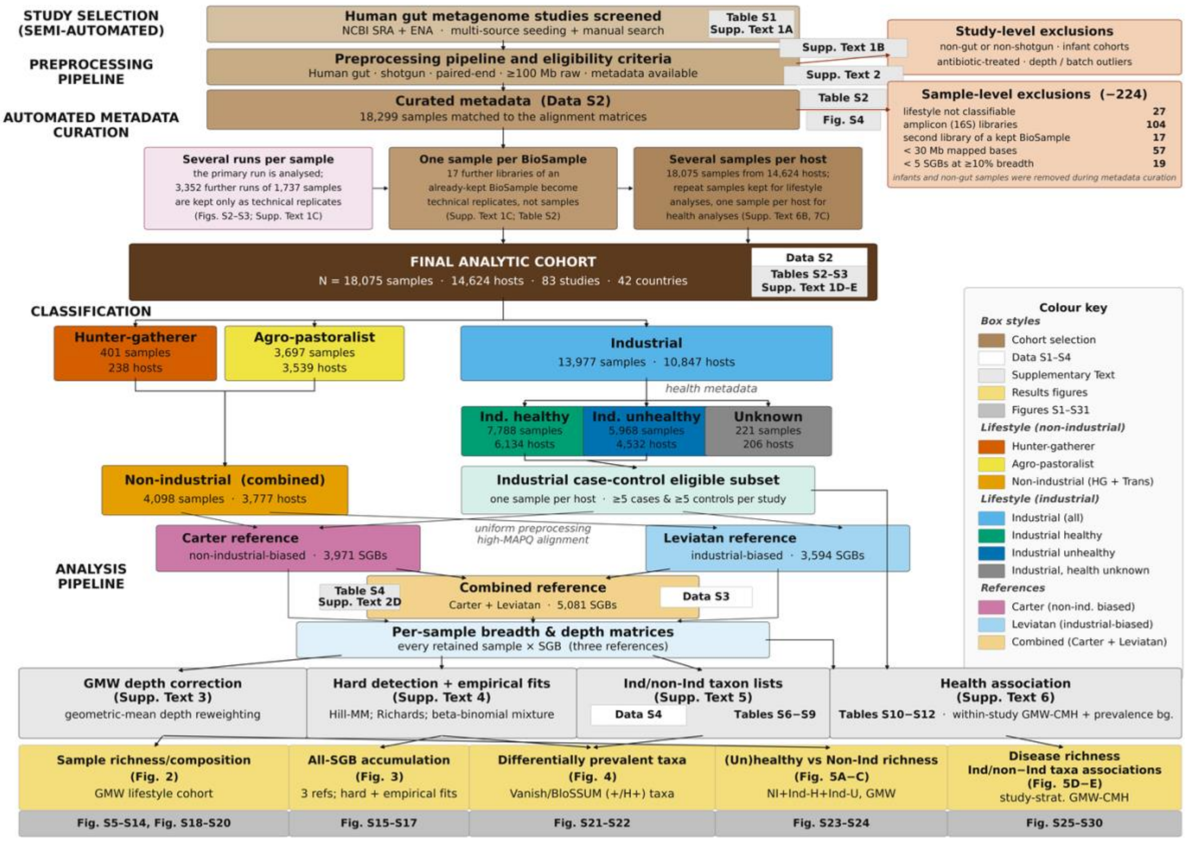

**Figure S31. Roadmap to the study, its supplementary text, tables and figures.** How the analytic cohort was assembled and which analysis each part of it feeds. Reading down: studies screened from the NCBI SRA and ENA, the preprocessing pipeline and its eligibility criteria, and the curated metadata (**Data S2**), with study- and sample-level exclusions broken out on the right. The final analytic cohort is then split by lifestyle (hunter-gatherer, agro-pastoralist, industrial) and, within the industrial arm, by health status; the industrial case–control eligible subset is the one sample per host from studies contributing at least five cases and five controls. Below that, reads from every retained sample are aligned to the Carter and Leviatan catalogues and their merge (**Data S3**), giving the per-sample breadth and depth matrices on which the four analysis blocks draw. Each block names the supplementary text that specifies it, each yellow box the main-text figure it produces, and each grey box the supplementary figures that support it. Badges throughout point to the supplementary text, table or figure that documents a step; "**Supp. Text n**" abbreviates Supplementary Text n. Colour encodes the kind of object, not the result: see the key at right.

#### Supplementary Text 1. Study selection, exclusions, and analytic cohorts

This note defines the cohorts used throughout the study. **Tables S1–S3** summarize study-level metadata, exclusion steps, and the exact cohorts used in each main analysis. To avoid ambiguity, we distinguish three levels of dataset construction: (i) the curated public-domain inventory, (ii) the primary analytic cohort after run consolidation, (iii) the exact-label subsets used for lifestyle and health analyses. Unless otherwise stated, counts reported below refer to these analysis-ready cohorts rather than the larger run-level curation inventory.

**Table S1. High-level summary of the studies included in this meta-analysis.**

| Author | Year | Country | N | Age median [IQR] | Age N | Gbp/sample median [IQR] | Lifestyle | Unhealthy% | Conditions | Reference |
|---|---|---|---|---|---|---|---|---|---|---|
| **AsnicarF** | 2017 | Italy | 8 | | 0 | 1.00 [0.81–1.48] | Industrial | 0.0 | Healthy controls only | (*16*) |
| **AsnicarF** | 2021 | United Kingdom | 1,000 | 46.0 [37.0–54.0] | 1,000 | 3.58 [3.21–4.15] | Industrial | 14.4 | Obesity, Overweight | (*17*) |
| **AsnicarF** | 2021 | United States of America | 97 | 42.0 [31.0–54.0] | 97 | 3.63 [3.30–4.18] | Industrial | 16.5 | Obesity, Overweight | (*17*) |
| **BackhedF** | 2015 | Sweden | 81 | | 0 | 1.96 [1.78–2.30] | Industrial | 0.0 | Healthy controls only | (*18*) |
| **BedarfJR** | 2017 | Germany | 37 | | 0 | 2.49 [1.44–3.07] | Industrial | 56.8 | Parkinson's disease, Obesity, Overweight | (*19*) |
| **BeghiniF** | 2023 | Honduras | 1,887 | 38.0 [27.2–53.0] | 1,782 | 4.18 [3.44–4.99] | Agro-pastoralist | 15.3 | Obesity, Overweight | (*20*) |
| **BrambleMS** | 2021 | DRC Congo | 180 | 13.0 [13.0–13.8] | 6 | 0.56 [0.47–0.65] | Agro-pastoralist | 0.0 | Healthy controls only | (*21*) |
| **BritoIL** | 2016 | Fiji | 222 | 33.0 [17.0–45.0] | 157 | 3.50 [2.78–4.05] | Agro-pastoralist | 0.0 | Healthy controls only | (*22*) |
| **BrownCT** | 2011 | Finland | 8 | | 0 | 1.17 [0.95–1.43] | Industrial | 50.0 | Type 1 diabetes | (*23*) |
| **CarterM** | 2023 | Nepal | 56 | | 0 | 9.89 [8.43–12.27] | Agro-pastoralist | 0.0 | Healthy controls only | (*7*) |
| **CarterM** | 2023 | Tanzania | 300 | 35.0 [9.0–47.0] | 69 | 7.48 [5.80–11.52] | Hunter-gatherer | 0.7 | Obesity, Overweight | (*7*) |
| **CarterM** | 2023 | United States of America | 12 | | 0 | 15.12 [13.72–16.65] | Industrial | 0.0 | Healthy controls only | (*7*) |
| **Chengping W** | 2017 | China | 201 | 39.0 [32.0–45.0] | 201 | 1.58 [1.06–2.04] | Industrial | 44.3 | Ankylosing spondylitis, Overweight | (*24*) |
| **ContevilleLC** | 2015 | Brazil | 6 | 55.0 [43.8–55.0] | 6 | 0.10 [0.07–0.32] | Hunter-gatherer | 0.0 | Healthy controls only | (*25*) |
| **CosteaPI** | 2017 | Germany | 93 | 36.0 [7.0–40.0] | 53 | 2.49 [1.81–2.84] | Industrial | 0.0 | Healthy controls only | (*26*) |
| **CosteaPI** | 2017 | Kazakhstan | 171 | 49.5 [42.0–58.0] | 80 | 1.12 [0.65–2.42] | Agro-pastoralist | 25.1 | Acute diarrhoea, Obesity, Overweight | (*26*) |
| **DavidLA** | 2015 | Bangladesh | 25 | | 0 | 0.17 [0.11–0.36] | Industrial | 84.0 | Acute diarrhoea | (*27*) |
| **Eloe-FadroshEA** | 2015 | United States of America | 9 | | 0 | 5.64 [4.44–6.06] | Industrial | 0.0 | Healthy controls only | (*28*) |
| **FengQ** | 2015 | Austria | 156 | 68.0 [63.0–72.0] | 156 | 2.31 [1.96–2.55] | Industrial | 85.3 | Type 2 diabetes, MASLD, Hypertension, Colorectal cancer, Colorectal adenoma, Obesity, Overweight | (*29*) |

| | | | | | | | | | | |
|---|---|---|---|---|---|---|---|---|---|---|
| **FrankelAE** | 2017 | United States of America | 39 | 68.0 [57.5–75.5] | 39 | 4.12 [2.66–4.74] | Industrial | 100.0 | Inflammatory bowel disease | (*30*) |
| **HMP** | 2012 | United States of America | 435 | 26.0 [23.0–28.0] | 147 | 2.26 [0.86–3.21] | Industrial | 0.0 | C. difficile infection, Obesity | (*31*) |
| **HallAB** | 2017 | United States of America | 247 | | 0 | 1.29 [0.80–1.85] | Industrial | 71.3 | Inflammatory bowel disease | (*32*) |
| **HanniganGD** | 2017 | Canada | 26 | 64.5 [58.0–68.8] | 26 | 0.22 [0.15–0.38] | Industrial | 88.5 | Colorectal cancer, Colorectal adenoma, Obesity, Overweight | (*33*) |
| **HanniganGD** | 2017 | United States of America | 52 | 55.5 [50.5–61.0] | 52 | 0.27 [0.22–0.41] | Industrial | 67.3 | Colorectal cancer, Colorectal adenoma, Obesity, Overweight | (*33*) |
| **HansenLBS** | 2018 | Denmark | 206 | 51.0 [45.0–57.0] | 206 | 3.09 [2.49–3.92] | Industrial | 34.0 | Obesity, Overweight | (*34*) |
| **HeQ** | 2017 | China | 108 | 23.0 [17.0–29.0] | 98 | 2.71 [2.39–3.11] | Industrial | 50.0 | Crohn's disease, Coeliac disease, Obesity, Overweight | (*35*) |
| **IaniroG** | 2022 | Italy | 110 | | 0 | 3.01 [1.73–4.35] | Industrial | 90.9 | Inflammatory bowel disease, C. difficile infection | (*36*) |
| **IjazUZ** | 2017 | United Kingdom | 92 | 13.0 [11.0–35.0] | 91 | 0.54 [0.37–0.73] | Industrial | 58.7 | Inflammatory bowel disease | (*37*) |
| **JacobsonDK** | 2021 | Burkina Faso | 90 | 34.5 [20.0–52.8] | 90 | 2.30 [1.69–3.44] | Agro-pastoralist | 0.0 | Atherosclerotic CVD | (*38*) |
| **JieZ** | 2017 | China | 385 | | 0 | 2.13 [1.60–2.74] | Industrial | 58.4 | Impaired glucose tolerance, Atherosclerotic CVD, Obesity, Overweight | (*39*) |
| **KarlssonFH** | 2013 | Sweden | 145 | 70.4 [70.0–71.0] | 145 | 1.18 [0.97–1.64] | Industrial | 54.5 | Type 2 diabetes, Impaired glucose tolerance, Obesity, Overweight | (*40*) |
| **KimJ** | 2022 | South Korea | 58 | | 0 | 2.62 [2.26–3.16] | Industrial | 0.0 | Healthy controls only | (*41*) |
| **KliminaKM** | 2019 | Russia | 5 | 31.0 [22.5–38.2] | 4 | 0.81 [0.81–1.43] | Industrial | 20.0 | Type 2 diabetes | (*42*) |
| **KuangYS** | 2017 | China | 36 | 29.0 [26.0–30.2] | 36 | 2.92 [2.54–3.17] | Industrial | 0.0 | Overweight | (*43*) |
| **LeChatelierE** | 2013 | Denmark | 129 | 56.0 [51.0–62.0] | 129 | 3.51 [2.89–4.09] | Industrial | 62.0 | Melanoma, Obesity, Overweight | (*44*) |
| **LeeKA** | 2022 | Netherlands | 55 | | 0 | 1.47 [0.87–2.28] | Industrial | 100.0 | Melanoma, Obesity, Overweight | (*45*) |
| **LeeKA** | 2022 | Spain | 12 | | 0 | 3.68 [3.06–5.60] | Industrial | 100.0 | Melanoma, Obesity, Overweight | (*45*) |
| **LeeKA** | 2022 | United Kingdom | 97 | | 0 | 3.10 [1.96–4.80] | Industrial | 100.0 | Melanoma, Obesity, Overweight | (*45*) |
| **LiJ** | 2014 | Denmark | 109 | 60.0 [51.0–65.0] | 109 | 2.69 [2.40–2.96] | Industrial | 78.0 | Type 2 diabetes, Obesity, Overweight | (*46*) |
| **LiJ** | 2014 | Spain | 138 | 41.0 [30.0–50.0] | 137 | 3.01 [2.44–3.52] | Industrial | 12.3 | Obesity, Overweight | (*46*) |
| **LiJ** | 2017 | China | 196 | | 0 | 2.02 [1.53–2.44] | Industrial | 79.1 | Hypertension | (*47*) |
| **LiSS** | 2016 | Netherlands | 55 | | 0 | 2.08 [1.63–2.52] | Industrial | 90.9 | Metabolic syndrome, Hypertension | (*48*) |
| **LiuR** | 2017 | China | 217 | | 0 | 2.50 [2.08–2.73] | Industrial | 51.6 | Obesity, Overweight | (*49*) |
| **LiuW** | 2016 | China | 47 | | 0 | 3.08 [2.14–3.71] | Agro-pastoralist | 0.0 | Healthy controls only | (*50*) |

| | | | | | | | | | | |
|---|---|---|---|---|---|---|---|---|---|---|
| **LiuW** | 2016 | Mongolia | 63 | | 0 | 2.68 [2.11–3.47] | Agro-pastoralist | 0.0 | Healthy controls only | (*50*) |
| **Lloyd-PriceJL** | 2019 | United States of America | 1125 | 21.0 [14.0–41.0] | 1120 | 1.01 [0.75–1.29] | Industrial | 70.4 | Crohn's disease, Ulcerative colitis | (*51*) |
| **LokmerA** | 2019 | Cameroon | 16 | 49.0 [32.5–59.5] | 16 | 1.87 [1.50–2.07] | Hunter-gatherer | 0.0 | Healthy controls only | (*52*) |
| **LokmerA** | 2019 | Cameroon | 41 | 53.0 [44.0–62.0] | 41 | 1.79 [1.58–1.97] | Agro-pastoralist | 7.3 | MASLD, Obesity, Overweight | (*52*) |
| **LomanNJ** | 2013 | Germany | 35 | | 0 | 0.52 [0.36–0.72] | Industrial | 100.0 | MASLD | (*53*) |
| **LoombaR** | 2017 | United States of America | 86 | | 0 | 1.37 [1.02–1.82] | Industrial | 100.0 | MASLD, Liver cirrhosis | (*54*) |
| **LouisS** | 2016 | Germany | 92 | | 0 | 0.97 [0.78–1.17] | Industrial | 16.3 | Obesity, Overweight | (*55*) |
| **Maldonaldo-GomezMX** | 2016 | United States of America | 64 | 27.0 [24.0–30.0] | 64 | 1.23 [0.85–1.57] | Industrial | 0.0 | Melanoma, Overweight | (*56*) |
| **MatsonV** | 2018 | United States of America | 39 | | 0 | 4.12 [3.62–6.39] | Industrial | 100.0 | Unspecified illness | (*57*) |
| **McDonaldD** | 2018 | Canada | 1 | 5.0 [5.0–5.0] | 1 | 0.07 [0.07–0.07] | Industrial | 0.0 | Healthy controls only | (*58*) |
| **McDonaldD** | 2018 | Philippines | 58 | 40.5 [26.2–49.2] | 58 | 0.07 [0.05–0.08] | Industrial | 15.5 | Type 2 diabetes, Obesity, Overweight | (*58*) |
| **McDonaldD** | 2018 | United Kingdom | 3 | 29.0 [24.5–44.5] | 3 | 0.12 [0.09–0.18] | Industrial | 33.3 | Crohn's disease | (*58*) |
| **McDonaldD** | 2018 | United States of America | 77 | 39.0 [18.5–51.2] | 76 | 0.09 [0.06–0.14] | Industrial | 18.2 | Type 2 diabetes, Crohn's disease, Ulcerative colitis, Obesity, Overweight | (*58*) |
| **MehtaRS** | 2018 | United States of America | 803 | | 0 | 0.83 [0.62–1.04] | Industrial | 0.0 | Healthy controls only | (*59*) |
| **Nagy-SzakalD** | 2017 | United States of America | 100 | 51.0 [51.0–51.0] | 100 | 2.55 [1.71–3.67] | Industrial | 58.0 | Obesity, Overweight | (*60*) |
| **NielsenHB** | 2014 | Denmark | 177 | 55.0 [49.0–59.8] | 82 | 1.09 [0.86–1.61] | Industrial | 55.9 | Inflammatory bowel disease, Obesity, Overweight | (*61*) |
| **NielsenHB** | 2014 | Spain | 214 | 41.0 [32.0–49.8] | 202 | 1.78 [1.23–2.03] | Industrial | 43.5 | Inflammatory bowel disease, Obesity, Overweight | (*61*) |
| **Obregon-TitoAJ** | 2015 | Peru | 20 | 15.0 [4.8–30.0] | 20 | 2.16 [1.90–2.38] | Hunter-gatherer | 0.0 | Overweight | (*62*) |
| **Obregon-TitoAJ** | 2015 | Peru | 11 | 30.5 [14.8–36.0] | 10 | 0.59 [0.45–0.67] | Agro-pastoralist | 9.1 | Obesity, Overweight | (*62*) |
| **Obregon-TitoAJ** | 2015 | United States of America | 22 | 27.5 [26.0–34.8] | 22 | 2.17 [1.96–2.35] | Industrial | 9.1 | Obesity, Overweight | (*62*) |
| **PasolliE** | 2019 | Madagascar | 111 | 33.0 [26.0–44.0] | 111 | 1.57 [0.80–2.52] | Agro-pastoralist | 0.0 | Overweight | (*5*) |
| **PehrssonEC** | 2016 | El Salvador | 106 | 29.0 [11.0–55.0] | 106 | 0.32 [0.16–0.43] | Agro-pastoralist | 0.0 | Melanoma | (*63*) |
| **PehrssonEC** | 2016 | Peru | 66 | 28.5 [13.0–43.2] | 66 | 0.44 [0.27–0.52] | Industrial | 0.0 | Healthy controls only | (*63*) |
| **PetersBA** | 2019 | United States of America | 27 | 72.0 [62.5–80.0] | 27 | 2.22 [1.66–3.83] | Industrial | 100.0 | Obesity, Overweight | (*64*) |

| | | | | | | | | | | |
|---|---|---|---|---|---|---|---|---|---|---|
| **PetersenLM** | 2017 | United States of America | 89 | | 0 | 0.18 [0.14–0.53] | Industrial | 0.0 | Healthy controls only | (*65*) |
| **QinJ** | 2012 | China | 370 | 48.0 [39.0–58.0] | 370 | 1.54 [1.09–1.97] | Industrial | 50.0 | Type 2 diabetes, Obesity, Overweight | (*66*) |
| **QinN** | 2014 | China | 237 | 45.0 [38.0–54.0] | 237 | 1.32 [0.89–2.05] | Industrial | 51.9 | Liver cirrhosis, Hepatitis, Obesity, Overweight | (*67*) |
| **RaethongN** | 2021 | Thailand | 10 | | 0 | 6.43 [5.78–7.02] | Industrial | 0.0 | Healthy controls only | (*68*) |
| **RampelliS** | 2015 | Italy | 11 | 30.0 [28.0–34.0] | 11 | 0.21 [0.17–0.65] | Industrial | 0.0 | Healthy controls only | (*69*) |
| **RampelliS** | 2015 | Tanzania | 27 | 30.0 [21.5–40.0] | 27 | 0.62 [0.37–1.06] | Hunter-gatherer | 0.0 | Healthy controls only | (*69*) |
| **RaymondF** | 2016 | Canada | 72 | 24.0 [22.8–26.8] | 72 | 5.87 [4.58–8.22] | Industrial | 0.0 | Overweight | (*70*) |
| **RettedalAE** | 2021 | New Zealand | 79 | 63.0 [60.0–67.0] | 79 | 1.26 [1.02–1.38] | Industrial | 70.9 | Obesity, Overweight | (*71*) |
| **RosaBA** | 2018 | Indonesia | 20 | 21.4 [14.7–30.9] | 8 | 5.94 [5.32–6.81] | Agro-pastoralist | 100.0 | Soil-transmitted helminth infection | (*72*) |
| **RosaBA** | 2018 | Liberia | 4 | | 0 | 2.80 [2.25–3.17] | Agro-pastoralist | 100.0 | Soil-transmitted helminth infection | (*72*) |
| **RubelMA** | 2020 | Cameroon | 32 | 32.0 [26.8–39.5] | 32 | 0.78 [0.63–0.90] | Hunter-gatherer | 25.0 | Soil-transmitted helminth infection, Obesity, Overweight | (*73*) |
| **RubelMA** | 2020 | Cameroon | 71 | 45.0 [32.0–53.5] | 71 | 0.83 [0.68–1.01] | Agro-pastoralist | 16.9 | Soil-transmitted helminth infection, Obesity, Overweight | (*73*) |
| **SankaranarayananK** | 2015 | United States of America | 12 | 46.0 [29.8–57.5] | 12 | 1.94 [1.62–2.09] | Industrial | 83.3 | Type 2 diabetes, Obesity, Overweight | (*74*) |
| **SchirmerM** | 2016 | Netherlands | 471 | 23.0 [21.0–27.0] | 465 | 1.11 [0.94–1.32] | Industrial | 1.7 | Obesity, Overweight | (*75*) |
| **TamburiniFB** | 2022 | South Africa | 51 | | 0 | 0.39 [0.34–0.48] | Industrial | 0.0 | Healthy controls only | (*76*) |
| **TamburiniFB** | 2022 | South Africa | 117 | | 0 | 0.92 [0.63–1.13] | Agro-pastoralist | 0.0 | Healthy controls only | (*76*) |
| **TettAJ** | 2019 | Ethiopia | 27 | 26.5 [25.0–29.5] | 26 | 2.24 [1.74–2.44] | Agro-pastoralist | 0.0 | Healthy controls only | (*77*) |
| **TettAJ** | 2019 | Ghana | 42 | 24.0 [7.0–34.0] | 41 | 1.50 [1.39–2.06] | Agro-pastoralist | 0.0 | Healthy controls only | (*77*) |
| **TettAJ** | 2019 | Tanzania | 66 | 20.0 [9.0–28.0] | 66 | 1.94 [1.54–2.64] | Agro-pastoralist | 0.0 | Healthy controls only | (*77*) |
| **ThomasAM** | 2019 | Italy | 140 | 67.0 [60.8–75.2] | 80 | 1.77 [1.40–2.54] | Industrial | 65.0 | Colorectal cancer, Colorectal adenoma, Obesity, Overweight | (*78*) |
| **Valles-ColomerM** | 2023 | Argentina | 13 | 27.0 [21.0–31.0] | 13 | 4.14 [2.54–5.58] | Agro-pastoralist | 0.0 | Healthy controls only | (*79*) |
| **Valles-ColomerM** | 2023 | China | 8 | | 0 | 4.04 [3.65–7.16] | Agro-pastoralist | 0.0 | Healthy controls only | (*79*) |
| **Valles-ColomerM** | 2023 | Colombia | 12 | 21.5 [18.8–32.0] | 12 | 1.55 [0.82–2.70] | Agro-pastoralist | 0.0 | Healthy controls only | (*79*) |
| **Valles-ColomerM** | 2023 | Guinea-Bissau | 319 | 19.0 [9.0–39.0] | 306 | 1.99 [1.50–2.76] | Agro-pastoralist | 0.0 | Healthy controls only | (*79*) |
| **Valles-ColomerM** | 2023 | Italy | 169 | 25.3 [6.4–36.8] | 169 | 2.80 [1.57–4.51] | Industrial | 0.0 | Healthy controls only | (*79*) |
| **VangayP** | 2018 | Thailand | 13 | 39.0 [32.0–50.0] | 13 | 1.77 [1.57–1.85] | Agro-pastoralist | 7.7 | Obesity, Overweight | (*80*) |

| | | | | | | | | | | |
|---|---|---|---|---|---|---|---|---|---|---|
| **VangayP** | 2018 | United States of America | 2 | | 0 | 1.41 [1.39–1.42] | Industrial | 0.0 | Healthy controls only | (*80*) |
| **VogtmannE** | 2016 | United States of America | 61 | 65.0 [55.8–71.0] | 58 | 2.75 [2.25–3.35] | Industrial | 59.0 | Colorectal cancer, Obesity, Overweight | (*81*) |
| **WallenZD** | 2022 | United States of America | 369 | 68.0 [62.0–73.0] | 369 | 2.83 [2.29–3.73] | Industrial | 73.4 | Ulcerative colitis, Parkinson's disease, Obesity, Overweight | (*82*) |
| **WirbelJ** | 2018 | Germany | 82 | 60.0 [53.0–65.8] | 82 | 2.00 [1.69–2.75] | Industrial | 34.1 | Type 2 diabetes, Colorectal cancer, Obesity, Overweight | (*83*) |
| **XieH** | 2016 | United Kingdom | 250 | 62.0 [56.0–67.0] | 250 | 3.67 [3.33–4.03] | Industrial | 19.6 | Type 2 diabetes, Obesity, Overweight | (*84*) |
| **XiongR** | 2023 | United States of America | 214 | 42.5 [31.0–56.0] | 214 | 1.04 [0.73–1.85] | Industrial | 57.0 | Myalgic encephalomyelitis | (*85*) |
| **YachidaS** | 2019 | Japan | 640 | 64.0 [55.0–70.0] | 612 | 2.76 [2.20–3.41] | Industrial | 57.3 | Colorectal cancer, Obesity, Overweight | (*86*) |
| **YassourM** | 2016 | South Korea | 36 | | 0 | 1.20 [1.07–1.29] | Industrial | 0.0 | Healthy controls only | (*87*) |
| **YassourM** | 2018 | Finland | 116 | | 0 | 1.61 [1.42–1.84] | Industrial | 0.0 | Healthy controls only | (*88*) |
| **YeZ** | 2018 | China | 66 | 37.0 [28.5–46.0] | 66 | 2.21 [1.51–2.91] | Industrial | 33.3 | Obesity, Overweight | (*89*) |
| **YeZ** | 2020 | China | 55 | 38.0 [31.5–48.5] | 55 | 2.24 [1.52–2.93] | Industrial | 3.6 | Obesity, Overweight | (*90*) |
| **YuJ** | 2017 | China | 128 | 64.0 [58.8–70.0] | 128 | 2.44 [2.06–2.78] | Industrial | 71.9 | Type 2 diabetes, Colorectal cancer, Obesity, Overweight | (*91*) |
| **ZeeviD** | 2015 | Israel | 882 | | 0 | 1.31 [1.04–1.67] | Industrial | 5.7 | Obesity, Overweight | (*92*) |
| **ZellerG** | 2014 | France | 127 | 63.0 [56.0–68.5] | 127 | 2.16 [1.53–2.88] | Industrial | 68.5 | Colorectal cancer, Colorectal adenoma, Obesity, Overweight | (*93*) |
| **ZellerG** | 2014 | Germany | 39 | 59.0 [46.0–70.5] | 39 | 2.52 [2.23–3.27] | Industrial | 89.7 | Type 2 diabetes, Colorectal cancer, Obesity, Overweight | (*93*) |
| **ZhangX** | 2015 | China | 228 | 46.0 [39.0–54.0] | 228 | 2.93 [2.54–3.26] | Industrial | 59.6 | Rheumatoid arthritis, Schizophrenia, Obesity, Overweight | (*94*) |
| **ZhuF** | 2020 | China | 171 | 27.0 [22.0–37.0] | 171 | 4.42 [3.15–5.35] | Industrial | 52.6 | Schizophrenia, Overweight | (*95*) |
| **iMSMS** | 2022 | Argentina | 320 | 40.0 [32.0–53.5] | 320 | 0.10 [0.07–0.13] | Industrial | 55.3 | Multiple sclerosis, Obesity, Overweight | (*96*) |
| **iMSMS** | 2022 | Spain | 1 | 47.0 [47.0–47.0] | 1 | 0.16 [0.16–0.16] | Industrial | 100.0 | Multiple sclerosis | (*96*) |
| **iMSMS** | 2022 | United Kingdom | 262 | 55.0 [48.0–61.0] | 262 | 0.08 [0.06–0.13] | Industrial | 55.3 | Multiple sclerosis, Obesity, Overweight | (*96*) |
| **iMSMS** | 2022 | United States of America | 549 | 55.0 [44.0–62.0] | 549 | 0.08 [0.06–0.13] | Industrial | 60.5 | Multiple sclerosis, Obesity, Overweight | (*96*) |

#### 1A. Study identification and metadata curation

We sought to assemble a large, reproducible, public-domain dataset of human gut metagenomes suitable for testing how lifestyle structure, sequencing depth, and population sampling affect estimates of microbial richness, population-level persistence, and health association. We therefore restricted the study to publicly accessible metagenomic datasets deposited in the NCBI Sequence Read Archive (SRA) or European Nucleotide Archive (ENA), so that both the curated metadata and the raw sequencing data could be independently reanalyzed in future work.

Study identification was seeded from existing curated resources and then extended by manual review. In particular, we drew on the curated non-industrial metadata assembled by Carter et al., curated health-associated industrial datasets compiled by Silva et al. (*97*), MicrobiomeDB (*98*), recent population-scale metagenomic datasets such as Beghini et al. (*20*), and additional studies identified through the Valles-Colomer datasets (*79*) and manual literature searches. Our goal was not to exhaustively capture every published human gut metagenome, but rather to assemble the broadest possible set of deeply sequenced public-domain gut metagenomes with sufficiently rich metadata for lifestyle and health-stratified analyses. This emphasis was deliberate: for the questions addressed here, deeply sequenced samples with usable metadata are substantially more informative than a larger number of shallow or poorly annotated datasets.

**1B. Eligibility criteria and sample-level quality control**

Studies were eligible if they profiled the human gut microbiome using shotgun metagenomic sequencing and provided enough information to support downstream lifestyle and health annotation. We prioritized paired-end sequencing and required each retained sample to meet both of the following depth thresholds: at least 100 million raw bases prior to preprocessing and at least 30 million bacterial bases mapped at MAPQ >= 30 after host/PhiX removal and alignment. We selected on bases rather than reads because read length varies systematically across studies, with many older industrial datasets sequenced with 100 bp length reads and many non-industrial discovery-oriented datasets sequenced with 150 bp length reads; selecting on read count alone would therefore introduce an additional technical bias.

Only gut-derived metagenomes were retained. Non-gut samples, including wastewater, oral, skin, milk, and other non-fecal microbiome datasets, were excluded during study-level review. We also excluded samples lacking usable metadata, samples failing preprocessing or alignment quality checks, and samples from children <3 years old. This age filter was imposed because neonate and infant gut microbiomes are both biologically distinct and disproportionately represented in industrial cohorts; including them would make lifestyle comparisons of adult-like gut richness artificially more extreme and biologically harder to interpret.

**1C. Run consolidation, longitudinal sampling, and technical replicate handling**

Several additional harmonization steps were required before the curated inventory could be converted into a stable analysis cohort.

First, when the same biosample was represented by multiple sequencing runs, we consolidated runs at the biosample level by concatenating their reads across runs. In cases where a biosample was represented by both paired-end and single-end runs, the paired-end run was preferred. Repeated samples from the same individual (distinct biosamples) were retained as separate samples in the lifestyle analyses. For accumulation and pooled detection this is conservative: serial samples from one person resemble each other more than samples from different people, so they contribute less new diversity than an equal number of independent hosts and can at worst slow the accumulation of newly detected taxa. Additional sample size is crucial to find specific evidence for rare taxa that are right at the threshold of detectability and may vary temporally within an individual For per-sample richness comparisons they are not independent observations; their effect is reported in **Supplementary Text 7C**. In the health analyses, repeat samples from one person would inflate the apparent size of a case–control comparison, so each comparison retains one sample per host (**Supplementary Text 6B**). The 18,075 samples represent 14,624 hosts (**Table S2**); the largest repeat-sampling studies are Lloyd-Price et al. 2019 (*51*) (1,125 samples from 104 hosts), iMSMS 2022 (*96*) (1,132 from 611) and Mehta et al. 2018 (*59*) (803 from 301). Samples without an archived host identifier (5,042) are counted as separate hosts, so 14,624 is an upper bound.

Second, older studies sometimes deposited multiple accessions corresponding to what were nominally technical replicates of the same biosample across different sequencing runs. These were not automatically treated as independent samples. Instead, one run per biosample was chosen as the primary run for the primary analytic cohort, while all other accession-level provenance was retained in the metadata for reproducibility. Where a biosample carried more than one library across different SRS/ERS accessions, the deepest library was kept. This arose once: Carter et al. 2023 (*7*) registered 16 Hadza stools with both a MoBio PowerSoil and a phenol-chloroform library, and re-sequenced one Nepali stool. For the paired stools we kept the phenol-chloroform library, which was the more deeply sequenced in all 16 (median 2.05× mapped bases).

Third, we explicitly screened secondary runs for non-redundancy. In total, 3,352 secondary runs linked to 1,737 primary samples were evaluated and were used as technical replicates for the primaries.

Technical-replicate reproducibility robustness checks are shown in **Figs. S2–3**.

##### 1D. Canonical cohort definitions

The broadest post-curation inventory contained 18,303 qualified public-domain metagenomic runs spanning 83 study datasets and 42 countries. These counts refer to the curated run-level inventory after metadata screening and sample-level eligibility filtering, not yet to the final analysis-ready sample cohorts.

After study-level manual review, run consolidation, and restriction to one primary run per biosample, as well as subsequent requirements for >100 million raw bases and >30 million bases at MAPQ≥30, and a minimum of 5 SGBs found per sample, the 18,303-run curated inventory was reduced by 228 runs to the final analytic cohort: 18,075 samples from 14,624 hosts, from 83 study datasets across 42 countries (**Figure 1A**; the stepwise reduction is given in **Table S2**).

This subset contained 401 hunter-gatherer samples (238 hosts), 3,697 agro-pastoralist samples (3,539 hosts), and 13,977 industrial samples (10,847 hosts). For two-group analyses, hunter-gatherer and agro-pastoralist samples were combined into a single non-industrial group ($n$ = 4,098). The remaining 27 samples had unclear or mixed lifestyle assignments; they were excluded from further analyses.

Health metadata were then layered onto the industrial subset. Among the 13,977 industrial samples, 7,788 were classified as industrial healthy, 5,968 as industrial unhealthy, and 221 as industrial, health status unknown. These labels defined the broad health-stratified cohorts used in **Fig. 5A-C**. The stricter industrial case-control eligible subset used for the study-stratified health-association framework was defined downstream by requiring analyzable control/disease strata within study; exact per-condition counts for those analyses are given in **Table S3**.

The stepwise reduction from curated run inventory to analysis-ready cohorts is summarized in **Table S2** and schematized in **Fig. 1A** in the main text. **Table S2** also carries a chimpanzee column. Every retained sample was additionally aligned against a chimpanzee-derived SGB catalogue, which is used only as a negative control for the pooled-detection analyses (**Supplementary Text 4K**) and contributes to none of the human results.

**1E. Figure-specific cohorts**

The purpose of the cohort hierarchy above is to ensure that each main-text analysis can be traced to an explicitly defined sample set.

Two names recur below. The lifestyle-classified subset is the final analytic cohort of 18,075 samples, each carrying an unambiguous hunter-gatherer, agro-pastoralist or industrial label. The industrial case–control eligible subset is the part of the industrial arm that enters the health analyses: one sample per host, drawn only from studies contributing at least five cases and five controls.

**Fig. 1** uses the lifestyle-classified subset ($n$ = 18,075) and provides a global overview of the curated dataset and its geographic distribution.

**Fig. 2** uses the lifestyle-classified subset **(**$n$ = 18,075**)** to compare per-sample richness and composition across hunter-gatherer, agro-pastoralist, and industrial lifestyles after correcting for depth imbalance.

**Fig. 3** also uses the lifestyle-classified subset **(**$n$ = 18,075**),** pooling samples within lifestyle groups to estimate population-level accumulation and detectability under both reference catalogues.

**Fig. 4** likewise draws from the lifestyle-classified subset **(**$n$ = 18,075**),** with the primary two-group lifestyle comparison defined as industrial ($n$ = 13,977) versus non-industrial ($n$ = 4,098)**.**

**Fig. 5A–C** uses samples with exact lifestyle labels and usable broad health assignments, namely non-industrial ($n$ = 4,098)**,** industrial healthy ($n$ = 7,788), and industrial unhealthy ($n$ = 5,968), for a total of 17,854 samples. Industrial samples with unknown health status **(**$n$ = 221**)** are excluded from these panels.

**Fig. 5D–H** uses only the industrial case-control eligible subset**,** defined at the study-stratum level for each disease analysis. Because these counts vary by condition and by the availability of analyzable healthy and disease groups within study, they are reported explicitly in **Table S3** rather than as a single global *N*. Each comparison retains one sample per host.

Exact cohort definitions for **Fig. 1–5** are summarized in **Table S3**.

#### Table S2. Cohort derivation and exclusion cascade

| Step | Carter | Leviatan | Combined | Chimp | Notes |
|---|---|---|---|---|---|
| **External curation** | | | | | |
| Curated metadata (M1) matched to depth/breadth matrix | 18,299 | 18,299 | 18,299 | 4,431 | M1 rows with a profile in both matrices. |
| **Pipeline derivation** | | | | | |
| Duplicate IDs resolved | 17 | 17 | 17 | 0 | Two libraries for the same BioSample |
| Lifestyle not Industrial/Non-Industrial | 27 | 27 | 27 | 27 | Unclear or mixed lifestyle. |
| Amplicon exclusion | 104 | 104 | 104 | 90 | 16S libraries registered as WGS: mapped bases accumulate in a small conserved window. |
| Below 30 Mb mapped bp | 80 | 69 | 57 | 209 | Too little sequence to estimate richness. |
| Below 5 SGBs at >=10% breadth | 25 | 22 | 19 | 218 | Clears 30 Mb but resolved no community. |
| **Final analytic cohort· Supplementary Text 1** | | | | | |
| **Final analytic cohort** | 18,046 | 18,060 | 18,075 | 3,887 | One primary run per biosample; secondary runs kept as technical replicates throughout. |
| **Hosts** | 14,613 | 14,613 | 14,624 | 2,942 | One primary run per biosample; secondary runs kept as technical replicates throughout. |
| **Role in this study** | Non-industrial-enriched reference bound | Industrial-enriched reference bound | Primary reference for the reported results | Negative control only – **Supplementary Text 4K** and **Table S5** | The chimpanzee column contributes to no reported human result. It is carried through the same cascade so the control cohort is reproducible on the same terms as the three human references. |

#### Table S3. Figure-specific cohort matrix

Detection spec throughout: ≥10% breadth coverage (10pct). Source files are given relative to the archived analysis outputs (DOI 10.5281/zenodo.23444503, {Citation}); paths of the form results/<reference>/… refer to that archive.

| Figure / panel | Analysis question | Parent cohort | Downstream analytic subset | Exact N / group counts | Detection spec | Primary source file(s) | Notes |
|---|---|---|---|---|---|---|---|
| *Block A — Main-text figure cohorts (frozen from Supplementary Text 1)* | | | | | | | |
| Fig. 1 | Lifestyle diversity and geographic distribution | Final analytic cohort | Lifestyle-classified subset (no further restriction) | 18,075 total<br>Hunter-gatherer: 401<br>Agro-pastoralist: 3,697<br>Industrial: 13,977 | 10% breadth | section1_diversity/main/dataset_overview.csv; section1_diversity/supplement/per_study_metadata.csv | Global overview; all lifestyle strata included |
| Fig. 2A–C | Richness across lifestyle groups (raw counts) | Lifestyle-classified subset | Same as parent; no additional matching applied | 18,075 total<br>Hunter-gatherer: 401<br>Agro-pastoralist: 3,697<br>Industrial: 13,977 | 10% breadth | gmw_richness_summary.csv; threshold-sweep outputs | Richness panels only; no depth matching at this step |
| Fig. 2D–F | Beta-diversity / composition across lifestyle groups | Lifestyle-classified subset | Depth-matched subset (Industrial vs Non-industrial) | Industrial matched: 3,727<br>Non-Industrial matched: 3,727<br>Total matched: 7,454 | 10% breadth | alpha_diversity_matched_Industrial_vs_Non-Industrial_10pct.csv; beta_diversity_pcoa_matched.csv | Matched subset only. Do not carry forward the lifestyle-classified N here. |
| Fig. 3 | Taxon accumulation by lifestyle; | Lifestyle-classified subset | Same as parent; both Carter and | 18,075 total; both references used | 10% breadth | Accumulation outputs (Carter + Leviatan); threshold-sweep outputs | Pooled by lifestyle; reference- |

| Figure / panel | Analysis question | Parent cohort | Downstream analytic subset | Exact N / group counts | Detection spec | Primary source file(s) | Notes |
|---|---|---|---|---|---|---|---|
| | reference sensitivity | | Leviatan references applied | | | | specific detail from accumulation outputs |
| Fig. 4A–D | VANISH / BloSSUM pooling and detection summaries | Lifestyle-classified subset | Industrial 13,977 (VANISH); Non-industrial 4,098 (BloSSUM) | Industrial: 13,977<br>Non-industrial: 4,098 | 10% breadth | vanish_pooling_industrial_10pct.csv; vanish_plus_pooling_industrial_10pct.csv; blossum_pooling_ni_10pct.csv; blossum_plus_pooling_ni_10pct.csv; macro_pooling_detection_10pct.csv | Pooling / detection panels; feeds **Tables S6** and **S7** |
| Fig. 4E–F | List membership and list overlap | Lifestyle-classified subset | Same as parent | 18,075 | 10% breadth | prevalence_gmw_*.csv; table_s4_list_membership.csv; table_s5_overlap_summary.csv | List-definition and overlap panels; distinct from pooling panels above |
| Fig. 5A–C | Sequencing depth distribution (A); raw richness (B); GMW-corrected richness (C) across broad health triplet | Final analytic cohort | Non-Industrial + Industrial healthy + Industrial unhealthy | 17,854 total (excl. 221 Industrial unknown + mixed/unclear lifestyle) | 10% breadth | health_richness_gmw_{ref}_summary_NI_health_10pct_perSample.csv | Per-sample file; 221 Industrial-unknown excluded from this panel only |
| ***Block B — Fig. 5D–H condition-specific cohorts (from health_per_condition_cmh_10pct.csv, Method = Geom-mean-weighted CMH)*** | | | | | | | |
| Fig. 5D Obesity | GMW richness: disease vs control | Industrial case-control eligible | Study-stratified CMH subset | 29 studies<br>Controls: 6,603<br>Cases: 1,219 | 10% breadth | health_richness_gmw_10pct.csv (filter: Method = Geom-mean-weighted CMH) | Plotted in Fig. 5D |
| Fig. 5D Glucose Intolerance | GMW richness: disease vs control | Industrial case-control eligible | Study-stratified CMH subset | 6 studies<br>Controls: 856<br>Cases: 440 | 10% breadth | Same as above | Plotted in Fig. 5D |
| Fig. 5D IBD | GMW richness: disease vs control | Industrial case-control eligible | Study-stratified CMH subset | 6 studies<br>Controls: 726<br>Cases: 253 | 10% breadth | Same as above | Plotted in Fig. 5D |
| Fig. 5D CRC / Adenoma | GMW richness: disease vs control | Industrial case-control eligible | Study-stratified CMH subset | 8 studies<br>Controls: 610<br>Cases: 801 | 10% breadth | Same as above | Plotted in Fig. 5D |
| Fig. 5E | Prevalence-shift distributions: **VANISH$^{H+}$ and BLOSSUM$^{H+}$** vs prevalence-matched background | Industrial case-control eligible | Same per-condition CMH subsets as 5D (see rows above) | Same N as 5D rows above | 10% breadth | health_per_taxon_gmw_10pct.csv; health_per_taxon_prevmatched_10pct.csv | Plotted in Fig. 5E |
| Fig. 5F-G | Comparison of the health-association for the 6 focal lists | Industrial case-control eligible | Same per-condition CMH subsets as 5D (see rows above) | Same N as 5D rows above | 10% breadth | table_s12_method_comparison_combined.csv | Plotted in Fig. 5F-G |
| Fig. 5H | Head-to-head comparison: do the derived lists lie further from the prevalence-matched background than the original lists? | Industrial case-control eligible | Same per-condition CMH subsets as 5D (see rows above) | Same N as 5D rows above; 8 paired comparisons per derived tier (4 conditions × 2 list families) | 10% breadth | table_s12_method_comparison_combined.csv | Plotted in Fig. 5H; Wilcoxon signed-rank over the 8 pairs, \|mean z\| against the original list |
| ***Block C — Supplementary diagnostics*** | | | | | | | |
| Technical replicate screen | Identify and flag within-study replicate pairs | Full run inventory | Secondary runs linked to primary samples | 3,346 runs 1,731 primary samples | — | tech_rep_summary_10pct.csv; tech_rep_pairs_10pct.csv | — |
| Read-length sensitivity | Richness stability across read-length terciles | Lifestyle-classified subset | Read-length tercile subsets (Short / Medium / Long) | **8,751 samples total (10% breadth)**<br>Short tercile: 2,914 (median RL 88.8 bp)<br>Medium tercile: 2,923 (median RL 100.0 bp)<br>Long tercile: 2,914 (median RL 151.0 bp) | 10% breadth | read_length_richness_summary.csv; figure_read_length_diagnostic.png | Predicted richness at 3 Gb: Short 177.1; Medium 212.7; Long 218.0 |
| Carter-only robustness | Richness ratio (Carter vs Other NI) under three correction methods | Lifestyle-classified subset | Carter HG subset vs Other Non-industrial | **Raw N not reported in robustness_summary_carter.csv.**<br>10% breadth ratios:<br>Uncorrected: 4.59 (95% CI 4.23–4.86)<br>Depth-matched: 3.52 (95% CI 3.25–3.86)<br>IPW: 2.69 (95% CI 2.58–2.79) | 10% breadth | robustness_summary_carter.csv | Ratio = Carter richness / Other NI richness. Fill raw N from parent lifestyle-classified subset if needed for table. |

#### Supplementary Text 2. Processing pipeline, reference catalogues, and quality control

This study does not compare author-supplied count tables or previously processed abundance matrices. Instead, all retained metagenomes were reprocessed from raw reads through a common alignment and quality-control workflow and then analyzed against two complementary gut bacterial reference catalogues. This note explains why a reference-based approach is necessary for the present question, what processing steps were applied uniformly to all samples, how the Carter and Leviatan catalogues differ, and why the central results are unlikely to be artifacts of read length, technical replication, or reference choice.

##### 2A. Why a reference-based approach is necessary for this question

Our primary question is whether taxa that appear depleted in industrial populations are truly absent or instead persist at low prevalence and are difficult to detect. That question requires sensitivity to the low-abundance tail of the abundance distribution. A de novo assembly-and-binning approach is invaluable for reference construction, but within any single sample it preferentially recovers the most abundant species and is strongly biased towards detecting more species in deeply sequenced samples. In a dataset where industrial samples are, on average, more shallowly sequenced than non-industrial samples, an assembly-first comparison would therefore exaggerate apparent “missingness” in industrial microbiomes.

We therefore use reference-based alignment as the primary detection framework. Aligning reads to an existing catalogue lowers the effective detection threshold relative to de novo genome recovery and preserves sample-level information needed for per-sample richness, pooled accumulation, and health-association analyses. The trade-off is that the reference itself can be biased by how and from which populations it was constructed. Rather than assuming any single reference is neutral, we treat reference choice as a potential source of bias and address it explicitly by aligning all samples to two complementary catalogues assembled from contrasting source populations.

##### 2B. Uniform preprocessing and alignment

All retained metagenomes were reprocessed from raw reads through the same workflow. Reads were downloaded from the NCBI SRA or ENA, quality- and adapter-trimmed with Trim Galore, filtered against PhiX and the human hg38 reference genome to remove non-microbial host sequence, repaired with bbtools repair.sh, and aligned independently to the Carter and Leviatan reference catalogues (*3*) to generate BAM alignment files (*4*). Coverage was then summarized with samtools coverage and aggregated from the contig level to the SGB level.

The output of this stage is four core sample-by-SGB matrices: breadth and depth for the Carter reference and breadth and depth for the Leviatan reference. The Carter matrices contain 3,971 SGBs and the Leviatan matrices 3,594 SGBs. The full workflow is implemented in Snakemake (*2*) so that all samples are processed identically and failed jobs can be rerun automatically with increased memory or runtime allocations.

A major challenge in large bacterial reference catalogues is multimapping. As the number of related genomes in the reference grows, so does the probability that a 100–150 bp fragment can align to multiple SGBs. We therefore use a deliberately conservative filter and discard all reads with MAPQ < 30. This removes a substantial fraction of mapped reads but greatly reduces incorrect assignment among closely related genomes already present in the reference. A second problem survives this filter. Reference genomes can carry short mobile elements, most often plasmids, that are shared across species, so reads from one taxon may map confidently to a short stretch of another taxon's genome. This is why we gate detection on breadth of coverage rather than on depth: a depth threshold can be met by reads piling up on a single shared region, whereas a breadth threshold requires the reads to be distributed across the genome, which contamination of this kind cannot achieve.

#### 2C. The Carter and Leviatan references capture different discovery biases

Neither reference catalogue is unbiased. Both were built by recovering genomes from metagenomic data, and both therefore inherit the lifestyle composition and sampling design of the datasets from which they were assembled.

The Carter catalogue (*7*) is enriched for genomes recovered from deeply sequenced Hadza and Nepali non-industrial samples and is therefore expected to be more sensitive to taxa common in non-industrial settings or taxa that require very deep per-sample sequencing to be discovered. By contrast, the Leviatan catalogue (*8*) is enriched for genomes recovered from a very large industrial dataset, dominated by Israeli and other industrial cohorts, and is therefore expected to be more sensitive to taxa common in industrial settings or taxa discoverable through broad sampling across many hosts.

These differences matter for any claim about apparent microbial absence. A taxon that was preferentially discovered in a deeply sequenced non-industrial cohort will, by construction, be easier to "find" later when reads are mapped back to that reference. Conversely, a taxon preferentially discovered in a large industrial cohort will be easier to detect in industrial-aligned analyses. Reference choice can therefore bias apparent missingness in opposite directions depending on which catalogue is used.

For this reason, all key persistence analyses are performed against both references. Our inference is not that either catalogue is free of ascertainment bias, but rather that robust conclusions should survive both opposing biases. In that sense, the Carter and Leviatan references act as complementary bounds: the former is enriched for non-industrial discovery, the latter for industrial discovery, and the most credible biological signal is the one that persists across both.

In addition to lifestyle composition, the two catalogues also differ in discovery design. Broadly, Carter reflects a deep dive into relatively few non-industrial individuals, whereas Leviatan reflects a much broader sweep across many industrial individuals. These sampling designs probe different parts of the abundance distribution: deep per-sample sequencing is better at recovering rare members of an individual microbiome, whereas broad population sampling is better at capturing taxa that bloom in only a subset of hosts. The relevant comparison for the present study is therefore not "which catalogue is correct?", but whether the major conclusions hold across two catalogues whose biases point in opposite directions.

The biases inherent in the Carter and Leviatan references are detailed in **Table S4**.

#### 2D. Creating a combined reference from the Carter and Leviatan references

We integrated both human gut microbiome species-level genome bin (SGB) (*5*) catalogues detailed in the previous section and in **Table S4** into a single non-redundant reference: namely, the Leviatan catalogue (3,594 SGBs) (*8*) and the Carter catalogue (3,971 SGBs) (*7*). Each catalogue's source FASTA was split into one file per SGB based on contig-name prefix (Leviatan: `Rep_xxxx`; Carter: `GCA_xxxxxx.V` or `MGYGxxxxxxxxx`). All 7,565 SGBs were sketched with sourmash v4.9.4 at $k = 31$, scaled = 1,000, using the `manysketch` command of the branchwater plugin (v0.9.15) so that each sketch is named by its source genome. The results of this section are summarised in **Data S3**.

**Pairwise containment and symmetric ANI.** All-vs-all cross-catalogue containment was computed with `manysearch` at a minimum containment of 0.2 (*9*) (approximately 95% ANI at $k = 31$ (*10*)), retaining both directional containment-derived ANI estimates per pair. Because containment is asymmetric and can be inflated in one direction by large or chimera-prone MAGs that "swallow" the hashes of smaller genomes, we used the floor of the two directional estimates, min(query_containment_ani, match_containment_ani), as the operational ANI between any two SGBs. Two SGBs from different catalogues were considered same-species candidates when this symmetric min-ANI was ≥ 0.95. The symmetric criterion excluded 1,940 of the 8,413 pairs that a max-ANI rule would have admitted; the excluded pairs were enriched for high containment in one direction only and low containment in the reverse, consistent with hash inflation by large MAGs.

**Combined catalogue by complete-linkage clustering.** We built a 7,565 × 7,565 distance matrix (1 − min-ANI; missing entries set to 1.0) and clustered with complete-linkage hierarchical agglomeration (`scipy.cluster.hierarchy.linkage` (*99*), method = "complete"), cutting the dendrogram at distance 0.05 (min-ANI = 0.95). Complete-linkage was chosen over single-linkage (the implicit basis for "connected components") because the latter chains transitively-related but mutually distant SGBs into spuriously large clusters. For example, a single MAG with ≥ 95% min-ANI to 80 distinct partners in the opposing catalogue would, under single-linkage, fuse all 80 + 1 into one cluster even though the partners themselves were defined as separate SGBs at ≥ 95% ANI by their source catalog. Complete-linkage guarantees, by contrast, that every pair of SGBs *within* the same cluster shares ≥ 95% min-ANI, and never merges sets whose members are not all mutually within the threshold.

Because each input catalogue already enforces ≥ 95% ANI between its own SGBs, every resulting combined cluster contains at most one SGB from each catalog. This produced 5,081 combined SGBs**:** 2,484 "shared" 1:1 Leviatan ↔ Carter pairs, 1,110 SGBs unique to Leviatan, and 1,487 SGBs unique to Carter.

**Representative selection for shared clusters.** For each of the 2,484 shared clusters, the representative used downstream was selected empirically based on mapping performance rather than catalogue of origin. We computed, for each candidate representative, the sum of per-sample mean-coverage values across the full cohort using each catalogue's pre-existing read mapping, and retained the higher-scoring representative. Carter genomes were selected in 1,396 / 2,484 (56.2 %) of shared clusters and Leviatan genomes in 1,088 (43.8 %); no exact ties occurred. Leviatan-only and Carter-only clusters had no choice and used their single available representative. The combined reference FASTA was then assembled by streaming the chosen representative's contigs from the appropriate source file (`leviatan_rep_all_renamed.fa` or `Hadza_UHGG.fna`).

**GTDB taxonomy.** Each original SGB was classified against the GTDB-RS226 representative sketch collection (143,384 genomes) (*100*) with branchwater's `fastmultigather`, and lineages were assigned per genome using `sourmash tax genome` at a permissive ANI threshold of 0.80 so that divergent SGBs received a lineage at the deepest confident rank rather than being reported as unclassified. 3,506 / 3,594 Leviatan SGBs and 3,244 / 3,971 Carter SGBs received GTDB lineages.

**Coverage matrices.** Combined mean-coverage and mean-depth matrices restricted to the 5,081 representatives were constructed by selecting the `<rep>_mean` and `<rep>_std` columns corresponding to each combined SGB's chosen representative from the source coverage matrices and outer-joining on sample identifier, so that samples mapped against only one catalogue retained their available values.

**Within-cluster redundancy and a boundary-case caveat.** By construction, the combined catalogue contains no two SGBs in the same cluster that fail to share ≥ 95% min-ANI: every cluster is a clique at the species threshold. Across clusters, however, the complete-linkage criterion necessarily admits a small number of boundary pairs at ≥ 95% min-ANI — these are pairs whose joint co-clustering would have required including a third member at < 95% ANI to one of them. We identified 3,989 such cross-cluster pairs (compared with 2,484 within-cluster pairs). 3,359 of these 3,989 pairs (84 %) involve SGBs in the family *Coriobacteriaceae*, a clade with well-documented continuous strain variation around the 95% ANI species threshold, where roughly 80 SGBs in each catalogue hover within 95% min-ANI of one another. The remaining 16 % (630 pairs involving only "clean" SGBs with fewer than ten cross-catalogue partners) are scattered across diverse families and represent the natural boundary noise of any operational species threshold.

We chose to retain these boundary pairs as separate clusters rather than collapse them, because merging them would either reintroduce single-linkage chaining (forming clusters whose members are not all mutually ≥ 95% ANI) or require non-uniform clustering thresholds across the tree. Affected SGBs (especially those in *Coriobacteriaceae* and *Eggerthellaceae*) are tagged in the combined metadata table (`fuzzy_boundary_member = True`) so that downstream analyses sensitive to species-level boundary placement – for example, within-species strain abundance estimation – can be interpreted with appropriate caution. Family- and higher-rank analyses are unaffected.

#### 2E. Why breadth and depth are carried forward jointly

A fixed breadth threshold is a useful first-pass filter, but breadth alone discards information that is especially important for low-abundance detections. Host and PhiX reads can be removed directly, yet short contaminant or plasmid-like regions that are embedded within a microbial reference may still generate a misleading signal. Such regions tend to produce relatively high depth at very low breadth, because reads pile up on a short fragment rather than accumulating across a large portion of the genome.

Genuine representation of an SGB produces a different signature. As sequencing depth increases, coverage spreads across a longer region of the genome, and the joint behavior of breadth and depth becomes inconsistent with a short fragment alone. We therefore retain both breadth and depth for every sample-SGB pair and use their joint relationship in the downstream probabilistic detection framework. The full Bayesian detection and accumulation model is described in **Supplementary Text 4**. The important QC point here is that detection is not inferred from breadth in isolation: the breadth signal must also be plausible given the observed depth.

#### 2F. Read-length and technical-replicate quality control

Read length varies substantially across studies and is partially confounded with lifestyle, because many older industrial datasets were sequenced with shorter reads whereas several non-industrial datasets used longer reads. In the present dataset, longer reads yield modestly higher observed richness at a fixed number of mapped bases. However, this effect is small relative to the between-lifestyle richness difference and remains secondary after conditioning on sequencing depth (see **Fig. S4**). We therefore treat read length as a minor technical covariate rather than a plausible primary explanation for the major lifestyle-associated differences reported in the main text.

Technical replicate analyses further support the stability of the pipeline. Richness estimates are highly reproducible across primary and secondary runs derived from the same biosample, and within-sample Bray-Curtis dissimilarity is usually close to zero. Most secondary runs were therefore collapsed with the corresponding primary run as technical replicates. A minority of replicate pairs showed elevated Bray-Curtis dissimilarity. These were nonetheless retained as technical replicates rather than promoted to independent samples, because the dissimilarity distribution decays smoothly with no natural threshold, and the pairs that would have been promoted are concentrated in repeat sampling of the same participant and in one study's duplicate DNA-extraction protocol. This choice is conservative with respect to richness and keeps replicate handling symmetric between the compared lifestyle groups.

Together, these processing and QC steps address the main technical concerns relevant to the downstream analyses. All retained metagenomes are processed identically, ambiguous mappings are filtered conservatively, and reference bias is bounded by dual alignment to catalogues enriched for opposite lifestyle contexts. The technical covariates that we could evaluate directly do not explain the core patterns identified in the study, namely reduced individual-level richness but broad population-level persistence.

Read-length effects are summarized in **Fig. S14**, and technical-replicate reproducibility diagnostics are shown in **Fig. S2–S3**.

**Table S4. Comparison of the Carter and Leviatan reference catalogues.** The Carter catalogue is enriched for genomes discovered in deeply sequenced non-industrial cohorts, whereas the Leviatan catalogue is enriched for genomes discovered in a very large industrial cohort with shallower sequencing. Because these ascertainment biases point in opposite directions, reference choice can influence apparent taxon missingness, prevalence, and pooled richness. The primary persistence analyses are therefore evaluated against both catalogues.

| Feature | Carter reference | Leviatan reference | Why it matters here |
|---|---|---|---|
| **Source paper** | Carter et al. 2023 (*7*) | Leviatan et al. 2022 (*8*) | Identifies the two complementary catalogues used throughout the study. |
| **SGBs in current analysis freeze** | 3,971 SGBs | 3,594 SGBs | Defines the size of the sample × SGB breadth/depth matrices produced by the preprocessing pipeline. |
| **Dominant discovery cohorts** | Deeply sequenced Hadza and Nepali gut metagenomes | Very large Israeli-centered industrial gut metagenome collection | The catalogues were not assembled from the same ecological or sampling background. |
| **Dominant lifestyle enrichment** | Non-industrial-enriched | Industrial-enriched | Each reference is expected to be more sensitive to taxa preferentially discovered in its source lifestyle context. |
| **Discovery design** | Deep sequencing of comparatively fewer individuals | Broader population sampling across many individuals | The two catalogues sample different parts of the abundance/prevalence landscape. |
| **Approximate discovery sequencing regime** | Very deep per-sample sequencing | Much shallower per-sample sequencing, but much larger cohort breadth | Deep sequencing favors recovery of low-abundance taxa within individuals; broad sampling favors taxa prevalent across many hosts. |
| **Taxa the catalogue is best positioned to recover** | Taxa detectable only with very deep sequencing, especially in non-industrial hosts | Taxa that recur across many industrial hosts and can be discovered through broad cohort coverage | Explains why apparent missingness can depend on reference choice. |
| **Expected blind spot if used alone** | Taxa mainly recoverable through broad industrial sampling or taxa poorly represented in non-industrial discovery cohorts | Taxa requiring very deep per-sample sequencing or taxa strongly associated with non-industrial contexts | One reference alone could distort conclusions about absence, persistence, or relative richness. |
| **Likely direction of ascertainment bias in this manuscript** | May overstate apparent non-industrial richness and industrial 'missingness' | May overstate apparent industrial richness and understate non-industrial enrichment | The two biases point in opposite directions, which is why results are evaluated against both references. |
| **Role in this study** | Non-industrial-enriched reference bound | Industrial-enriched reference bound | Concordant signals across both references are less likely to be artifacts of catalogue construction. |
| **Shared SGBs with the other reference** | 2,484 | 2,484 | Quantifies the common reference core used by both catalogues. |
| **SGBs unique to this reference** | 1,487 | 1,110 | Quantifies non-overlap and helps interpret reference-specific detections. |

#### Supplementary Text 3. Correcting sequencing-depth bias in per-sample richness and composition

##### 3A. What Fig. 2 measures

We first implemented a geometric-mean-target weighting (GMW) method to re-weight microbial samples (*11*) and directly compare groups with systematically different sequencing depths (**Figs. S5–S13**). In the context of this study, diversity refers to richness (*101*), namely the total number of microbial species detected in an individual sample. Composition refers to the identities and abundances of the detected microbial species in that sample. **Fig. 2** therefore combines a depth-corrected richness analysis (panels **A–C**) with complementary composition analyses based on between-sample dissimilarity (panels **D–F**). The primary supporting diagnostics for this note are **Figs. S9–S13**, with sensitivity analyses using the samples analyzed by Carter et al. 2023 (*7*) shown in **Figs. S5–S8**.

##### 3B. Why raw richness comparisons are biased by sequencing depth

Industrial and non-industrial samples differ systematically in sequencing depth. A sample with 10× more reads will detect more taxa simply due to sampling depth, not biology. Any raw richness comparison between groups therefore conflates a real biological signal with a depth artifact. 1:1 nearest-neighbor matching addresses this by discarding unmatched samples, but this loses statistical power. The GMW (geometric-mean-target weighting) approach instead retains all samples and corrects by reweighting (*11*). The systematic lifestyle-level depth imbalance motivating this correction is shown in **Fig. S4**.

##### 3C. The geometric-mean-target weighting (GMW) method

*The target distribution*

The key design choice is what reference distribution to reweight towards. We use the geometric mean of the two groups' depth densities (*12*); if we wish to compare industrial (I) and non-industrial (NI) samples this is:

$$f_{target}(x) = \sqrt{\left(f_I(x) \cdot f_{NI}(x)\right)}$$

where $f_I$ and $f_{NI}$ are their respective sample density distributions as a function of sequencing depth mapped to a metagenome reference (smoothed using kernel density estimation (*102*), see below). This is not arbitrary. We can quantify how different two distributions are using Kullback-Leibler-divergence; choosing the geometric mean as the target distribution ensures that we are equally different from both distributions, so it favors neither group. Reweighting industrial (I) towards this target and reweighting non-industrial (NI) towards the same target produces two populations that are both compatible with the same depth distribution, so any remaining richness difference is not a depth artifact.

We can extend this definition to three groups (e.g. hunter-gatherer (HG), agro-pastoralist (T), industrial (I)), like so:

$$f_{target}(x) = \sqrt[3]{\left(f_{HG}(x) \cdot f_T(x) \cdot f_I(x)\right)}$$

where $f_{HG}(x)$, $f_T(x)$, and $f_I(x)$ are the sample density distributions of HG, T, and I samples mapped to the reference.

*Computing the weights*

For each sample $i$ in group A (say industrial), its weight is:

$$w_i = \sqrt{\left(f_B(\log_{10} mdepth_i)/f_A(\log_{10} mdepth_i)\right)}$$

where $f_A$ and $f_B$ are the density distributions for groups A and B respectively (smoothed using kernel density estimation (*102*), see below). These are expressed in terms of the $\log_{10}$ of the total number of bp mapped to the reference $mdepth_i$ for each group.

The weights are then mean-normalized (divided by their mean so $\text{mean}(w) = 1$) and hard-clipped at 10. This prevents any single sample from dominating the weighted estimate and overly biasing group statistics (as individual microbiome samples still vary significantly in diversity and composition).

*Kernel Density Estimation (KDE) implementation*

To compute the weights, we need to know, for any given sequencing depth, how common that depth is within each group. We estimate this by fitting a smooth continuous curve to the histogram of $\log_{10}$(mapped bases) values within each group. Smoothing is necessary because a raw histogram is coarse and irregular: bins that happen to contain few or no samples produce density estimates of zero, which would cause the weight formula (which involves dividing by the group's own density) to become undefined or arbitrarily large for samples that fall in those gaps. Even where bins are populated, adjacent bins of very different heights create abrupt jumps that make weights unstable: a sample at depth 1.2 Gb and a sample at 1.3 Gb might receive wildly different weights simply due to bin-boundary artifacts rather than any real difference in how common those depths are. The smooth KDE curve eliminates both problems by interpolating continuously across the full range of observed depths.

This curve describes the shape of the depth distribution: for most groups it is roughly bell-shaped on a log scale, but the center and spread differ between industrial and non-industrial samples. The weight assigned to a sample depends on the height of its group's KDE curve at that sample's depth, relative to the other group's curve at the same depth. A sample whose depth is common in its own group but rare in the comparison group receives a low weight, and vice versa.

We use a fast binned implementation (_BinnedKDE) rather than a standard KDE, because evaluating a naive KDE for groups of 3,000–14,000 samples is computationally prohibitive. The implementation works by: (1) binning depth values into 512 histogram bins; (2) choosing a smoothing bandwidth using Silverman's rule, $h = 1.06 \cdot \sigma \cdot n^{-0.2}$, where $\sigma$ is the standard deviation of log-depths and $n$ is the group size (*102*); and (3) convolving the histogram with a Gaussian kernel via Fast Fourier Transform.

*Weighted richness estimate*

For each group, the corrected richness estimate is the weighted mean:

$$\mu_{Industrial} = \Sigma w_i \cdot richness_i / \Sigma w_i$$

$$\mu_{NI} = \Sigma w_j \cdot richness_j / \Sigma w_j$$

Here, we are interested in the difference $\mu_{Ind} - \mu_{NI}$, that is, the estimated richness difference after removing the depth confound.

*Bootstrap inference*

The KDEs are held fixed because they define the estimand rather than estimate a nuisance parameter: the weights $\sqrt{f_{\text{other}}/f_{\text{own}}}$ standardise both groups to the same reference depth distribution, the geometric mean $\sqrt{f_I/f_{NI}}$. Re-fitting them within each replicate would move that reference, so each replicate would estimate the richness contrast standardised to a slightly different depth distribution, and the resulting spread would conflate sampling variation in richness with drift in the definition of the contrast. Samples, not hosts, are resampled; a host-level bootstrap and a one-sample-per-host analysis are reported in **Supplementary Text 7C**. This is faster and avoids the instability of re-fitting KDEs on small bootstrap samples. Confidence intervals are percentile-based ($2.5^{th}$–$97.5^{th}$). The p-value is computed as a two-sided tail area from the bootstrap distribution of $\mu_{Ind} - \mu_{NI}$:

$$p = 2 \cdot min(\text{fraction of bootstraps with } \mu_{Ind} - \mu_{NI} > 0, \text{ fraction with } \mu_{Ind} - \mu_{NI} < 0)$$

If the true richness difference were zero, roughly half the bootstraps would give a positive difference and half negative; a p-value near zero means nearly all bootstraps agree on the direction, which is strong evidence of a real difference.

The number of bootstrap replicates scales adaptively with the smaller group size: $n_{boot} = clip\left(1000 \cdot \sqrt{(n_min/500)}, 1000{,}5000\right)$. This ensures enough replicates to characterise the bootstrap distribution reliably when samples are few, while avoiding unnecessary computation when groups are large and the distribution is already well-defined by fewer replicates.

*Effective sample size*

The ESS (effective sample size) after weighting is reported as:

$$ESS = (\Sigma w_i)^2 / \Sigma w_i^2$$

This is the standard Kish ESS formula, which quantifies how many equally-weighted samples would give the same precision as the weighted sample. High weight dispersion (a few samples with $w$ near 10 and many near 0) produces a low ESS fraction, indicating the correction is expensive and precision is reduced. For example, if all 1,000 samples in a group receive weight ≈1.0 (the depth distributions largely overlap), ESS ≈ 1,000. If a few hundred samples dominate with high weights while the rest contribute almost nothing, ESS might be only 300, meaning that we are effectively working with a much smaller dataset despite having 1,000 samples available. We typically achieve an ESS fraction (normalized to 1) of >60%. This is good relative to a naïve rarefaction or depth-matching method (where we might lose an order of magnitude).

*Did the depth correction actually matter?*

We also test, separately for each group, whether the depth correction meaningfully changed that group's own richness estimate. This is important because the correction is only doing real work if the depth distributions were genuinely imbalanced to begin with.

For each group, we compare the raw (unweighted) mean richness to the distribution of corrected means produced by the bootstrap. If industrial samples happened to be evenly spread across sequencing depths, the weights would all be close to 1.0, the correction would change almost nothing, and the raw and corrected means would be statistically indistinguishable. If, on the other hand, industrial samples were systematically more deeply sequenced than non-industrial samples, the correction would downweight the deepest industrial samples and upweight the shallowest ones, shifting the corrected industrial mean noticeably below the raw mean, and this test would confirm that shift was larger than expected by chance.

Concretely, the test computes a z-score by dividing the difference between the raw mean and the bootstrap mean of the corrected estimate by the bootstrap standard deviation of the corrected estimate. A significant result (shown as a bracket on each group's violin in **Figs. S6, S11–S12, S24**) means the correction moved that group's estimate by more than sampling noise alone can explain. Seeing this bracket significant for both groups in opposite directions (Industrial corrected downward, non-industrial corrected upward) is positive confirmation that depth confounding was present and has been successfully removed.

**3D. Richness results and "did correction matter?" diagnostics**

After correcting for sequencing depth, industrial populations still harbored lower per-sample richness than non-industrial populations. The correction substantially reduced the apparent gap relative to the raw comparison, confirming that lower sequencing depth in industrial cohorts had inflated the magnitude of the richness deficit, but it did not eliminate the deficit itself (**Fig. 2C; Figs. S5–S13**). Across detection thresholds, the corrected estimate remained consistently below the non-industrial value, indicating that the correction is doing real work but is not driving the biological result.

**3E. Composition analyses for Fig. 2D–F**

To quantify between-sample composition in **Fig. 2D-F**, we used two complementary dissimilarity measures. For composition analyses, we used exact depth-matched subsets rather than GMW weighting, because distance-based ordination and pairwise dissimilarity summaries are not naturally handled by the weighting framework used for richness analyses.

Panels **D** and **E** are computed on a 1:1 industrial-to-non-industrial depth-matched set. Samples were paired by greedy nearest-neighbor matching on $\log_{10}$ total mapped bacterial bases, without replacement, accepting a pair only where the two depths differed by at most 3%. The tolerance is pragmatic rather than principled: a tighter window leaves too few matched pairs to estimate the within- and between-group distance distributions reliably, and a wider one reintroduces the depth difference the matching is there to remove. This yielded 3,727 matched pairs, the cohort reported for these panels in **Table S3**; the non-industrial side of that set comprises 277 hunter-gatherer and 3,450 agro-pastoralist samples.

Because the number of pairwise distances grows with the square of group size, each distribution was subsampled before its distance matrix was computed. Within-group distributions were capped at 1,500 samples per group, so all 277 hunter-gatherer samples were retained while the agro-pastoralist and industrial groups were reduced to 1,500 each, giving 1,124,250 pairs per group. The between-group reference distribution was computed from a further random draw of 1,000 industrial and 1,000 non-industrial samples, giving 1,000,000 cross-group distances. These caps are applied after matching and are purely computational; they do not change which samples are eligible for the comparison.

Panel **F** is not depth-matched. Samples are instead assigned to equal-count sequencing-depth bins and compared within bins, as described below.

Jaccard distance is computed on the presence/absence matrix, while Bray-Curtis dissimilarity is computed on a relative abundance matrix, where SGB abundance is estimated as mean depth × genome length and then normalized by each sample's total mapped bacterial bases.

**3F. Coordinate-invariant axis projection for Fig. 2F**

To visualize how hunter-gatherer, agro-pastoralist, and industrial microbiomes differ in composition without relying on arbitrary axis orientation from a single ordination, we projected samples into a coordinate-invariant two-dimensional space defined directly from the lifestyle centroids within each depth bin. The input remained the same depth-matched composition data described above: Bray-Curtis dissimilarities were computed on the relative-abundance matrix, and the matched subset was used to reduce confounding by sequencing depth. Rather than interpreting raw PCoA axes directly, however, we used the ordination only as an intermediate embedding from which a more interpretable geometric summary could be derived.

For each depth bin (**Fig. S18**), let $H$, $T$, and $I$ denote the hunter-gatherer, agro-pastoralist, and industrial centroids in the ordination space. We first defined the primary lifestyle-contrast vector as the directed line from $H$ to $I$. This vector captures the main compositional contrast between the two endpoint lifestyles. We then expressed every sample position relative to this line using two quantities.

First, we computed the sample's fractional position along the hunter-gatherer→industrial axis, denoted $p$. This is the normalized projection of the sample onto the line joining the hunter-gatherer and industrial centroids, with $p = 0$ at the hunter-gatherer centroid and $p = 1$ at the industrial centroid. Values between 0 and 1 therefore indicate positions intermediate between the two endpoint lifestyles, while values below 0 or above 1 indicate extension beyond either endpoint.

Second, we computed the sample's signed off-axis displacement, denoted $q$. This is the perpendicular distance from the sample to the hunter-gatherer→industrial axis, with the sign chosen so that positive values point toward the side of the space occupied by agro-pastoralist centroids. Thus, $q$ measures how far a sample departs from the direct hunter-gatherer→industrial trajectory, and in which direction.

Formally, if $x$ is a sample coordinate, $u = (I - H)/||I - H||$ is the unit vector from hunter-gatherer to industrial, and $m = ||I - H||$ is the hunter-gatherer–industrial separation, then

$$p = \big((x - H) \cdot u\big)/m$$

and

$$q = (x - H) \cdot u_{\perp}$$

where $u_{\perp}$ is the unit vector perpendicular to $u$, oriented so that positive values point toward the agro-pastoralist side of the space.

This transformation is useful because it separates two biologically different questions. The coordinate $p$ asks how far a sample has moved along the main hunter-gatherer→industrial contrast. The coordinate $q$ asks whether that movement follows the direct hunter-gatherer→industrial path or instead deviates into a distinct compositional direction. If agro-pastoralist microbiomes were simply mixtures of the two endpoint states, they would be expected to lie near intermediate $p$ values with $q \approx 0$. Instead, we observed that agro-pastoralist samples were intermediate in $p$ but consistently positive in $q$, indicating that agro-pastoralist microbiomes occupy a distinct side-branch of the compositional landscape rather than a simple linear midpoint.

This distinction is biologically important. The direct hunter-gatherer→industrial contrast approximates the signal emphasized by the original VANISH/BloSSUM framing, whereas the off-axis agro-pastoralist displacement suggests an additional component of change associated with agriculturalization. The later shift from agro-pastoralist toward industrial composition then occurs along a partially orthogonal direction. In other words, the broader non-industrial→industrial contrast is not fully collinear with the original hunter-gatherer→industrial contrast.

To assess whether this pattern was driven by a few large cohorts, we also summarized each study by its centroid in the same coordinate system (**Fig. S19**). The per-study centroids reproduced the same broad geometry: hunter-gatherer studies clustered near $p \approx 0$, industrial studies near $p \approx 1$, and agro-pastoralist studies occupied intermediate $p$ values but remained displaced toward positive $q$. This confirms that the off-axis position of agro-pastoralist microbiomes is not just a property of pooled sample density, but is also visible at the study level.

#### Supplementary Text 4. Finite-catalogue accumulation modelling of SGB discovery

The analyses in **Figs. 3** and **4** and **Figs. S15–S17** address a finite-catalogue discovery problem. We have a fixed reference catalogue of species-level genome bins (SGBs), and we ask how many of these reference SGBs would be detected as sampling effort increases. The central inferential target is the asymptote, i.e., the number of reference SGBs that would eventually be detected under a specified sampling protocol.

The mathematical issues differ depending on the response being plotted.

First, for the main results, an SGB is counted as detected in a group once it has been detected at least once. This is the case $k = 1$. For $k = 1$, the accumulation curve is a first-detection curve. This is why simple three- and four-parameter empirical curves, such as Hill-type Michaelis-Menten and Richards curves, can be appropriate for the main figure.

Second, the supplementary analyses in **Fig. S16** count an SGB as detected only once it appears in at least $k$ samples or studies, for example $k = 2,3,5$. These repeated-detection curves are not slower versions of the first-detection curve but a different object altogether (specifically, binomial-tail or Poisson-binomial-tail functions) which is why they call for the beta-binomial and beta-mixture models developed below rather than the empirical curves used for the main figure.

Third, samples and studies differ in sequencing depth. Equal-depth beta-binomial models are therefore approximations. This issue can be partially resolved by fitting the median accumulation bootstrap (see next paragraph). We suggest a sequencing depth-aware model that could account for differences in sequencing depth but find it to be slow and not necessary given the very accurate fits obtained by an equal-depth model.

Fourth, asymptote uncertainty is estimated by likelihood profiling. For the main $k = 1$ figure, this can be done with a curve-level likelihood for the bootstrap median accumulation curve. For the beta-binomial and beta-mixture supplementary analyses, this can also be done with a catalogue-level likelihood based on the incidence counts of all reference SGBs.

##### 4A. Notation and fixed-catalogue asymptote

Let $S$ be the number of SGBs in the reference catalogue, where we index reference SGBs by:

$$i = 1, \dots, S.$$

Let $n$ denote the number of sampled units. A sampled unit may be an individual metagenomic sample or a study, depending on the panel. Repeat samples from one host are separate units. Because they share more taxa than samples from different hosts, they slow accumulation, so curves in samples are conservative relative to curves in hosts. Let $k$ denote the minimum number of units in which an SGB must be detected in order to be counted. The ordinary accumulation curve has $k = 1$. Repeated-detection curves use $k \geq 2$.

Let $A_k(n)$ be the expected number of reference SGBs detected in at least $k$ units after sampling effort $n$:

$$A_k(n) = \mathbb{E}\{\text{number of reference SGBs detected in at least } k \text{ units after } n \text{ units}\}.$$

Each reference SGB has an underlying detectability under the sampling protocol. We denote this detectability by $\lambda_i$ when using a continuous sequencing-effort model and by $p_i$ when using an equal-depth Bernoulli model. The asymptote is:

$$S_\infty = \sum_{i=1}^{M} \mathbf{1}\,(\lambda_i > 0).$$

Equivalently, in an equal-depth Bernoulli model,

$$S_\infty = \sum_{i=1}^{M} \mathbf{1}\,(p_i > 0).$$

Since the catalogue is finite,

$$0 \le S_\infty \le M.$$

For any fixed $k$, the infinite-sampling limit is

$$\lim_{n\to\infty} A_k(n) = S_\infty.$$

The reason is that if an SGB has positive detectability, then with infinitely many independent sampling opportunities it will eventually be detected at least $k$ times with probability one. If it has zero detectability under the sampling protocol, it will never be detected.

This means that $S_\infty$ is not merely a curve-fitting constant, but rather the number of reference SGBs that are reachable under the sampling protocol.

Two consequences of estimating $S_\infty$ by curve fitting should be stated explicitly. First, the fitted asymptote is constrained to the interval [max observed richness, $M$], so it can never exceed the catalogue size by construction. Upper confidence limits are therefore censored at $M$ rather than free, and across the accumulation fits reported here roughly three quarters of bootstrap upper quantiles lie within 0.5% of $M$. Second, because of that censoring, proximity of an upper confidence limit to $M$ is not evidence that $S_\infty$ equals $M$. We therefore also report, for every fit, the bootstrap probability that $S_\infty$ lies strictly below $M$ less a tolerance; its median across fits is 0.72 to 0.75 depending on the reference catalogue, meaning that about one quarter of the bootstrap mass cannot distinguish the asymptote from the catalogue ceiling. Reported asymptotes should be read with that limit in mind.

#### 4B. Accumulation of number of SGBs found in at least one sample or study ($k = 1$)

##### 4B.1. $k = 1$ accumulation is a first-detection curve

For $k = 1$, an SGB contributes to the accumulation curve once it has been detected at least once. Let $T_i$ be the sampling effort at which SGB $i$ is first detected. Then:

$$\Pr\{\text{SGB } i \text{ has been detected by effort } n\} = \Pr(T_i \le n).$$

Therefore,

$$A_1(n) = \sum_{i=1}^{M} \Pr(T_i \le n).$$

If we restrict attention to the $S_\infty$ detectable SGBs and model their first-detection efforts as draws from a distribution $F_T$, then:

$$A_1(n) = S_\infty F_T(n).$$

This identity is the key mathematical reason that simple sigmoidal or saturating curves can fit ordinary SGB accumulation. The $k = 1$ accumulation curve is the cumulative distribution function of first-detection efforts, multiplied by the finite-catalogue asymptote.

##### 4B.2. A depth-aware detection model for $k = 1$

Let $e_j$ be the normalized effective sequencing depth of sample $j$. The normalization can be chosen so that $e_j = 1$ corresponds to a standard-depth sample. Let $\lambda_i$ be the intrinsic detectability of SGB $i$ per unit sequencing effort. A standard detection model is:

$$\Pr\{\text{detect SGB } i \text{ in sample } j \mid \lambda_i, e_j\} = 1 - \exp(-\lambda_i e_j).$$

The probability of not detecting SGB $i$ in sample $j$ is therefore

$$\exp(-\lambda_i e_j).$$

For a subset $B$ of samples, we can define the total effective sequencing effort as

$$E_B = \sum_{j \in B} e_j.$$

The probability of not detecting SGB $i$ in any sample in $B$ is

$$\prod_{j \in B} \exp(-\lambda_i e_j) = \exp\left(-\lambda_i \sum_{j \in B} e_j\right) = \exp(-\lambda_i E_B).$$

Thus, the probability of detecting SGB $i$ at least once in the subset is

$$1 - \exp(-\lambda_i E_B).$$

Summing over detectable SGBs gives

$$A_1(E_B) = \sum_{i:\lambda_i > 0} \{1 - \exp(-\lambda_i E_B)\}.$$

If $\lambda$ denotes the detectability of a randomly chosen detectable SGB, then

$$A_1(E) = S_\infty[1 - \mathbb{E}\{\exp(-\lambda E)\}].$$

where the term

$$\mathbb{E}\{\exp(-\lambda E)\}$$

is the Laplace transform of the SGB detectability distribution. Thus, different assumptions about the distribution of SGB detectabilities generate different closed-form accumulation curves.

**4B.3. Derivation of Michaelis-Menten from exponential detectabilities**

Assume that positive SGB detectabilities follow an exponential distribution with rate $K$:

$$\lambda \sim \text{Exponential}(K).$$

The density is

$$f(\lambda) = K\exp(-K\lambda), \qquad \lambda > 0.$$

The probability that a randomly chosen detectable SGB remains undetected after effort $E$ is

$$\mathbb{E}\{\exp(-\lambda E)\} = \int_0^\infty \exp(-\lambda E) K\exp(-K\lambda)\, d\lambda.$$

Combining exponentials,

$$\mathbb{E}\{\exp(-\lambda E)\} = K \int_0^\infty \exp\{-(K+E)\lambda\}\, d\lambda.$$

Because

$$\int_0^\infty \exp\{-(K+E)\lambda\}\, d\lambda = \frac{1}{K+E},$$

we obtain

$$\mathbb{E}\{\exp(-\lambda E)\} = \frac{K}{K+E}.$$

Therefore,

$$A_1(E) = S_\infty\left(1 - \frac{K}{K+E}\right) = S_\infty \frac{E}{K+E}.$$

Thus, Michaelis-Menten accumulation is obtained exactly when SGB detectabilities are exponentially distributed under the Poisson detection model:

$$A_1(E) = S_\infty \frac{E}{K+E}.$$

If the x-axis is the number of samples $n$, and the total effective effort is approximately

$$E \approx \bar{e}n,$$

then

$$A_1(n) = S_\infty \frac{\bar{e}n}{K + \bar{e}n} = S_\infty \frac{n}{K^* + n},$$

where

$$K^* = \frac{K}{\bar{e}}.$$

Thus, a Michaelis-Menten curve in sample number is an equal-average-depth version of the same detection model.

##### 4B.4. Derivation of Hill-type Michaelis-Menten from effort to first-detection

A Hill-type Michaelis-Menten curve has the form

$$A_1(n) = S_\infty \frac{n^h}{K^h + n^h}.$$

This curve is obtained exactly if first-detection effort $T$ follows a log-logistic distribution. Specifically, suppose

$$\Pr(T \leq n) = \frac{n^h}{K^h + n^h}.$$

Since

$$A_1(n) = S_\infty \Pr(T \leq n),$$

we get

$$A_1(n) = S_\infty \frac{n^h}{K^h + n^h}.$$

Equivalently, let

$$Z = \log(T).$$

The Hill curve corresponds to a logistic distribution for $Z$:

$$\Pr(Z \leq x) = \frac{1}{1 + \exp\{-h(x - \log(K))\}}.$$

Setting $x = \log(n)$,

$$\Pr(T \leq n) = \Pr(\log(T) \leq \log(n)) = \frac{1}{1 + \exp\{-h(\log(n) - \log(K))\}} = \frac{n^h}{K^h + n^h}.$$

Thus, the Hill curve is a logistic cumulative distribution function on the log-effort scale. This is biologically plausible as an empirical model when SGB detectabilities span orders of magnitude, because first-detection effort is approximately inversely related to detectability:

$$T_i \approx \frac{1}{\lambda_i}.$$

If $\log(\lambda_i)$ is roughly unimodal, then $\log(T_i)$ is also roughly unimodal, and a logistic distribution can be a useful approximation.

The Hill curve is not particularly accurate at small $n$ and should be interpreted as an empirical first-detection cumulative distribution function over the observed sampling range.

##### 4B.5. Richards curves are empirically useful to model logistic effort for first-detection

The Richards curve is a generalized logistic curve. Let

$$x = \log(n).$$

Define

$$F_R(x) = [1 + \nu \exp\{-b(x - c)\}]^{-1/\nu},$$

where

$$b > 0, \qquad \nu > 0.$$

If the log first-detection effort

$$Z = \log(T)$$

has cumulative distribution function $F_R$, then

$$\Pr(T \leq n) = \Pr(\log(T) \leq \log(n)) = F_R(\log(n)).$$

Therefore,

$$A_1(n) = S_\infty[1 + \nu \exp\{-b(\log(n) - c)\}]^{-1/\nu}.$$

The Hill curve is the special case $\nu = 1$, with a reparameterization of $K$ and $h$. Thus, Richards curves are not introduced because they are assumed to be the unique biological model. Rather, they are flexible four-parameter cumulative distribution functions for first-detection effort on the log-sampling scale.

This justifies using Hill-type Michaelis-Menten and Richards curves in the main $k = 1$ figure: for $k = 1$, accumulation is a first-detection distribution, and these curves are explicit parametric distributions for first-detection effort.

##### 4C. Sequencing-depth variation and the bootstrap median for $k = 1$

The preceding derivations used effective effort $E$, not raw sample count. We therefore need to justify why fitting a curve against sample number $n$ can still be reasonable for the median bootstrap accumulation curve.

Let the available dataset contain $N$ samples with normalized depths

$$e_1, \ldots, e_N.$$

For a random bootstrap subset or random-ordering prefix $B_n$ of size $n$, define

$$E_{B_n} = \sum_{j \in B_n} e_j.$$

Under the $k = 1$ depth-aware detection model,

$$A_1(B_n) = g(E_{B_n}),$$

where

$$g(E) = \sum_{i:\lambda_i > 0} \{1 - \exp(-\lambda_i E)\}.$$

Thus, for $k = 1$, the depth composition of the subset affects the curve only through total effective effort $E_{B_n}$.

Let

$$\bar{e} = \frac{1}{N} \sum_{j=1}^{N} e_j$$

and

$$s_e^2 = \frac{1}{N-1} \sum_{j=1}^{N} (e_j - \bar{e})^2.$$

For a random subset of size $n$ sampled without replacement,

$$\mathbb{E}(E_{B_n}) = n\bar{e}.$$

The finite-population variance is

$$\mathrm{Var}(E_{B_n}) = n\left(1 - \frac{n}{N}\right) s_e^2.$$

Therefore, the relative standard deviation of total effective effort is approximately

$$\frac{\sqrt{\mathrm{Var}(E_{B_n})}}{\mathbb{E}(E_{B_n})} = \frac{s_e}{\bar{e}} \sqrt{\frac{1 - n/N}{n}}.$$

Equivalently, if $\mathrm{CV}(e) = s_e/\bar{e}$, then

$$\frac{\sqrt{\mathrm{Var}(E_{B_n})}}{\mathbb{E}(E_{B_n})} = \mathrm{CV}(e) \sqrt{\frac{1 - n/N}{n}}.$$

This decreases approximately as $1/\sqrt{n}$ for moderate $n$. A Chebyshev bound makes the concentration explicit:

$$\Pr\left(\left|E_{B_n} - n\bar{e}\right| \geq \varepsilon n\bar{e}\right) \leq \frac{\mathrm{CV}(e)^2(1 - n/N)}{n\varepsilon^2}.$$

Because $g(E)$ is smooth and monotone, a first-order Taylor expansion gives

$$g\left(E_{B_n}\right) \approx g(n\bar{e}) + g'(n\bar{e})\left(E_{B_n} - n\bar{e}\right).$$

The derivative is

$$g'(E) = \sum_{i:\lambda_i>0} \lambda_i \exp(-\lambda_i E).$$

This derivative decreases as $E$ increases, so the same amount of total-depth variation has less effect later in the accumulation curve.

Since $\mathbb{E}\left(E_{B_n} - n\bar{e}\right) = 0$, the center of the bootstrap distribution of $g\left(E_{B_n}\right)$ is approximately

$$g(n\bar{e}).$$

Also, because $g$ is monotone,

$$\mathrm{median}\{g\left(E_{B_n}\right)\} = g\{\mathrm{median}\left(E_{B_n}\right)\}.$$

When $E_{B_n}$ is concentrated, its median is close to its mean $n\bar{e}$. Therefore,

$$\mathrm{median}_{B_n}\{A_1(B_n)\} \approx g(n\bar{e}).$$

This explains why the pointwise median bootstrap $k = 1$ accumulation curve can be well approximated by an equal-depth curve, even when individual bootstrap replicates vary because they include different mixtures of low- and high-depth samples.

In this interpretation, the curve parameter $K$ absorbs average effective sequencing depth. Therefore, $K$ is a protocol-scale parameter, not a purely biological parameter.

The approximation is weakest at very small $n$, when total depth is not yet concentrated, and for study-level accumulation if study sizes or sequencing depths are extremely heterogeneous.

##### 4D. Why $k \geq 2$ accumulation requires a different model

For $k \geq 2$, an SGB must be detected repeatedly. This is not the same as a first-detection problem.

Assume first that all sampling units have equal depth, so SGB $i$ has the same per-unit detection probability $p_i$. Then

$$X_i(n) \mid p_i \sim \mathrm{Binomial}(n, p_i).$$

The probability that SGB $i$ is detected in at least $k$ units is

$$\Pr\{X_i(n) \geq k \mid p_i\} = 1 - \sum_{r=0}^{k-1} \binom{n}{r} p_i^r (1-p_i)^{n-r}.$$

Summing over all reference SGBs,

$$A_k(n) = \sum_{i=1}^{M} \left[ 1 - \sum_{r=0}^{k-1} \binom{n}{r} p_i^r (1-p_i)^{n-r} \right].$$

For $k = 1$, this becomes

$$A_1(n) = \sum_{i=1}^{M} \{1 - (1-p_i)^n\}.$$

For $k = 2$, it becomes

$$A_2(n) = \sum_{i=1}^{M} \{1 - (1-p_i)^n - np_i(1-p_i)^{n-1}\}.$$

The two subtracted terms are the probabilities of observing the SGB zero times and exactly once. Thus, $A_2(n)$ counts only SGBs that have crossed the two-detection threshold.

The local shape near zero also differs. In a Poissonized approximation,

$$X_i(t) \mid \lambda_i \sim \text{Poisson}(\lambda_i t).$$

Then

$$\Pr\{X_i(t) \geq k\} = 1 - \exp(-\lambda_i t) \sum_{r=0}^{k-1} \frac{(\lambda_i t)^r}{r!}.$$

For small $t$,

$$\Pr\{X_i(t) \geq k\} = \frac{(\lambda_i t)^k}{k!} + O(t^{k+1}).$$

Therefore,

$$A_k(t) = \frac{t^k}{k!} \sum_i \lambda_i^k + O(t^{k+1}).$$

The first $k - 1$ derivatives at zero vanish. A Michaelis-Menten curve has nonzero initial slope, so it has the wrong local structure for $k \geq 2$. A Hill curve can be forced to mimic a $t^k$ shoulder by choosing a Hill coefficient near $k$, but it still does not encode the correct binomial-tail probabilities or guarantee that the $k = 1,2,3,5$ curves arise from a shared underlying prevalence distribution.

This is why beta-binomial and beta-mixture models are used for repeated-detection accumulation.

##### 4E. Equal-depth beta-binomial model for $k \geq 1$

The beta-binomial model assumes that non-zero-prevalence SGBs differ in their per-unit detection probabilities. Specifically,

$$p_i \mid p_i > 0 \sim \text{Beta}(\alpha, \beta).$$

The beta density is

$$f(p \mid \alpha, \beta) = \frac{p^{\alpha-1}(1-p)^{\beta-1}}{B(\alpha, \beta)}, \qquad 0 < p < 1.$$

Conditional on $p_i$,

$$X_i(n) \mid p_i \sim \text{Binomial}(n, p_i).$$

The marginal probability that a non-zero-prevalence SGB is detected exactly $r$ times is

$$q_r(n; \alpha, \beta) = \int_0^1 \binom{n}{r} p^r (1-p)^{n-r} \frac{p^{\alpha-1}(1-p)^{\beta-1}}{B(\alpha, \beta)}\, dp.$$

Combining powers of $p$ and $1-p$,

$$q_r(n; \alpha, \beta) = \binom{n}{r} \frac{1}{B(\alpha, \beta)} \int_0^1 p^{r+\alpha-1} (1-p)^{n-r+\beta-1}\, dp.$$

The integral is a beta function:

$$\int_0^1 p^{r+\alpha-1} (1-p)^{n-r+\beta-1}\, dp = B(r+\alpha, n-r+\beta).$$

Therefore,

$$q_r(n; \alpha, \beta) = \binom{n}{r} \frac{B(r+\alpha, n-r+\beta)}{B(\alpha, \beta)}.$$

Define the beta-binomial tail probability

$$T_k(n; \alpha, \beta) = 1 - \sum_{r=0}^{k-1} q_r\,(n; \alpha, \beta).$$

The expected accumulation curve is

$$A_k(n) = S_\infty T_k(n; \alpha, \beta).$$

Expanded,

$$A_k(n) = S_\infty \left[1 - \sum_{r=0}^{k-1} \binom{n}{r} \frac{B(r+\alpha, n-r+\beta)}{B(\alpha,\beta)}\right].$$

This formula works for all $k \geq 1$. For $k = 1$,

$$A_1(n) = S_\infty \left[1 - \frac{B(\alpha, n+\beta)}{B(\alpha,\beta)}\right].$$

For $k = 2$,

$$A_2(n) = S_\infty \left[1 - \frac{B(\alpha, n+\beta)}{B(\alpha,\beta)} - n\frac{B(1+\alpha, n-1+\beta)}{B(\alpha,\beta)}\right].$$

**4E.1. Special case connecting beta-binomial accumulation to Michaelis-Menten**

If

$$p_i \mid p_i > 0 \sim \mathrm{Beta}(1,\beta),$$

then

$$A_1(n) = S_\infty \frac{n}{n+\beta}.$$

Thus, Michaelis-Menten arises as the $k = 1$ beta-binomial curve under a $\mathrm{Beta}(1,\beta)$ prevalence distribution.

However, the corresponding $k > 1$ curves are not Michaelis-Menten. They are

$$A_k(n) = S_\infty \prod_{j=0}^{k-1} \frac{n-j}{n+\beta-j}.$$

For $k = 2$,

$$A_2(n) = S_\infty \frac{n(n-1)}{(n+\beta)(n+\beta-1)}.$$

This demonstrates that even when the $k = 1$ curve is exactly Michaelis-Menten, the repeated-detection curves have a different structure.

**4E.2. Rare-tail behavior and slow approach to the asymptote**

The beta-binomial model also describes how quickly the curve approaches its asymptote. Define the deficit

$$D_k(n) = S_\infty - A_k(n).$$

Under the beta-binomial model,

$$D_k(n) = S_\infty \sum_{r=0}^{k-1} \binom{n}{r} \frac{B(r+\alpha, n-r+\beta)}{B(\alpha,\beta)}.$$

For large $n$, using gamma-ratio asymptotics, this behaves as

$$D_k(n) \sim C_k n^{-\alpha},$$

where

$$C_k = S_\infty \frac{\Gamma(\alpha+\beta)}{\Gamma(\beta)} \sum_{r=0}^{k-1} \frac{(\alpha)_r}{r!}.$$

Here $(\alpha)_r$ is the rising factorial:

$$(\alpha)_r = \alpha(\alpha+1)\cdots(\alpha+r-1).$$

Therefore,

$$A_k(n) = S_\infty - C_k n^{-\alpha} + o(n^{-\alpha}).$$

This power-law approach to the asymptote is important. If $\alpha$ is small, there are many ultra-rare SGBs, and the curve may continue increasing slowly over a very long sampling range. Thus, a finite-catalogue asymptote near $M$ can be compatible with an observed curve that has not visibly saturated.

**4F. Beta-mixture model**

A single beta distribution may be too restrictive because the catalogue can contain common core SGBs, intermediate-prevalence SGBs, and a long tail of rare SGBs. To allow this while retaining a mechanistic finite-catalogue model, we use a finite mixture of beta distributions:

$$p_i \sim (1-\rho)\delta_0 + \rho \sum_{h=1}^{H} v_h \,\mathrm{Beta}(\alpha_h, \beta_h).$$

Here

$$\rho = \frac{S_\infty}{M},$$

$$v_h \geq 0,$$

and

$$\sum_{h=1}^{H} v_h = 1.$$

The point mass $(1-\rho)\delta_0$ represents reference SGBs with zero detectability under the sampling protocol. The non-zero-prevalence mixture represents detectable SGBs. The asymptote is

$$S_\infty = M\rho.$$

For component $h$, define

$$q_{r,h}(n) = \binom{n}{r}\frac{B(r+\alpha_h, n-r+\beta_h)}{B(\alpha_h,\beta_h)}.$$

The mixture probability among detectable SGBs is

$$q_r^{\text{mix}}(n) = \sum_{h=1}^{H} v_h\, q_{r,h}(n).$$

The mixture tail is

$$T_k^{\text{mix}}(n) = 1 - \sum_{r=0}^{k-1} q_r^{\text{mix}}(n).$$

Equivalently,

$$T_k^{\text{mix}}(n) = \sum_{h=1}^{H} v_h\, T_k(n;\alpha_h,\beta_h).$$

The expected accumulation curve is

$$A_k(n) = S_\infty T_k^{\text{mix}}(n).$$

Expanded,

$$A_k(n) = S_\infty \sum_{h=1}^{H} v_h \left[1 - \sum_{r=0}^{k-1}\binom{n}{r}\frac{B(r+\alpha_h, n-r+\beta_h)}{B(\alpha_h,\beta_h)}\right].$$

This is the equal-depth beta-mixture model used for the $k \geq 2$ supplementary analyses. The cases $H = 1,2,3$ correspond to one-, two-, and three-component non-zero-prevalence distributions.

##### 4G. Depth-aware beta-binomial and beta-mixture models for $k \geq 1$

###### 4G.1. General solution for $k \geq 1$ (for sample accumulation)

The equal-depth beta-binomial model assumes that each sampled unit has the same detection probability for a given SGB. When sequencing depth varies, the per-sample detection probability should depend on depth.

Let $e_j$ be normalized effective depth for sample $j$. Let $p_i$ be the detection probability in a unit-depth sample. A convenient depth-aware Bernoulli model is

$$q_{ij} = 1 - (1-p_i)^{e_j}.$$

This is the beta-scale analogue of the Poisson detection model $1 - \exp(-\lambda_i e_j)$.

For a subset $B$ of samples, define

$$X_i(B) = \sum_{j \in B} Y_{ij},$$

where

$$Y_{ij} \mid p_i, e_j \sim \text{Bernoulli}\{1 - (1 - p_i)^{e_j}\}.$$

Conditional on $p_i$, $X_i(B)$ is a Poisson-binomial random variable if the $e_j$'s differ. Its probability-generating function is

$$G_i(z \mid p_i, B) = \prod_{j \in B} [(1 - p_i)^{e_j} + \{1 - (1 - p_i)^{e_j}\} z].$$

The probability of exactly $r$ detections is the coefficient of $z^r$:

$$\Pr\{X_i(B) = r \mid p_i\} = [z^r] G_i(z \mid p_i, B).$$

Therefore,

$$\Pr\{X_i(B) \geq k \mid p_i\} = 1 - \sum_{r=0}^{k-1} [z^r]\, G_i(z \mid p_i, B).$$

For beta-mixture component $h$, define

$$T_{k,h}^{\text{depth}}(B) = \mathbb{E}_{p \sim \text{Beta}(\alpha_h, \beta_h)} \left[ 1 - \sum_{r=0}^{k-1} [z^r]\, G(z \mid p, B) \right].$$

Then the depth-aware beta-mixture accumulation curve is

$$A_k^{\text{depth}}(B) = S_\infty \sum_{h=1}^{H} v_h\, T_{k,h}^{\text{depth}}(B).$$

This is the general depth-aware solution for all $k \geq 1$ under the Bernoulli model above, in the sense that the Poisson-binomial coefficient is exact for any $k$; closed forms in beta functions can be found for $k = 1$ and $k = 2$, and higher $k$ requires evaluating the coefficient numerically.

##### 4G.2. Study-level depth and study-size variation

For study-level accumulation, a unit is an entire study. If study $j$ contains samples $s = 1, \ldots, m_j$, and sample $s$ in study $j$ has depth $e_{js}$, then the probability that SGB $i$ is detected somewhere in study $j$ is

$$q_{ij}^{\text{study}} = 1 - \prod_{s=1}^{m_j} \left(1 - q_i(e_{js})\right).$$

If the per-sample detection function is

$$q_i(e_{js}) = 1 - (1 - p_i)^{e_{js}},$$

then

$$q_{ij}^{\text{study}} = 1 - \prod_{s=1}^{m_j} (1 - p_i)^{e_{js}} = 1 - (1 - p_i)^{E_j},$$

where

$$E_j = \sum_{s=1}^{m_j} e_{js}.$$

Thus, in the simplest model, a study behaves like a unit with effective depth $E_j$. However, study-level accumulation should still be interpreted carefully, because studies can differ in sample size, population composition, geography, sequencing depth, and ascertainment.

**4H. Bootstrap accumulation curves and the median curve**

For each analysis, bootstrap accumulation curves were generated over a grid of sampling efforts

$$n_1, n_2, \dots, n_J.$$

Let

$$y_{b,j}$$

denote the number of accumulated SGBs in bootstrap replicate $b$ at sampling effort $n_j$, where

$$b = 1, \dots, B, \qquad j = 1, \dots, J.$$

The pointwise bootstrap median curve is

$$\tilde{y}_j = Q_{0.50}\{y_{b,j}\}.$$

The pointwise 95% bootstrap envelope is

$$q_{0.025,j} = Q_{0.025}\{y_{b,j}\}, \qquad q_{0.975,j} = Q_{0.975}\{y_{b,j}\}.$$

The median curve was used for the left column of the supplementary figures, where the goal is to show the central accumulation pattern and fitted model curve. The 95% bootstrap envelope was used to visualize the range of bootstrap accumulation trajectories. For weighted curve fitting, the pointwise scale was estimated from this 95% envelope as

$$\hat{\sigma}_j = \frac{q_{0.975,j} - q_{0.025,j}}{2z_{0.975}}, \qquad z_{0.975} = 1.96.$$

This scale estimate gives less weight to portions of the curve where bootstrap trajectories are more variable. It is a fitting weight and visualization aid; the final uncertainty interval for $S_\infty$ is defined from the empirical 2.5th and 97.5th percentiles of the bootstrap asymptote estimates.

##### 4H.1. Fitting the bootstrap-median curve

For each panel, the fitted curve shown in the left column was estimated from the pointwise bootstrap median curve. The median-curve objective was

$$\mathrm{nll}_{\mathrm{med}}(S_\infty,\eta) = \frac{1}{2}\sum_{j=1}^{J}\left[\frac{\tilde{y}_j - \mu_k(n_j;S_\infty,\eta)}{\hat{\sigma}_j}\right]^2.$$

The fitted median-curve parameters were

$$(\hat{S}_{\infty,\mathrm{med}},\hat{\eta}_{\mathrm{med}}) = \mathrm{argmin}(\mathrm{nll}(\mathrm{med}(S_\infty,\eta))).$$

Goodness of fit to the bootstrap-median curve was summarized by

$$R^2 = 1 - \frac{\sum_{j=1}^{J}[\tilde{y}_j - \mu_k(n_j;\hat{S}_{\infty,\mathrm{med}},\hat{\eta}_{\mathrm{med}})]^2}{\sum_{j=1}^{J}(\tilde{y}_j - \bar{y})^2},$$

where

$$\bar{y} = \frac{1}{J}\sum_{j=1}^{J}\tilde{y}_j.$$

This median-curve fit describes the central accumulation trajectory. It does not by itself define the 95% uncertainty interval for $S_\infty$, because the median curve does not represent the full range of bootstrap accumulation trajectories.

##### 4H.2. Bootstrap refitting and 95% asymptote intervals

To propagate uncertainty from the accumulation process into the asymptote estimate, the selected model was refit to each bootstrap replicate. For bootstrap replicate $b$, the curve-fitting objective was

$$\mathrm{nll}_b(S_\infty,\eta) = \frac{1}{2}\sum_{j=1}^{J}\left[\frac{y_{b,j} - \mu_k(n_j;S_\infty,\eta)}{\hat{\sigma}_j}\right]^2.$$

The replicate-specific asymptote estimate was

$$\hat{S}_\infty^{(b)} = \mathrm{argmin}\{\min(\mathrm{nll}_b(S_\infty,\eta)\},$$

with the catalogue constraint

$$\max(y_{b,j}) \le S_\infty \le M.$$

The empirical distribution

$$\hat{S}_\infty^{(1)}, \hat{S}_\infty^{(2)}, \dots, \hat{S}_\infty^{(B)}$$

is the bootstrap distribution of the inferred asymptote. The right column of the supplementary figures shows this distribution. The bootstrap median asymptote is

$$\hat{S}_{\infty,0.5}^{\text{boot}} = Q_{0.50}\{\hat{S}_\infty^{(b)}\},$$

and the 95% bootstrap interval is

$$\left[Q_{0.025}\{\hat{S}_\infty^{(b)}\},\ Q_{0.975}\{\hat{S}_\infty^{(b)}\}\right].$$

These 2.5th and 97.5th percentiles are the intervals used to summarize uncertainty in $S_\infty$ in the supplementary sensitivity figures. If the upper percentile reaches the catalogue cap $M$, the interval should be interpreted as boundary-constrained rather than as evidence of zero upper uncertainty.

This bootstrap-refitting step is essential. If only the pointwise median curve were fitted, the resulting asymptote interval would reflect the curvature of one smoothed curve and would ignore the range of plausible accumulation trajectories observed across bootstrap replicates. By refitting each bootstrap curve, the asymptote interval reflects both uncertainty in the accumulation trajectory and uncertainty in extrapolating that trajectory to the catalogue-constrained asymptote.

##### 4H.3. Bootstrap refitting for beta-mixture repeated-detection curves

For the beta-mixture repeated-detection panels, the component tail functions are

$$T_{k,h}(n) = 1 - \sum_{r=0}^{k-1} \binom{n}{r} \frac{B(r + \alpha_h, n - r + \beta_h)}{B(\alpha_h, \beta_h)}.$$

A general beta-mixture curve is

$$\mu_k(n) = S_\infty \sum_{h=1}^{H} v_h\, T_{k,h}(n).$$

Equivalently, define non-negative component masses

$$c_h = S_\infty v_h.$$

Then

$$\mu_k(n) = \sum_{h=1}^{H} c_h\, T_{k,h}(n), \qquad S_\infty = \sum_{h=1}^{H} c_h.$$

This parameterization is useful for bootstrap refitting because each bootstrap replicate can be fitted by non-negative weighted least squares over the component masses. For replicate $b$, the component masses are estimated by

$$\hat{\mathbf{c}}^{(b)} = \arg\min_{c_1,\dots,c_H \geq 0} \frac{1}{2}\sum_{j=1}^{J}\left[\frac{y_{b,j} - \sum_{h=1}^{H} c_h\, T_{k,h}(n_j)}{\hat{\sigma}_j}\right]^2,$$

with

$$\sum_{h=1}^{H} c_h \leq M.$$

The bootstrap asymptote estimate is

$$\hat{S}_\infty^{(b)} = \sum_{h=1}^{H} \hat{c}_h^{(b)}.$$

The normalized bootstrap mixture weights are

$$\hat{v}_h^{(b)} = \frac{\hat{c}_h^{(b)}}{\hat{S}_\infty^{(b)}}$$

when $\hat{S}_\infty^{(b)} > 0$. This refitting step allows each bootstrap curve to imply its own asymptote and mixture composition, while preserving the finite-catalogue beta-mixture structure used for the repeated-detection curves.

##### 4H.4. Bootstrap support profiles shown in the middle panels

The middle column of the supplementary figures visualizes support for possible values of $S_\infty$. This profile is meant to be read together with the bootstrap histogram in the right column. The histogram gives the 95% bootstrap interval for the asymptote. The support profile shows how rapidly the curve fit deteriorates as $S_\infty$ is moved away from values preferred by the bootstrap curves.

For a candidate asymptote $s$, the replicate-specific profiled objective is

$$\mathrm{nll}_{\mathrm{prof},b}(s) = min(\,\mathrm{nll}_b(s, \eta)).$$

The subscript "prof" indicates that this is a profile negative log-likelihood: $s$ is held fixed, while all other fitted parameters are optimized.

The replicate-specific relative support score is

$$\Delta_b(s) = 2\left[\mathrm{nll}_{\mathrm{prof},b}(s) - min(\,\mathrm{nll}_{\mathrm{prof},b}(s'))\right].$$

A value of $\Delta_b(s) = 0$ indicates the best-supported asymptote for bootstrap replicate $b$. Larger values indicate that the replicate is less compatible with that candidate asymptote.

To display a single support curve while retaining bootstrap variability, the replicate-specific support profiles can be aggregated as a bootstrap mixture support,

$$L_{\text{boot}}(s) = \frac{1}{B}\sum_{b=1}^{B} \exp\left[-\frac{1}{2}\Delta_b(s)\right],$$

and rescaled to a relative support score,

$$\Delta_{\text{boot}}(s) = -2\log\left\{\frac{L_{\text{boot}}(s)}{\max(L_{\text{boot}}(s'))}\right\}.$$

This aggregation preserves support for values of $S_\infty$ that are favored by a substantial subset of bootstrap replicates. It is preferable to treating all bootstrap curves as independent observations, which would artificially overstate precision.

When a pointwise bootstrap band for the support profile is shown, it is calculated from the 2.5$^{\text{th}}$ and 97.5$^{\text{th}}$ percentiles of the replicate-specific profiles,

$$[Q_{0.025}\{\Delta_b(s)\},\ Q_{0.975}\{\Delta_b(s)\}],$$

at each candidate value of $s$. Thus, the support-profile band is also a 95% bootstrap band.

The horizontal reference line in the support-profile panels is

$$\Delta = 3.84.$$

This is the usual one-parameter 95% likelihood-ratio reference value. In this context it should be interpreted as a visual guide to relative support, not as a replacement for the bootstrap percentile interval. The reported uncertainty in $S_\infty$ is taken from the empirical 2.5$^{\text{th}}$ and 97.5$^{\text{th}}$ percentiles of $\hat{S}_\infty^{(b)}$. For readability, support profiles may be truncated at $\Delta = 20$ in the plotted figures.

**4H.5. Why the bootstrap procedure is not a stacked likelihood**

The bootstrap accumulation curves are not independent datasets. They are resampled summaries of the same underlying sample-by-SGB detection matrix. Therefore, we did not form a single likelihood by stacking all bootstrap curves as though they were independent observations. Such a likelihood would have the form

$$\sum_{b=1}^{B}\sum_{j=1}^{J}\left[\frac{y_{b,j} - \mu_k(n_j; S_\infty, \eta)}{\hat{\sigma}_j}\right]^2,$$

and would greatly overstate the precision of the asymptote because it would count the same underlying data many times.

Instead, each bootstrap curve was refit separately, producing one asymptote estimate per bootstrap replicate. The resulting distribution of $\hat{S}_\infty^{(b)}$ is the inferential object used for the 95% bootstrap interval. This is why the supplementary figures show both the support profile and the bootstrap histogram: the profile describes relative fit as a function of $S_\infty$, while the histogram directly displays bootstrap uncertainty in the preferred asymptote.

**4I. Interpretation of boundary fits**

If $\hat{S}_\infty$ lies at the catalogue cap $M$, this does not prove that the asymptote is known exactly. It means that, under the chosen model, the likelihood is maximized by assigning positive detectability to essentially all reference SGBs.

There are two possible explanations. The first is biological: nearly all reference SGBs may truly be detectable under the sampling protocol, though some may be extremely rare. The second is model misspecification: the positive-detectability distribution may be too simple, causing the model to push $S_\infty$ to the boundary in order to explain the low-count tail.

For this reason, boundary fits should be interpreted using all of the following:

$$\hat{S}_\infty,$$

$$\Delta(M),$$

and the curve diagnostics $R^2$ and RMSE.

If a model with $H = 2$ or $H = 3$ fits the plotted accumulation curve well and still gives $\hat{S}_\infty \approx M$ with small $\Delta(M)$, then the data support a near-catalogue asymptote under a model that also describes the observed accumulation pattern. If increasing model flexibility moves $\hat{S}_\infty$ away from $M$, the one-component boundary result was likely a model artifact. If the profile remains broad, the data are compatible with a near-catalogue asymptote but do not uniquely identify it.

**4J. Summary of model usage**

For the main $k = 1$ figure, we use Hill-type Michaelis-Menten and Richards curves as simple empirical first-detection distibution functions. This is justified because ordinary SGB accumulation is a first-detection problem. Michaelis-Menten and generalized Michaelis-Menten also have explicit derivations from exponential and gamma detectability distributions.

For $k \geq 2$, we use beta-binomial or beta-mixture models because repeated-detection accumulation is a binomial-tail problem. These models preserve the finite-catalogue asymptote and make the $k = 1,2,3,5$ curves coherent with an underlying distribution of SGB prevalences or detectabilities. The values of $k$ shown are a sparse ladder rather than a complete series: each step roughly doubles the evidence required for a detection, so intermediate values add rows without adding information, and the ladder stops where the remaining catalogue is too small for the asymptote to be identified (there are rather few admissible non-industrial studies, which is why we do not extend the search further).

When sequencing depth varies, the equal-depth models are approximations. For $k = 1$, the median bootstrap curve can still be close to an equal-depth curve because the model depends mainly on total effective depth, which concentrates around $n\bar{e}$ in random subsets. For $k \geq 2$, depth-aware beta-mixture Poisson-binomial models are the more general mathematical description.

Asymptote uncertainty is assessed by profiling $S_\infty$. The same framework can be used for empirical curve fits to bootstrap medians and for catalogue-level beta-binomial or beta-mixture likelihoods. The key reported quantities are the median fitted asymptote $\hat{S}_\infty$, its bootstrap replicates, and the profile interval $\Delta(M)$ representing the combined likelihood of the actual asymptote $\hat{S}_\infty$ given the boostrapped asymptotes.

##### 4K. Negative control: a chimpanzee catalogue does not saturate

The accumulation analyses in this note conclude that pooled human samples recover close to the entire reference catalogue, and therefore that rarely-detected taxa are rare rather than absent. That inference depends on the detection rule being specific: a rule that determined taxa as being present too readily would produce saturation regardless of biology. We therefore repeated the analysis against a catalogue of 1,682 chimpanzee-derived SGBs (*103*), using the same reads, the same ≥10% breadth-of-coverage detection rule, the same bootstrap pooling scheme and the same 4-parameter Richards fit.

A subset of the cohort was used: the 3,887 samples (24 studies, 15 countries) processed against the chimpanzee catalogue, of which 3,802 are industrial. Within this subset the fitted asymptote reached only 762 SGBs, 45.3% of the chimpanzee catalogue (95% bootstrap interval 569–1,241) compared with 99.5–100% for each of the three human catalogues under the identical fit (**Table S5**).

At a stricter ≥30% breadth threshold, the chimpanzee asymptote remains far below the catalogue, at 26.6%. Only 492 of the 1,682 chimpanzee SGBs (29.3%) were detected in any industrial sample, and the read-mapping rate against the chimpanzee catalogue was 8.3% versus 38.8% for the Carter catalogue on the same samples.

The accumulation framework therefore distinguishes "rare in humans" from "not a human taxon": applied to human catalogues it saturates, and applied to a chimpanzee catalogue over the same reads it does not. The magnitude is consistent with (*103*), who report that ~44% of co-diversifying gut bacterial clades found in African apes lack human representatives (~54% for industrialized populations).

**Table S5. Reference-catalogue recovery under sample pooling, with the chimpanzee negative control.** For each reference catalogue, the number of SGBs detected when all industrial samples are pooled at the ≥10% breadth-coverage threshold, and the asymptote of the accumulation curve. Asymptotes are the 4-parameter Richards fit to the bootstrap-median curve in log-N space, weighted by the bootstrap spread and bounded above by the catalogue size (the identical fit used for **Fig. 3**) with 95% bootstrap intervals in brackets. The chimpanzee catalogue comprises 1,682 bins assembled from wild chimpanzee gut metagenomes and is processed from the same reads, through the same pipeline, under the same detection rule as the three human catalogues. Human catalogues are recovered essentially completely under pooling, whereas the chimpanzee catalogue is not, confirming that saturation reflects genuine presence of these taxa in human guts rather than a permissive detection rule. Chimpanzee counts are for the 3,887-sample subset processed against that catalogue, of which 3,802 are industrial; the non-industrial chimpanzee series is not reported because 85 samples are not sufficient to constrain the fit.

| Catalogue | Catalogue SGBs | Industrial Samples Pooled | SGBs detected | Observed | Fitted asymptote [95% CI] | % of catalogue |
|---|---|---|---|---|---|---|
| Combined (Carter + Leviatan) | 5,081 | 13,977 | 4,233 | 83.3% | 5,081 [4,833–5,081] | 100% |
| Carter | 3,971 | 13,951 | 3,289 | 82.8% | 3,952 [3,745–3,971] | 99.5% |
| Leviatan | 3,594 | 13,966 | 3,326 | 92.5% | 3,594 [3,594–3,594] | 100% |
| Chimpanzee | 1,682 | 3,802 | 492 | 29.3% | 762 [569–1,241] | 45.3% |

#### Supplementary Text 5. Construction and interpretation of the VANISH/BloSSUM, VANISH$^+$/BloSSUM$^+$, and VANISH$^{H+}$/BloSSUM$^{H+}$ taxon lists

##### 5A. Why multiple taxon-list families are needed

The manuscript uses three related but non-identical taxon-list families because they answer three different questions. The original VANISH and BloSSUM lists were imported from prior Hadza-centered work (*7*) and are retained to preserve continuity with the existing literature. VANISH$^+$ and BloSSUM$^+$ are defined de novo in the present meta-analysis to capture the taxa most differentially prevalent between industrial and non-industrial microbiomes in the full dataset analyzed here. VANISH$^{H+}$ and BloSSUM$^{H+}$ are defined analogously, but with the industrial reference restricted to healthy industrial individuals, to ask whether the industrialization contrast changes when disease-enriched industrial cohorts are excluded. These list families should therefore not be interpreted as interchangeable.

**Fig. 4** is supported primarily by **Figs. S21–S22**, by **Table S3** for the parent cohort definition, and by **Tables S6–9** for overlap, detection, pooled absence, rare taxon candidates and **Data S4** for full list membership.

##### 5B. Original VANISH and BloSSUM lists

We retained the published Carter et al. VANISH and BloSSUM lists unchanged for comparison to prior work. These lists contain 124 VANISH taxa and 63 BloSSUM taxa and were originally defined in a Hadza-versus-industrial framework (*7*). In the present study, these lists serve two main purposes. First, they provide a literature-anchored benchmark for evaluating claims of local extinction or persistence. Second, they allow us to quantify how much our empirically derived lists recapitulate, refine, or depart from prior Hadza-centered definitions. Because the original lists were optimized to distinguish Hadza hunter-gatherers from industrial cohorts, they do not isolate the industrial transition from broader differences across the non-industrial spectrum (e.g., the agricultural revolution).

##### 5C. Construction of VANISH+ and BloSSUM+

To identify the taxa most strongly associated with industrialization in the present dataset, we compared all industrial samples with all non-industrial samples, where the latter comprised both hunter-gatherer and agro-pastoralist populations. For each SGB $j$, per-sample presence was defined at the standard detection threshold used in the differential-prevalence analysis (≥10% breadth coverage unless otherwise noted in sensitivity analyses). Because industrial and non-industrial samples differ systematically in sequencing depth, raw prevalence contrasts are biased. We therefore applied geometric-mean-target weighting (GMW) to reweight the two lifestyle groups toward a common sequencing-depth target distribution before estimating taxon prevalences.

For each taxon $j$, weighted prevalence in group $g$ was computed as:

$$\hat{p}_{g,j} = \frac{\sum_i w_i \, \mathbf{1}(x_{ij} \geq \tau)}{\sum_i w_i}$$

where $x_{ij}$ is the breadth coverage of taxon $j$ in sample $i$, $\tau$ is the detection threshold, and $w_i$ is the GMW importance weight for sample $i$. We then ranked all taxa by the weighted prevalence difference

$$\Delta p_j = \hat{p}_{Industrial,j} - \hat{p}_{Non\text{-}Industrial,j}.$$

Taxa with the most negative values of $\Delta p_j$ were designated VANISH$^+$, and taxa with the most positive values were designated BloSSUM$^+$. By construction, each list contained 100 taxa: the 100 most non-industrial-enriched taxa and the 100 most industrial-enriched taxa, respectively.

The statistical test associated with this ranking used weighted 2×2 present/absent tables, in which each sample contributed its GMW weight rather than a unit count. The resulting p-values were used for visualization and descriptive calibration in volcano plots, but statistical significance was not used as the membership criterion. This fixed-size rank-based definition was chosen so that list size would not depend on an arbitrary p-value threshold or on differences in power across taxa with different baseline prevalences.

List membership is robust to repeat sampling of the same hosts. Rebuilding the lists with one sample per host retained 97 of 100 VANISH$^+$ (and VANISH$^{H+}$) taxa and 94 of 100 BloSSUM$^+$ (and BloSSUM$^{H+}$) taxa. The rank of all 5,081 SGBs was almost unchanged (Spearman $\rho$ = 0.994–0.996), and every taxon that changed status lay within about one percentage point in prevalence of the rank-100 cut-off. The lists were as stable when bootstrap resamples drew whole hosts as when they drew individual samples (expected overlap with the published list 95.8–97.6 versus 96.2–97.7 taxa), so treating repeat samples as independent does not inflate their apparent stability.

The industrial versus non-industrial parent cohort used to define VANISH$^+$/BloSSUM$^+$ is given in **Table S3**.

##### 5D. VANISH$^+$/BloSSUM+ are differential-prevalence lists, not rarity lists

An important conceptual point is that VANISH$^+$ and BloSSUM$^+$ are not defined by rarity or absence. They are defined by ranked differential prevalence between industrial and non-industrial populations after correcting for sequencing-depth imbalance. As a result, many VANISH$^+$ taxa remain detectable at moderate prevalence in industrial populations, and many BloSSUM$^+$ taxa remain detectable in non-industrial populations. This is expected: these lists capture the taxa most shifted across the industrial/non-industrial contrast, not the taxa closest to extinction. For this reason, the degree to which a taxon belongs to VANISH$^+$ should not be interpreted as a direct measure of "missingness," but rather as a measure of how strongly that taxon is depleted in industrial populations relative to the non-industrial reference.

##### 5E. Relationship to abundance

The same taxa were also visualized on CLR-abundance volcano plots to ask whether the prevalence-defined extremes show concordant abundance shifts. These abundance plots are descriptive and are not used to determine membership in VANISH$^+$ or BloSSUM$^+$. Unless otherwise stated, references to VANISH$^+$ and BloSSUM$^+$ in the main text therefore refer to prevalence-defined lists.

##### 5F. Construction of VANISH$^{H+}$ and BloSSUM$^{H+}$

To generate a health-anchored version of the industrialization contrast, we repeated the same GMW-weighted prevalence procedure after restricting the industrial reference group to samples labeled "healthy." Non-industrial samples were unchanged. The 100 most non-industrial-enriched taxa from this comparison were designated **VANISH$^{H+}$**, and the 100 most healthy-industrial-enriched taxa were designated **BloSSUM$^{H+}$**. In each case, the H superscript indicates that the industrial reference set is restricted to healthy individuals; it does not imply that these taxa are themselves health-associated taxa.

The H+-restricted lists are used primarily in the health-association analyses described in **Supplementary Text 6** and **Figs. S25–S30**.

##### 5G. Why the H+ lists are useful

The H+ lists address a specific concern: industrial cohorts in public metagenomic repositories are enriched for disease-focused case-control studies, so some industrial/non-industrial differences could reflect disease composition rather than industrialization per se. By redefining the industrial anchor using only healthy industrial samples, the H+ lists ask which taxa distinguish non-industrial microbiomes from a comparatively healthier industrial baseline. They are therefore most useful in the health-association sections of the manuscript, where they serve as a robustness check on the broader VANISH$^{+}$/BloSSUM$^{+}$ definitions.

##### 5H. Interpreting overlap and non-overlap among list families

Partial overlap between the original and empirically derived lists is expected and biologically informative. The original VANISH/BloSSUM lists were defined in a Hadza-centered comparison, whereas VANISH$^{+}$/BloSSUM$^{+}$ were defined from the broader industrial versus non-industrial contrast in the present dataset, and the VANISH$^{H+}$/BloSSUM$^{H+}$ lists further exclude unhealthy industrial samples from the industrial reference group. Taxa shared across list families represent robust lifestyle-associated signals that reflect the component of change between hunter-gatherer and industrial microbiomes that is attributable to industrialization. Taxa unique to the original VANISH/BloSSUM lists are more likely to represent taxa that either distinguish the Hadza hunter-gatherers or consistent changes in prevalence since the agricultural revolution. Taxa unique to VANISH$^{+}$/BloSSUM$^{+}$ (or H+ lists) are more likely to represent taxa that changed prevalence only with industrialization. **Table S6** quantifies these overlaps. VANISH$^{+}$ shares 40 of its 100 taxa with the 124 original VANISH taxa (Jaccard index 0.22) and 96 with VANISH$^{H+}$ (0.92); BloSSUM$^{+}$ shares 38 of its 100 taxa with the 63 original BloSSUM taxa (0.30) and 82 with BloSSUM$^{H+}$ (0.69). The + and H+ lists therefore differ in only a few taxa, whereas both depart substantially from the original lists.

**Table S6. Pairwise overlap between VANISH and BloSSUM taxon-list families. $n_A/n_B$: number of taxa in each list.** Overlap: taxa shared by both lists. A only / B only: taxa unique to each list. Jaccard $= \text{overlap}/(n_A + n_B - \text{overlap})$. Column shading follows the list-family palette used throughout the text. The VANISH$^{+}$/ VANISH$^{H+}$ and BloSSUM$^{+}$/ BloSSUM$^{H+}$ taxa only differ in a small fraction of their SGBs, reflecting the fact that the majority of the variation is driven by industrialization and not health status.

| Family | List A | List B | nA | nB | Overlap | A only | B only | Jaccard |
|---|---|---|---|---|---|---|---|---|
| **VANISH** | original VANISH | VANISH$^{+}$ | 124 | 100 | 40 | 84 | 60 | 0.2174 |
| **VANISH** | VANISH$^{+}$ | VANISH$^{H+}$ | 100 | 100 | 96 | 4 | 4 | 0.9231 |
| **VANISH** | original VANISH | VANISH$^{H+}$ | 124 | 100 | 42 | 82 | 58 | 0.2308 |
| **BloSSUM** | original BloSSUM | BloSSUM$^{+}$ | 63 | 100 | 38 | 25 | 62 | 0.304 |
| **BloSSUM** | BloSSUM$^{+}$ | BloSSUM$^{H+}$ | 100 | 100 | 82 | 18 | 18 | 0.6949 |

| Family | List A | List B | nA | nB | Overlap | A only | B only | Jaccard |
|---|---|---|---|---|---|---|---|---|
| **BloSSUM** | original BloSSUM | BloSSUM$^{H+}$ | 63 | 100 | 39 | 24 | 61 | 0.3145 |

#### 5I. Which list is primary for which question

We use the original VANISH/BloSSUM lists for continuity with prior literature and for evaluating whether previously nominated "missing" taxa persist under deep pooling. We use VANISH$^{+}$/BloSSUM$^{+}$ as the primary lists for quantifying the industrial versus non-industrial contrast in the present dataset. We use VANISH$^{H+}$/BloSSUM$^{H+}$ as a health-oriented robustness set for the within-industrial disease analyses. Concordance across these list families is interpreted as increased robustness of the underlying biological signal rather than as evidence that the lists are redundant.

#### 5J. Pooled detection of the list taxa

**Tables S7–S9** report how readily the list taxa are recovered under pooling: detection statistics across detection thresholds (**Table S7**), the original VANISH and BloSSUM taxa that remain undetected after pooling (**Table S8**), and the VANISH$^{+}$ and BloSSUM$^{+}$ taxa found in no more than three samples (**Table S9**).

**Table S7. Detection statistics across detection thresholds for the VANISH and BloSSUM taxa.** Typical SGB median N: median sample count across SGBs at each threshold. det.: fraction of list taxa detected (pooled count / list size). Med N: median sample count among detected taxa. Min det.: number of samples in which the rarest SGB is detected (>10% breadth coverage threshold). Column shading follows list-family palette.

| Threshold | Median N to detect SGB | | original VANISH | | | VANISH$^{+}$ | | | original BloSSUM | | | BloSSUM$^{+}$ (pooled / 100) | | |
|---|---|---|---|---|---|---|---|---|---|---|---|---|---|---|
| | Ind | NI | det. | Med N | Min det. | det. | Med N | Min det. | det. | Med N | Min det. | det. | Med N | Min det. |
| **10% breadth** | 485 | 95 | 115/124 | 192 | 0 | 100/100 | 23 | 21 | 63/63 | 8 | 13 | 100/100 | 10 | 19 |
| **30% breadth** | 1211 | 284 | 100/124 | 669 | 0 | 99/100 | 42 | 0 | 63/63 | 12 | 11 | 100/100 | 22 | 8 |
| **50% breadth** | 2422 | 710 | 77/124 | 2422 | 0 | 96/100 | 81 | 0 | 63/63 | 23 | 2 | 100/100 | 44 | 1 |
| **300 kb** | 646 | 135 | 111/124 | 220 | 0 | 100/100 | 24 | 10 | 63/63 | 7 | 12 | 100/100 | 10 | 15 |
| **500 kb** | 1077 | 237 | 105/124 | 360 | 0 | 100/100 | 30 | 1 | 63/63 | 9 | 12 | 100/100 | 13 | 11 |

**Table S8. Pooled absences among original VANISH and BloSSUM taxa across detection thresholds.**

| Threshold | Original VANISH in Industrial (pooled / 124) | | Original BloSSUM in Non-Industrial | |
|---|---|---|---|---|
| | det. | Pooled-missing taxa | det. | Pooled-missing taxa |
| **10% breadth** | 115/124 | GCA_934408725.1 (g__Marvinbryantia); GCA_934533025.1 (g__Gemmiger); GCA_934630535.1 (g__Senegalimassilia); GCA_934643435.1 (g__Collinsella); GCA_934654835.1 (g__Collinsella); GCA_934675635.1 (s__Treponema_D sp900541995); GCA_934692455.1 (g__UBA636); GCA_934717755.1 (g__Bulleidia); GCA_934725085.1 (g__UBA636) | 63/63 | none |

| Threshold | Original VANISH in Industrial (pooled / 124) | | Original BloSSUM in Non-Industrial | |
|---|---|---|---|---|
| | det. | Pooled-missing taxa | det. | Pooled-missing taxa |
| **30% breadth** | 100/124 | GCA_934196725.1 (g__Prevotella); GCA_934234275.1 (g__Phascolarctobacterium_A); GCA_934344725.1 (g__Agathobacter); GCA_934359255.1 (g__Ruminococcus_A); GCA_934408725.1 (g__Marvinbryantia); GCA_934501215.1 (s__Gemmiger variabilis_A); GCA_934533025.1 (g__Gemmiger); GCA_934589705.1 (g__CAG-110); GCA_934630535.1 (g__Senegalimassilia); GCA_934643285.1 (g__UMGS1773); GCA_934643435.1 (g__Collinsella); GCA_934647175.1 (g__SFFH01); GCA_934654835.1 (g__Collinsella); GCA_934660915.1 (g__Olsenella_E); GCA_934675635.1 (s__Treponema_D sp900541995); GCA_934692455.1 (g__UBA636); GCA_934717755.1 (g__Bulleidia); GCA_934725085.1 (g__UBA636); GCA_934725555.1 (g__Holdemanella); GCA_934747155.1 (g__Bulleidia); GCA_934755155.1 (g__Holdemanella); Rep_1226 (s__Catenibacterium sp900764565); Rep_1862 (s__Phascolarctobacterium_A sp934190775); Rep_2762 (s__UMGS124 sp902464015) | 63/63 | none |
| **50% breadth** | 77/124 | GCA_934196725.1 (g__Prevotella); GCA_934229735.1 (s__Prevotella sp900551985); GCA_934234275.1 (g__Phascolarctobacterium_A); GCA_934292805.1 (g__Roseburia); GCA_934317185.1 (g__Anaerobutyricum); GCA_934344725.1 (g__Agathobacter); GCA_934359255.1 (g__Ruminococcus_A); GCA_934373285.1 (g__Marvinbryantia); GCA_934394795.1 (s__Marvinbryantia sp900544685); GCA_934401805.1 (s__Acetatifactor sp003447295); GCA_934408725.1 (g__Marvinbryantia); GCA_934427075.1 (s__UBA2882 sp900549665); GCA_934501215.1 (s__Gemmiger variabilis_A); GCA_934503275.1 (g__Faecalibacterium); GCA_934512105.1 (g__Faecalibacterium); GCA_934524985.1 (g__Faecalibacterium); GCA_934525325.1 (g__Faecalibacterium); GCA_934526095.1 (g__Faecalibacterium); GCA_934532365.1 (g__Faecalibacterium); GCA_934533025.1 (g__Gemmiger); GCA_934539245.1 (g__Faecalibacterium); GCA_934561555.1 (g__Faecalibacterium); GCA_934561575.1 (g__Faecalibacterium); GCA_934589705.1 (g__CAG-110); GCA_934589835.1 (g__CAG-83); GCA_934591815.1 (s__CAG-83 sp900545495); GCA_934630535.1 (g__Senegalimassilia); GCA_934643285.1 (g__UMGS1773); GCA_934643435.1 (g__Collinsella); GCA_934647175.1 (g__SFFH01); GCA_934654835.1 (g__Collinsella); GCA_934660915.1 (g__Olsenella_E); GCA_934675635.1 (s__Treponema_D sp900541995); GCA_934692455.1 (g__UBA636); GCA_934700615.1 (g__RC9); GCA_934717755.1 (g__Bulleidia); GCA_934725085.1 (g__UBA636); GCA_934725555.1 (g__Holdemanella); GCA_934746575.1 (s__Holdemanella sp003436425); GCA_934747155.1 (g__Bulleidia); GCA_934755155.1 (g__Holdemanella); Rep_1226 (s__Catenibacterium sp900764565); Rep_1862 (s__Phascolarctobacterium_A sp934190775); Rep_2282 (s__Mediterraneibacter sp934309345); Rep_2762 (s__UMGS124 sp902464015); Rep_3076 (s__Faecalibacterium sp900772565); Rep_3080 (s__Faecalibacterium tardum) | 63/63 | none |
| **300 kb** | 111/124 | GCA_934234275.1 (g__Phascolarctobacterium_A); GCA_934408725.1 (g__Marvinbryantia); GCA_934533025.1 (g__Gemmiger); GCA_934630535.1 (g__Senegalimassilia); GCA_934643285.1 (g__UMGS1773); GCA_934643435.1 (g__Collinsella); GCA_934654835.1 (g__Collinsella); GCA_934675635.1 (s__Treponema_D sp900541995); GCA_934692455.1 (g__UBA636); GCA_934717755.1 (g__Bulleidia); GCA_934725085.1 (g__UBA636); GCA_934725555.1 (g__Holdemanella); Rep_1226 (s__Catenibacterium sp900764565) | 63/63 | none |
| **500 kb** | 105/124 | GCA_934234275.1 (g__Phascolarctobacterium_A); GCA_934408725.1 (g__Marvinbryantia); GCA_934526095.1 (g__Faecalibacterium); GCA_934533025.1 (g__Gemmiger); GCA_934589705.1 (g__CAG-110); GCA_934630535.1 (g__Senegalimassilia); GCA_934643285.1 (g__UMGS1773); GCA_934643435.1 (g__Collinsella); GCA_934647175.1 (g__SFFH01); GCA_934654835.1 (g__Collinsella); GCA_934675635.1 (s__Treponema_D sp900541995); GCA_934692455.1 (g__UBA636); GCA_934717755.1 (g__Bulleidia); GCA_934725085.1 (g__UBA636); GCA_934725555.1 (g__Holdemanella); GCA_934747155.1 (g__Bulleidia); GCA_934755155.1 (g__Holdemanella); Rep_1226 (s__Catenibacterium sp900764565); Rep_1862 (s__Phascolarctobacterium_A sp934190775) | 63/63 | none |

**Table S9.** Almost all VANISH+ and all BloSSUM+ SGBs were found in every sample but the following listed SGBs were not found in >3 samples.

| Threshold | VANISH+ in Industrial (pooled / 100) | | | BloSSUM+ in Non-Industrial (pooled / 100) | | |
|---|---|---|---|---|---|---|
| | det. | n candidates (≤3 hard det.) | Rare taxa | det. | n candidates (≤3 hard det.) | Rare taxa |
| 10% breadth | 100/100 | 0 | none | 100/100 | 0 | none |
| 30% breadth | 99/100 | 3 | GCA_934196725.1 (g__Prevotella); GCA_934215355.1 (s__Dialister sp934178345); GCA_934229735.1 (s__Prevotella sp900551985) | 100/100 | 0 | none |
| 50% breadth | 96/100 | 8 | GCA_934196725.1 (g__Prevotella); GCA_934215355.1 (s__Dialister sp934178345); GCA_934229735.1 (s__Prevotella sp900551985); MGYG000000707 (s__Prevotella sp900546575); MGYG000003347 (s__Prevotella sp900765465); MGYG000004456 (s__Prevotella sp900546535); Rep_236 (s__Prevotella sp900557035) GCA_934401805.1 (s__Acetatifactor hominis) | 100/100 | 3 | MGYG000002480 (s__Adlercreutzia rubneri); MGYG000004797 (s__Phocaeicola sartorii); Rep_3401 (s__Acutalibacter ornithocaccae) |
| 300 kb | 100/100 | 0 | none | 100/100 | 0 | none |
| 500 kb | 100/100 | 1 | GCA_934196725.1 (g__Prevotella) | 100/100 | 0 | none |

#### Supplementary Text 6. Within-industrial health association analyses: study-stratified, depth-corrected, prevalence-referenced testing

##### 6A. Overview and analytical problem

**Supplementary Text 6** describes the within-industrial health analyses underlying **Fig. 5**. These comparisons are vulnerable to three confounds acting at once: sequencing-depth imbalance between cases and controls, heterogeneity between studies, and baseline-prevalence effects that constrain how far a taxon can move between cases and controls. We address them in that order. Depth is equalized within each study stratum by geometric-mean-target weighting (GMW); study heterogeneity is handled by stratifying on study and combining strata by the Mantel-Haenszel construction; and baseline prevalence is handled by referencing every taxon to a fitted expectation for taxa of the same commonness, rather than to the raw pool of non-list taxa.

Two naming conventions are fixed here and used throughout. Cochran-Mantel-Haenszel (CMH) names the stratified test of conditional independence (*14*, *15*). Mantel-Haenszel (MH) names the two stratified estimators we take from the same tables: the common odds ratio (*15*) and the common risk difference (*104*, *105*). We write GMW-CMH for the weighted stratified construction as a whole, CMH when we mean the test, and MH when we mean an estimator, naming the scale in each case.

The key object in the CMH test is the per-study 2×2 table of taxon detection versus absence by control versus case status:

$$\begin{pmatrix} & \text{Present} & \text{Absent} \\ \text{Healthy} & a_k & b_k \\ \text{Unhealthy} & c_k & d_k \end{pmatrix}$$

The primary supporting materials for this note are **Tables S3**, **S10**, and **S11** and **Figs. S25–S30**.

##### 6B. Health cohorts, eligible strata, and primary estimand

Health associations were evaluated within industrial cohorts only. 4 disease/control comparisons cleared the threshold of 3 studies with at least 5 controls and 5 cases: obesity, glucose intolerance, IBD, and colorectal cancer/adenoma. For all CMH-based analyses, the unit of adjustment was the study stratum**:** each study contributed an independent case-control comparison, and only strata with at least 5 controls and 5 cases were eligible for inclusion. Raw sequencing depth differs across health groups before correction (**Fig. S23**), making within-study depth equalization necessary before any taxon-level health association is interpreted.

**Table S10** summarizes the size and quality of these comparisons. Cohort size ranged from 979 hosts in IBD to 7,822 hosts for obesity. For each condition, we report the number of contributing studies, the number of controls and cases, a matched-depth diagnostic (med_LD_diff), and the effective sample size fraction after GMW reweighting ($ESS_{frac}$). Across the four plotted conditions, the GMW ESS fractions remained high (0.842–0.958), indicating that depth equalization was generally achieved without strong loss of information. Conditions supported by fewer than three studies were excluded from **Fig. 5D–E** and **Tables S10–S11**.

Broad health-group depth and richness diagnostics are shown in **Figs. S23–S24**.

The primary taxon-level estimand in the health analyses is the Mantel–Haenszel common risk difference in taxon detection between controls and cases, after reweighting both groups to a shared geometric-mean-target depth distribution within each study stratum, with the Sato standard error (*105*). We write it $\mathrm{rd}_{\mathrm{MH}} = \mathrm{prev}_{\mathrm{control}} - \mathrm{prev}_{\mathrm{case}}$, so that positive values denote taxa that are more prevalent in controls and therefore depleted in disease. The Mantel–Haenszel common log-odds ratio is retained and reported alongside it. A second difference is also stored for each taxon, obtained by pooling every control across studies, pooling every case, and then subtracting; this is the quantity we refer to as $\mathrm{prev}_{\mathrm{diff}}$. It is not a within-study contrast and is used for no inference. Because studies differ both in their ratio of cases to controls and in how common a taxon is at baseline, subtracting after pooling allows study composition to masquerade as a health effect. This is Simpson's paradox (*106*): a taxon equally prevalent in cases and controls within every single study can still show a large pooled difference, and even one of the opposite sign, if the studies in which it is common happen to have contributed most of the controls. In this dataset the pooled and within-study differences disagree in sign for 12.7% of taxon-by-condition comparisons.

**Table S10. Health Cohort Summary (10% threshold)**. Dep = depletion fraction, Enr = enrichment fraction. Net depletion = Dep − Enr. Med. Log Depth Δ = median log sequencing depth difference. Comparisons. Obesity: Industrial non-obese vs Industrial obese (all health statuses). Glucose Intolerance: Industrial controls vs Industrial T2D / IGT. IBD: Industrial controls vs Industrial IBD / Crohn's / UC. CRC / Adenoma: Industrial controls vs Industrial CRC / Adenoma.

| | | | | | VANISH | | BloSSUM | | VANISH+ | | BloSSUM+ | |
|---|---|---|---|---|---|---|---|---|---|---|---|---|
| Condition | Studies | Controls | Cases | Mean ESS Frac | Dep | Enr | Dep | Enr | Dep | Enr | Dep | Enr |
| Obesity | 29 | 6,603 | 1,219 | 0.958 | 0.024 | 0.008 | 0.222 | 0.048 | 0.11 | 0.07 | 0.22 | 0.12 |
| Glucose Intolerance | 6 | 856 | 440 | 0.942 | 0.024 | 0.016 | 0.111 | 0 | 0.02 | 0.05 | 0.05 | 0.08 |
| IBD | 6 | 726 | 253 | 0.842 | 0.073 | 0.016 | 0.524 | 0.032 | 0.25 | 0.02 | 0.43 | 0.1 |
| CRC / Adenoma | 8 | 610 | 801 | 0.926 | 0.040 | 0.016 | 0.079 | 0.063 | 0.06 | 0.08 | 0.11 | 0.11 |

#### 6C. Primary taxon-level model: GMW-CMH

Naively pooling industrial case and control samples across studies would confound health associations with study membership and sequencing depth. Gut microbiome studies differ in population composition, sequencing depth, extraction methods, and other batch-linked features, so a taxon that is common in one study and rare in another can appear health-associated even when there is no consistent within-study relationship. To avoid this pooling fallacy – Simpson's paradox in its classic form, in which an association present within every stratum reverses or vanishes on aggregation (*106*) – we estimate taxon-level health associations within studies rather than across a single pooled dataset.

For each disease condition and taxon, we therefore use a study-stratified Cochran–Mantel–Haenszel (CMH) framework. Each eligible study contributes a 2×2 table of taxon detection versus absence by control versus case status, and strata are combined to estimate a common within-study association. Strata are eligible only when both groups contain at least five samples. Standard within-study CMH provides a study-adjusted log-odds ratio and p-value, while depth-matched CMH, using 1:1 nearest-neighbor matching on $\log_{10}$(total mapped bases) within each study, is retained as a robustness check rather than the primary analysis, because matching discards samples that lack close depth counterparts.

Our primary taxon-level model is the geometric-mean-target weighted Cochran–Mantel–Haenszel test (GMW-CMH), which extends CMH by equalizing case and control sequencing-depth distributions within each study stratum before the stratified association is computed. Briefly, within each eligible stratum we estimate the depth density separately in controls and cases, reweight both groups toward their shared geometric-mean-target distribution, mean-normalize the weights within group, and clip extreme weights to limit leverage. These weights are then used in place of raw counts when building the per-stratum present/absent tables. This preserves the study-stratified logic of CMH while correcting the within-study depth imbalance that would otherwise bias taxon detection. Full equations for the weighted cell counts and CMH statistic are given in **Supplementary Text 6G.1–6G.2**.

For each taxon, the weighted stratified tables yield the Mantel–Haenszel common risk difference with its Sato standard error, and the Mantel–Haenszel common log-odds ratio, both evaluated at the shared within-study depth target, together with the weighted prevalence in controls and cases, their difference $\mathrm{prev}_{\mathrm{diff}} = \mathrm{prev}_{\mathrm{ctrl}} - \mathrm{prev}_{\mathrm{case}}$, and the two-sided p-value of the CMH test. A Wald statistic and p-value for the risk difference are stored for completeness but carry no inference: the counts in **Supplementary Text 6D** use the CMH p-value, and the list-level inference in **Supplementary Text 6** used in **Fig. 5E** comes from the bootstrap described there.

For each taxon, GMW-CMH yields a common log-odds ratio of detection in controls versus cases, evaluated at the shared within-study depth target, together with the weighted prevalence in controls and cases, their difference $\mathrm{prev}_{\mathrm{diff}} = \mathrm{prev}_{\mathrm{ctrl}} - \mathrm{prev}_{\mathrm{case}}$, and a two-sided p-value. Positive log-odds ratios or positive $\mathrm{prev}_{\mathrm{diff}}$ values denote taxa that are more prevalent in controls and therefore depleted in disease; negative values denote taxa enriched in disease. We also report effective-sample-size diagnostics to quantify the cost of depth equalization within strata. These unthresholded per-taxon outputs are the inputs for the list-level summaries in **Supplementary Text 6D** and the prevalence-matched background analyses in **Supplementary Text 6E**, while standard CMH (*14*, *15*), depth-matched CMH, and Mundlak within–between GLMM (*107*) are treated as robustness analyses in **Supplementary Text 6F**.

**6D. Richness and list-level summary statistics**

Disease-associated shifts in taxon prevalence could, in principle, arise from a general change in microbial richness rather than from list-specific biology. We therefore first summarize richness differences within the same depth-corrected framework used for the taxon-level analyses. For each condition comparison, per-sample richness is reweighted to the shared geometric-mean depth target of the two groups, and weighted mean richness is compared between controls and cases. Confidence intervals are obtained by bootstrap resampling of weight–outcome pairs, with the corresponding disease-specific richness contrasts supporting the richness results shown in **Fig. 5A–D**.

To assess whether any richness difference is concentrated within the focal taxon lists rather than spread uniformly across the microbiome, we further decompose richness into contributions from focal-list taxa and from all remaining taxa. In the primary presentation, this decomposition is reported for VANISH$^{+}$, BloSSUM$^{+}$, and all other taxa; analogous decompositions for the original VANISH/BloSSUM and VANISH$^{H+}$/BloSSUM$^{H+}$ lists are treated as sensitivity analyses. This allows us to distinguish a broad whole-microbiome richness shift from a disproportionate contribution of the industrialization-associated taxa that motivate **Fig. 5**.

We then summarize the taxon-level GMW-CMH outputs at the list level. For each condition × list combination, we report the fraction of taxa significantly depleted in disease, the fraction significantly enriched, and Wilson score 95% confidence intervals for each direction (*108*). Because some individually significant taxa are expected by chance under a $p < 0.05$ threshold, we test whether the observed number of depleted or enriched taxa exceeds the 5% null expectation using a one-sided binomial excess test. A taxon detected in every control and no case, or the reverse, yields a valid table with a small p-value but an odds ratio of zero or infinity; these are counted in the direction given by the sign of the Mantel–Haenszel numerator rather than discarded, since discarding them would remove the most extreme taxa from every count. Where relevant, we also compare the directional profiles of VANISH-type and BloSSUM-type lists using a 2×2 directional contrast test. These list-level summaries provide the compact condition-by-list overview reported in **Table S11** and ensure that the directional signal is not reducible to a generic loss of richness alone.

**Table S11. Geometric-mean-target weighted CMH — depleted and enriched taxa counts per list family and disease condition.** These are ***raw directional counts***, not the prevalence-matched test. Each taxon is counted against zero, not against the shift expected at its own baseline prevalence; the prevalence-matched result is **Table S12**, and that is the comparison the main text reports. Column shading follows the list-family palette used throughout the text. Significance stars are bold. Original VANISH ($n$ = 124) and BloSSUM ($n$ = 63) are the lists published by (*7*). VANISH$^{+}$ / BloSSUM$^{+}$ ($n$ = 100 each) report differentially prevalent taxa between industrial and non-industrial samples. VANISH$^{H+}$ / BloSSUM$^{H+}$ ($n$ = 100 each) are built as the VANISH$^{+}$ / BloSSUM$^{+}$ lists but restrict the industrial samples to those without a disease condition. These counts are corrected for sequencing depth and for study, but each taxon is counted against zero rather than against the shift expected at its own baseline prevalence. Because that expectation falls steeply at high prevalence, a list whose members are common in controls accumulates depleted counts even where it is less depleted than comparable taxa: BloSSUM$^{H+}$ in IBD is 55 of 100 depleted here, and +1.39 SD above its prevalence-matched background in **Table S12**. The two are consistent, and **Table S12** is the comparison the text reports.

| Condition | VANISH | BloSSUM | VANISH$^{+}$ | BloSSUM$^{+}$ | VANISH$^{H+}$ | BLOSSUM$^{H+}$ |
|---|---|---|---|---|---|---|
| Obesity | 3↓ 1↑ /124 | 14↓*** 3↑ /63 | 11↓* 7↑* /100 | 22↓*** 12↑** /100 | 10↓* 7↑ /100 | 27↓*** 11↑* /100 |
| Glucose Intolerance | 3↓ 2↑ /124 | 7↓* 0↑ /63 | 2↓ 5↑ /100 | 5↓ 8↑ /100 | 2↓ 6↑ /100 | 6↓ 6↑ /100 |
| IBD | 9↓ 2↑ /124 | 33↓*** 2↑ /63 | 25↓*** 2↑ /100 | 43↓*** 10↑* /100 | 23↓*** 2↑ /100 | 55↓*** 4↑ /100 |
| CRC / Adenoma | 5↓ 2↑ /124 | 5↓ 4↑ /63 | 6↓ 8↑ /100 | 11↓* 11↑* /100 | 5↓ 9↑ /100 | 16↓*** 7↑ /100 |

Effect direction: ↓ = depleted in cases, ↑ = enriched in cases. Format: dep↓ enr↑ / total. Significance: one-sided binomial excess test against a 5% null, computed separately for depletion and enrichment. . p < 0.1, * p < 0.05, ** p < 0.01, *** p < 0.001.

#### 6E. Prevalence-matched background and the list-level statistic

List-level counts of significant taxa are informative but incomplete. They discard subthreshold directional information and, by themselves, do not address the fact that baseline prevalence constrains how much a taxon can move between controls and cases. A taxon present in 95% of controls has very little room to increase further, whereas a taxon present in 10% of controls can show large apparent changes even under substantial sampling noise. We therefore reference each focal list to a fitted expectation for taxa of the same baseline prevalence, rather than to the raw background pool.

For each disease condition and focal list, we first compute a study-aware control-prevalence coordinate for every taxon. Control-prevalence is calculated separately within each eligible study stratum and then collapsed to a single scalar prevalence by weighting each stratum according to its control sample count. This yields a matching coordinate that reflects only the study contexts in which both cases and controls are represented. On that coordinate we then fit, to the background taxa alone, the expected case-control shift as a function of baseline prevalence: a local-linear regression with a Gaussian kernel of fixed bandwidth $h = 0.12\log_{10}$ units, evaluated on a 400-point grid (*109*). This is a fixed-bandwidth kernel regression rather than LOWESS (*110*): the bandwidth is the same at every prevalence instead of spanning a fixed fraction of nearest neighbours, and no robustness iterations are performed. The local spread is obtained from the local mean absolute residual about that fit, scaled by $\sqrt{\pi/2}$. Each list taxon is then scored by its distance from the fitted curve in units of that local spread, $z = (\text{observed shift} - \text{expected shift})/\text{local spread}$. A background taxon enters the fit in proportion to how close its own control prevalence is to the taxon being scored, and according to nothing else: its standard error, its arm sizes and its number of contributing strata play no part in that weight.

Two constraints govern the fit. The local spread is floored at the taxon's own Sato standard error, so that a taxon's distance from the curve is never expressed in units smaller than its own sampling noise. A taxon detected in no sample of either arm is also excluded from the list mean. Its case-control shift is zero because nothing was observed rather than because nothing moved, and every such taxon falls on the prevalence floor, where all of them would receive the same score whatever their biology. In practice this is the exclusion that matters: across the four conditions it removes 11, 37, 44 and 38 of the 124 original VANISH taxa, which are too rare in industrial populations for the comparison to reach, against at most two taxa in any other list. The 15-neighbour support rule above excludes no taxon in any list or condition. Because the six focal lists overlap heavily (98 of the 100 $\text{VANISH}^{H+}$ taxa and 83 of the 100 $\text{BloSSUM}^{H+}$ taxa also belong to at least one other list) fitting the curve on all non-list taxa would fit it partly on near-duplicates of the list being scored, pulling the expectation toward that list and shrinking $z$. The overlap is not symmetric between the two families, so the attenuation differed between them. The background for every list is therefore the set of taxa belonging to none of the six, which makes the comparison identical in construction for each. Taxa at or above 85% control prevalence are retained but their expected shift is a one-sided extrapolation, since above that prevalence the support window reaches only taxa less prevalent than they are. **Fig. S25** shows this construction directly: every background taxon's shift against its control prevalence, the fitted curve with its ±1 SD band, and the list taxa scored against it. Main-text **Fig. 5E** is the same figure for the $\text{VANISH}^{H+}$ and $\text{BloSSUM}^{H+}$ lists.

Using this fitted expectation, we compare the per-taxon health effects in the focal list to what equally prevalent background taxa would show. Our primary list-level statistic is the mean of $z$ over supported list taxa, with a 95% interval from a bootstrap that resamples the background taxa, refits the curve on each resample, and rescores the resampled list taxa against it. Refitting is the essential step: it propagates the uncertainty in the background expectation itself. The energy distance (*111*) between the observed list and an importance-weighted reference background is retained as a descriptive shape statistic. Energy distance is zero only when the two distributions are identical and increases with any distributional difference, including a shift in center, a change in spread, or a difference in tails. In the present application, the distribution being compared is the distribution of study-stratified case–control shifts, so the energy distance analysis asks whether focal-list taxa show a collectively unusual pattern of depletion or enrichment relative to equally prevalent taxa outside the list.

One further statistic accompanies it. The energy distance between the list and an importance-weighted reference background is retained as a descriptive shape statistic, because it responds to differences in spread and in the tails that a mean does not; it is reported beside the mean $z$ rather than as an independent test of it. Evidence across the two detection specifications is combined by the harmonic mean p-value, as set out in **Supplementary Text 6G.8**. **Table S12** gives both statistics for every reported condition-by-list cell.

Robustness across list families is shown in **Fig. S25**, and taxon-level driver analyses are shown in **Figs. S27–S30**.

##### 6F. Robustness, threshold meta-analysis, and driver taxa

To ensure that the health-association signal is not an artifact of a single detection rule, we repeat the analyses under two detection specifications: the 10% genome-relative breadth threshold and the alternative flat 300 kb mapped-bases threshold. Because these two specifications are applied to the same taxa in the same samples, the resulting p-values are positively dependent rather than independent. For the primary mean-z analysis, we therefore combine evidence across thresholds using the harmonic mean p-value (HMP) (*112*) applied to the bootstrap p-values, and we report the mean of the mean-z values across specifications as a complementary effect-size summary. This provides a conservative measure of threshold robustness and ensures that the primary signal is not driven by only one detection definition.

Our primary presentation centers on the VANISH$^{+}$ and BloSSUM$^{+}$ lists, which are defined on the industrial versus non-industrial contrast used in the main text. We then repeat the same analyses for the original VANISH/BloSSUM lists and, where informative, the VANISH$^{H+}$ and BloSSUM$^{H+}$ lists to assess whether the sign and relative strength of the signal are stable across list definitions. At the model level, we compare the primary GMW-CMH results with standard within-study CMH (*14*, *15*), depth-matched CMH, and Mundlak within–between GLMM fits (*107*). In the GLMM framework, a Hausman-type (*113*) comparison between within-study and between-study coefficients is used to flag conditions in which between-study confounding may remain important. These analyses are intended as robustness checks rather than as co-primary inferential engines.

Finally, to identify the taxa most responsible for a list-level signal, we score each list taxon by its own distance from the background curve, $z$, for each case/control comparison (condition) and summarise it across conditions. A taxon is scored in a condition when the background curve can reach it and when it is detected in at least three of that condition's study cohorts, and it enters the cross-condition summary when at least three of the four conditions qualify. Significance comes from a cluster bootstrap over the 34 contributing study cohorts: each replicate resamples whole cohorts with replacement, recombines the Mantel–Haenszel shift and control prevalence per condition from whichever cohorts were drawn, and re-reads $z$ off the fixed background curve. Resampling cohorts, rather than treating the four condition values as independent observations, is necessary rather than cosmetic: a one-sample t-test on four values has three degrees of freedom and returns no significant taxa for BloSSUM$^{+}$ or BloSSUM$^{H+}$ even though 87 and 78 of their 100 scored taxa lie on the same side of the background. Per-taxon significance is reported after Benjamini–Hochberg correction (*114*) within each list, and the figures draw every taxon reaching $q < 0.20$, subject to a floor of ten rows so that a list with little signal does not collapse to a stub; the per-condition cells are annotated with the uncorrected within-condition bootstrap p, which describes where a taxon's signal sits rather than making a second corrected claim. Taxa detected in no sample of either arm are excluded, since their case–control shift is zero through absence of observation rather than absence of change.

**Table S12.** List-level tests compared across methods and detection specifications. One row per disease condition and taxon list, grouped by list. Mean $z$ is the mean distance of the list from the fitted prevalence background, in units of the local spread, given under each of the two detection specifications: a 10% genome-relative breadth threshold and a flat 300,000 mapped-bases threshold. Mean $z$ p is the two-sided bootstrap p-value under each specification. The two $p_{\mathrm{HMP}}$ columns give the harmonic mean p-value across the two specifications for the mean $z$ against the background curve, which is the test, and for the energy distance, which is descriptive. Agree counts how many of the two reach $p < 0.05$. Both are two-sided and both are referenced to the same prevalence-matched background, so their agreement is informative; they differ in what they are sensitive to, the mean $z$ to a shift in location and the energy distance also to spread and tails. Rows are the condition-by-list contrasts reported in **Fig. 5E** and **Fig. S25** (those resting on at least three studies with at least five cases and five controls).

| Condition | List | Two-sided, referenced to the background<br>n | Mean z<br>10% / 300 kb | Mean z p<br>10% / 300 kb | Mean z<br>p HMP | Energy<br>p HMP | Agree |
|---|---|---|---|---|---|---|---|
| Obesity | VANISH | 113 | +0.35 / +0.37 | ≤0.0005 / ≤0.0005 | **≤0.0005** *** | ≤0.0005 | 2/2 |
| Glucose Intolerance | VANISH | 87 | +0.04 / +0.03 | 0.636 / 0.752 | 0.689 | 0.384 | 0/2 |
| IBD | VANISH | 80 | -0.16 / -0.16 | 0.060 / 0.032 | **0.042** * | 0.344 | 1/2 |
| CRC / Adenoma | VANISH | 86 | +0.10 / +0.16 | 0.342 / 0.125 | 0.183 | 0.258 | 0/2 |
| Obesity | BloSSUM | 63 | +0.06 / +0.03 | 0.696 / 0.826 | 0.755 | 0.328 | 0/2 |
| Glucose Intolerance | BloSSUM | 63 | +0.20 / +0.10 | 0.145 / 0.451 | 0.219 | 0.307 | 0/2 |
| IBD | BloSSUM | 63 | +1.42 / +1.35 | ≤0.0005 / ≤0.0005 | **≤0.0005** *** | ≤0.0005 | 2/2 |
| CRC / Adenoma | BloSSUM | 63 | +0.61 / +0.62 | ≤0.0005 / ≤0.0005 | **≤0.0005** *** | 0.0007 | 2/2 |
| Obesity | VANISH$^{+}$ | 100 | +0.67 / +0.70 | ≤0.0005 / ≤0.0005 | **≤0.0005** *** | ≤0.0005 | 2/2 |
| Glucose Intolerance | VANISH$^{+}$ | 99 | +0.24 / +0.19 | 0.014 / 0.043 | **0.021** * | 0.336 | 1/2 |
| IBD | VANISH$^{+}$ | 99 | -0.22 / -0.23 | 0.013 / 0.006 | **0.008** ** | 0.010 | 2/2 |
| CRC / Adenoma | VANISH$^{+}$ | 100 | +0.11 / +0.15 | 0.244 / 0.114 | 0.155 | 0.368 | 0/2 |
| Obesity | BloSSUM$^{+}$ | 100 | +0.32 / +0.35 | 0.065 / 0.057 | 0.061 | 0.002 | 1/2 |
| Glucose Intolerance | BloSSUM$^{+}$ | 100 | +0.58 / +0.47 | ≤0.0005 / 0.001 | **0.0007** *** | ≤0.0005 | 2/2 |
| IBD | BloSSUM$^{+}$ | 100 | +1.83 / +1.68 | ≤0.0005 / ≤0.0005 | **≤0.0005** *** | ≤0.0005 | 2/2 |
| CRC / Adenoma | BloSSUM$^{+}$ | 100 | +0.91 / +0.81 | ≤0.0005 / ≤0.0005 | **≤0.0005** *** | ≤0.0005 | 2/2 |
| Obesity | VANISH$^{H+}$ | 100 | +0.70 / +0.73 | ≤0.0005 / ≤0.0005 | **≤0.0005** *** | ≤0.0005 | 2/2 |
| Glucose Intolerance | VANISH$^{H+}$ | 99 | +0.22 / +0.18 | 0.021 / 0.079 | **0.033** * | 0.370 | 1/2 |
| IBD | VANISH$^{H+}$ | 98 | -0.19 / -0.20 | 0.041 / 0.030 | **0.035** * | 0.023 | 2/2 |
| CRC / Adenoma | VANISH$^{H+}$ | 100 | +0.14 / +0.19 | 0.150 / 0.063 | 0.089 | 0.087 | 0/2 |
| Obesity | BloSSUM$^{H+}$ | 100 | +0.13 / +0.12 | 0.398 / 0.441 | 0.418 | 0.052 | 0/2 |
| Glucose Intolerance | BloSSUM$^{H+}$ | 100 | +0.44 / +0.33 | ≤0.0005 / 0.016 | **0.0010** *** | ≤0.0005 | 2/2 |
| IBD | BloSSUM$^{H+}$ | 100 | +1.39 / +1.31 | ≤0.0005 / ≤0.0005 | **≤0.0005** *** | ≤0.0005 | 2/2 |
| CRC / Adenoma | BloSSUM$^{H+}$ | 100 | +0.61 / +0.51 | ≤0.0005 / 0.001 | **0.0007** *** | ≤0.0005 | 2/2 |

† marks the one cell where the two detection specifications disagree in the sign of the mean $z$, at |mean $z$| = 0.009 and non-significant under either specification. The bootstrap draws 2,000 replicates, so a p-value cannot resolve below 1/2,000 and is reported as ≤0.0005. *** $p < 0.001$, ** $p < 0.01$, * $p < 0.05$, applied to the combined mean $z$.

**6G. Technical appendix for Supplementary Text 6**

Formal equations and low-level implementation details are collected here for completeness. These details do not change the estimands defined in the main body of **Supplementary Text 6**, but they are provided so that the study-stratified CMH, depth-matched CMH, GMW-CMH, prevalence-matched background construction, and downstream robustness analyses can be reproduced exactly.

**6G.1. Standard CMH and Mantel–Haenszel common log-odds ratio**

For each taxon and each eligible study stratum $k$, we construct a 2×2 table of taxon detection versus absence by control versus case status:

$$\begin{pmatrix} & \text{Present} & \text{Absent} \\ \text{Healthy} & a_k & b_k \\ \text{Unhealthy} & c_k & d_k \end{pmatrix}$$

where $a_k$, $b_k$, $c_k$, and $d_k$ denote the control-present, control-absent, case-present, and case-absent cells, respectively, and $n_k = a_k + b_k + c_k + d_k$ is the total number of samples in the stratum. Only strata with at least five controls and five cases are eligible.

Under the null hypothesis that taxon detection is unrelated to case-control status within a stratum, the expected value of the control-present cell is:

$$E[a_k] = \frac{(a_k + b_k)(a_k + c_k\ )}{n}$$

The CMH statistic aggregates the deviations between observed and expected counts across all studies $k$, weighted by how much information each study contributes:

$$\chi^2 = \frac{(|\sum_k (a_k - E[a_k])| - 0.5)^2}{\sum_k V[a_k]}$$

with Yates' continuity correction applied as in the standard one-degree-of-freedom CMH formulation (*115*). The variance $V[a_k]$ measures how much $a_k$ would fluctuate by chance under the null, given the row and column totals of that study's table:

$$V[a_k] = \frac{(a_k + b_k)(c_k + d_k)(a_k + c_k)(b_k + d_k)}{n^2(n-1)}$$

The resulting statistic is compared to a chi-square distribution with one degree of freedom to obtain a p-value.

To summarize direction and magnitude across strata, we also compute the Mantel–Haenszel common log-odds ratio:

$$\log(OR_{MH}) = \log\left(\frac{\sum_k a_k d_k / n_k}{\sum_k b_k c_k / n_k}\right)$$

A positive log-odds ratio indicates that the taxon is more prevalent in controls and therefore depleted in cases; a negative value indicates enrichment in cases.

**6G.2. GMW-CMH weighting and weighted 2×2 tables**

The primary taxon-level method is the geometric-mean-target weighted Cochran–Mantel–Haenszel test (GMW-CMH), which extends the standard study-stratified CMH framework by equalizing control and case depth distributions within each eligible study stratum before the 2×2 tables are aggregated.

For each eligible stratum, let $f_{\text{ctrl}}(m)$ and $f_{\text{case}}(m)$ denote kernel density estimates of $\log_{10}$(total mapped bases $m$) in controls and cases, respectively. Both densities are estimated using Silverman's bandwidth rule (*102*). The shared target distribution is the geometric mean:

$$f^*(m) = \sqrt{f_{\text{ctrl}}(m) f_{\text{case}}(m)}$$

which is symmetric in the two groups and therefore does not privilege either group's empirical depth distribution.

If a stratum contains fewer than five samples in either group or fewer than three distinct depth values in either group, all weights in that stratum default to 1.0, making that stratum equivalent to an unweighted CMH contribution.

For each control sample $i$ in stratum $k$, the weight is:

$$w_i = \sqrt{\left(f_{\text{case}}(m_i)/f_{\text{ctrl}}(m_i)\right)}$$

and for each case sample $j$:

$$w_j = \sqrt{\left(f_{\text{ctrl}}(m_j)/f_{\text{case}}(m_j)\right)}$$

Weights are mean-normalized within group and stratum so that the average weight is 1.0 in each group, then clipped at a maximum of 10 to prevent extreme leverage. Samples with missing depth values receive weight 1.0.

Weighted 2×2 tables are then constructed by replacing integer counts with weighted effective counts. Using $\mathbf{1}[\text{present}]$ and $\mathbf{1}[\text{absent}]$ as indicator functions for taxon detection:

$$a_k = \sum_i w_i \cdot \mathbf{1}[\text{present}_i], \quad b_k = \sum_i w_i \cdot \mathbf{1}[\text{absent}_i]$$

$$c_k = \sum_j w_j \cdot \mathbf{1}[\text{present}_j], \quad d_k = \sum_j w_j \cdot \mathbf{1}[\text{absent}_j]$$

These weighted cells are passed unchanged into the standard CMH aggregator. Because the weights are mean-normalized, the total weight in each group remains approximately equal to the original sample count, while the within-group composition of shallow and deep samples is adjusted so that controls and cases contribute as though they had been drawn from the same within-study depth distribution.

**6G.3. The Mantel–Haenszel common risk difference**

On the prevalence scale, let $n_{1k}$ and $n_{2k}$ be the weighted control and case arm sizes in stratum $k$, $A_k$ and $C_k$ the weighted detection counts, and $N_k = n_{1k} + n_{2k}$. With $W_k = n_{1k}n_{2k}/N_k$, the risk difference is defined as

$$\mathrm{rd}_{MH} = \Sigma_k W_k (A_k/n_{1k} - C_k/n_{2k})/\Sigma_k W_k.$$

Its variance follows Sato (*105*):

$$\mathrm{Var}(\mathrm{rd}_{MH}) = (\mathrm{rd}_{MH}\Sigma_k P_k + \Sigma_k Q_k)/(\Sigma_k W_k)^2,$$

where

$$P_k = ({n_{1k}}^2 C_k - {n_{2k}}^2 A_k + n_{1k} n_{2k}(n_{2k} - n_{1k})/2)/{N_k}^2,$$

$$Q_k = \big(A_k(n_{2k} - C_k) + C_k(n_{1k} - A_k)\big)/(2N_k).$$

This estimator is consistent in the sparse-data limit as well as the large-strata limit (*104*, *116*), which is the relevant regime here because many strata contribute only a few samples per arm. In the single-stratum limit it reduces exactly to $p_1(1 - p_1)/n_1 + p_2(1 - p_2)/n_2$.

##### 6G.4. Per-taxon outputs and ESS reporting

For every taxon × condition × method combination, we store the weighted prevalence in controls ($\mathrm{prev}_{\mathrm{ctrl}}$), the weighted prevalence in cases ($\mathrm{prev}_{\mathrm{case}}$), their difference

$$\mathrm{prev}_{\mathrm{diff}} = \mathrm{prev}_{\mathrm{ctrl}} - \mathrm{prev}_{\mathrm{case}}$$

the common log-odds ratio, the two-sided p-value, and the number of contributing strata. Positive $\mathrm{prev}_{\mathrm{diff}}$ values indicate taxa that are more prevalent in controls and therefore depleted in cases.

Weight dispersion is summarized using the Kish effective sample size:

$$ESS = (\Sigma w_i)^2/\Sigma w_i^2$$

Within each stratum and group, ESS is computed from the GMW weights. We then report the group-level ESS and the condition-level ESS fraction ($\mathrm{ESS}_{\mathrm{frac}}$), defined as the average of ESS per nominal sample size across the two groups and across contributing strata. $\mathrm{ESS}_{\mathrm{frac}}$ therefore quantifies the information cost of within-stratum depth equalization: values near 1 indicate minimal weighting cost, whereas lower values indicate stronger depth imbalance and greater reliance on a subset of weighted samples.

These unthresholded per-taxon outputs are used downstream for the prevalence-matched background comparisons, directional summaries, and taxon-driver analyses.

##### 6G.5. Background construction

For each disease condition and focal taxon list, we reference the list to taxa of the same baseline prevalence. Step 1 fixes the matching coordinate. Step 2 fits the expectation used by the primary statistic. Importance weighting survives only in the secondary tests of 6G.6, where a matched reference set rather than a fitted curve is required.

**Step 1: Stratum-aware control prevalence.** For each taxon, baseline prevalence is computed separately within each eligible study stratum in the control group only. Strata that do not meet the minimum case/control size requirement are excluded from this step. The stratum-specific control prevalences are then collapsed to a single scalar prevalence coordinate by weighting each stratum in proportion to its control sample count:

$$p_i = \sum_{s \in S} w_s \, p_{i,s}, \qquad w_s = \frac{n_s^{ctrl}}{\sum_{s' \in S} n_{s'}^{ctrl}}$$

This scalar control prevalence is the matching coordinate used throughout the prevalence-matching procedure.

**Step 2: The background curve.** Let $x_i = log_{10} p_i$ be the coordinate from Step 1 and $y_i$ the case-control shift of taxon $i$ in percentage points. Fitted on the background taxa alone, the expected shift is the local-linear Gaussian-kernel estimate with fixed bandwidth $h = 0.12$, evaluated on a 400-point grid over $[10^{-4}, 1]$; the kernel weight of background taxon $j$ at a point $x$ is $exp\left(-\left(x_j - x\right)^2 / 2h^2\right)$ and depends on nothing else about that taxon. The local scale is $\sqrt{\pi/2}$ times the local-linear fit of the absolute residuals, floored at the focal taxon's Sato standard error. Each list taxon is scored $z_i = \left(y_i - \mu(x_i)\right)/\sigma(x_i)$, and is supported, and so enters the list mean, only if at least 15 background taxa lie within 0.25 $\log_{10}$ of $x_i$. Background taxa are those belonging to none of the six focal lists.

##### 6G.6. List-level test implementation

Four list-level comparisons are performed between the focal list and its prevalence-matched background.

**Mean $z$ against the background curve (primary list-level test)**. The statistic is the mean of $z$ over supported list taxa. Its interval is obtained by resampling the background taxa with replacement and refitting the curve on each resample, resampling the supported list taxa and rescoring them against that refitted curve, and recording the mean. We take 2,000 replicates under a fixed seed and report the 2.5th and 97.5th percentiles, with a two-sided p-value as twice the smaller tail proportion. A percentile bootstrap (*13*) cannot resolve a tail finer than one replicate, so the p-value is floored at 1/2,000 and no significance band finer than that floor is used. This per-specification p-value is the quantity combined across detection rules in **Supplementary Text 6G.8**.

**IW-energy distance (descriptive shape statistic).** The energy distance between the observed focal-list values and a fixed importance-weighted reference background is retained as a descriptive summary, because it is sensitive to differences in spread and tails that a mean is not. The energy distance between two distributions $F$ and $G$ is:

$$E(F, G) = 2\mathrm{E}|X - Y| - \mathrm{E}|X - X'| - \mathrm{E}|Y - Y'|$$

where $X, X'$ are independent draws from $F$ and $Y, Y'$ from $G$. It equals zero if and only if $F = G$, and is positive otherwise. Unlike a mean-shift test, the energy distance is a summary statistic that is sensitive to any distributional difference: a shift in the center, a change in spread, a difference in skewness, or a difference in tails.

Implementation proceeds as follows. First, a matched reference distribution is constructed by drawing $20 \times n_{\text{list}}$ background observations with probability proportional to their importance weights. This reference represents the expected $\text{prev}_{\text{diff}}$ distribution for a set of taxa with the same control-prevalence profile as the focal list. The observed statistic $E_{\text{obs}}$ is the energy distance between the actual focal-list values and this fixed matched reference. The null distribution is then generated by drawing $n_{\text{list}}$ background taxa, again with importance-weighted probability, and computing the energy distance from the same fixed reference. This null draw is repeated 5,000 times. The empirical p-value is the fraction of null energy distances that are at least as large as the observed statistic, floored at $1/(n+1)$.

Because both the observed list and the null draws are compared to the same prevalence-matched reference, this is a contamination-free null: it asks specifically whether list membership predicts a distribution of health associations that departs from what equally prevalent non-list taxa would show.

##### 6G.7. Mundlak within–between GLMM and Hausman diagnostic

As an additional robustness analysis, we fit a generalized linear mixed model for each taxon using the Mundlak device (*107*) to decompose the group effect into within-study and between-study components. The within-study coefficient $\beta_w$estimates the same basic estimand as the CMH family of methods, namely the health association after removing study-level variation. The between-study coefficient $\beta_b$ captures the portion of the apparent signal that could be attributable to study-level confounding.

A Hausman-type comparison is then used to test whether $\beta_w$ and $\beta_b$ differ materially for a given taxon. Conditions in which more than 20% of taxa show a significant within-vs-between discrepancy may be flagged as potentially affected by residual between-study confounding and interpreted with additional caution. This GLMM/Hausman procedure is used only as a robustness diagnostic and not as the primary inferential engine.

##### 6G.8. Threshold combination across detection specifications

All primary analyses are repeated under two taxon-detection specifications: a genome-relative breadth threshold of 10% and an alternative flat mapped-bases threshold of 300,000 mapped bases. Because both specifications are applied to the same taxa in the same samples, the resulting $p$-values are positively dependent rather than independent.

For the primary mean-z analysis, evidence across thresholds is combined using the harmonic mean p-value (HMP):

$$p_{HMP} = \frac{k}{\sum_{i=1}^{k} 1/p_i}$$

where $k$ is the number of detection specifications combined (here $k = 2$: the 10% genome-relative breadth threshold and the 300,000 mapped-bases flat threshold), $p_i$ is the bootstrap p-value of the mean z under specification $i$ for a given condition and focal list, and $p_{HMP}$ is the resulting threshold-robust combined p-value (*112*). The two components are weighted equally, so the general weighted form of the harmonic mean p-value reduces to the expression above. The HMP is used because it remains valid under arbitrary dependence between the component tests (*112*), which is what two detection rules applied to the same taxa in the same samples require. The combined value is written to health_background_curve_meta.csv alongside each specification's own mean z and interval, and it is the value quoted in the main text. Because the bootstrap floors each $p_i$ at 1/2,000, the HMP inherits that floor. The same combination is applied to the energy-distance permutation p-values as a descriptive check, and the mean of the mean-z values across specifications is reported as a companion effect-size summary.

Agreement between the two specifications is tabulated separately rather than folded into the combined p-value (**Table S12**), since the HMP measures the strength of the combined evidence and not the concordance of its parts: across the thirty reported condition-by-list cells the two rules agree in the sign of the mean $z$ for twenty-nine, and the single disagreement has $|\text{mean } z| = 0.009$ and is non-significant under either rule. **Fig. 5E** and **Fig. S25** print the mean $z$, its interval and its bootstrap p under the primary rule, breadth ≥ 10%. In this run the significance at the 5% level for every condition-by-list cell agree when calculated for the breadth ≥ 10% detection rule or from the HMP combining the breadth ≥ 10% and breadth ≥ 300 kb rules.

##### 6G.9 Within-clade contrasts

To test whether the list-level signal survives holding taxonomy fixed, list members are compared with their own relatives (**Fig. S26**). Within each genus or family containing at least five catalogue taxa, the background curve is refitted with that clade excluded (otherwise a large clade is partly its own control) and every clade member is scored against the refitted curve. The clade is summarised by its mean $z$, against a null of 5,000 placebo clades whose members are drawn from non-clade taxa within one bandwidth of each real member's control prevalence, so that a clade cannot score merely by being unusually common or rare. Then, within each clade holding at least three list taxa and three non-list taxa, we take the mean $z$ of the list members minus that of the remaining members, with a p-value from permuting the list labels inside the clade. Taxa at or above 85% control prevalence are excluded here, since the clade-excluded refit requires a neighbourhood on both sides. Contrasts are corrected across clades by the Benjamini–Hochberg procedure (*114*).

#### Supplementary Text 7. Robustness analyses and limitations

The analyses in this study were designed to distinguish genuine biological differences from artifacts introduced by sequencing depth, reference choice, and study structure. Several robustness checks support the main conclusions. At the same time, these results (and in particular the persistence results) should be interpreted as claims about detectability and prevalence within current gut reference catalogues, not as proof of complete functional equivalence across lifestyles or of universal persistence of all gut lineages. This note summarizes the principal robustness analyses and clarifies what the conclusions do and do not imply.

##### 7A. Reference dependence and the bounds of inference

A central limitation of any reference-based metagenomic analysis is that inference depends on the genomes represented in the reference catalogue. This concern is especially relevant here because the two most complete currently available gut references were constructed from different source populations and therefore carry different ascertainment biases. The Carter reference is, by construction, enriched in genomes identified from deeply sequenced non-industrial samples, whereas the Leviatan reference is, by construction, enriched in genomes identified from industrial samples. Because these particular biases apply in opposite directions, the "ground truth" of the composition of a typical meta-microbiome is likely to lie within the bounds set by alignment to these two reference metagenomes. For this reason, we aligned all samples to both references when examining persistence and attenuation across lifestyles.

The main population-level conclusion is robust to this choice. Using both the non-industrial-biased Carter reference and the industrial-biased Leviatan reference, we found that industrial and non-industrial populations converge toward similar total detectable richness when sufficiently many samples are pooled, even though the path to saturation differs across references and lifestyles. This indicates that the broad conclusion is not an artifact of choosing a reference enriched for one lifestyle or the other. At the same time, exact taxon-level detectability remains reference-dependent, and differential missingness is inherently difficult to quantify. Accordingly, our conclusions concern the detectability of taxa represented in current gut reference catalogues; they do not rule out loss of unsampled lineages, strain-level diversity, or functions not captured by these references.

##### 7B. Read-length sensitivity and other sequencing-design effects

Read length is a real technical confound in comparative metagenomic analyses, and is one reason we measured sequencing effort as mapped bases rather than as reads per sample: the same read count represents different amounts of sequence at 100 bp and at 150 bp per read, with many older studies of industrial populations using shorter reads. **Fig. S14** shows the detected richness as a function of sequencing depth for short, medium, and long reads. There is a modest but significant decrease in detected richness with read length, but the result is inconsistent between industrial and non-industrial samples and appears to be driven by a few outlier data bins. We think that it is more likely that this difference is driven by study batch effects, and any read-depth related effect is much smaller than either the sequencing-depth confounding that we have identified and are principally correcting for or the real differences in richness that we have confirmed between industrial and non-industrial populations.

##### 7C. Sample independence: repeated sampling of hosts and technical replicates

The 18,075 samples come from 14,624 hosts. Twenty-three of the 83 studies sampled some participants more than once (longitudinally, before and after an intervention, or across seasons), so 3,451 samples are repeat samples of a host already represented. The largest contributors are Lloyd-Price et al. 2019 (*51*) (1,125 samples from 104 hosts), iMSMS 2022 (*96*) (1,132 from 611) and Mehta et al. 2018 (*59*) (803 from 301). Repeat samples are not independent observations, and how much that matters depends on the analysis.

*Richness comparison* (**Fig. 2**). The GMW bootstrap resamples samples. We therefore repeated the industrial versus non-industrial comparison (**Table S13**, **Fig. S13**) (i) resampling hosts rather than samples, keeping each host's samples together, which leaves the richness ratio at 0.614 and widens its 95% interval from 0.601–0.627 to 0.599–0.629; and (ii) retaining one sample per host (the sample used in the health analyses), which gives 0.643 (0.630–0.658). Industrial richness remains substantially lower after depth correction under both.

*Accumulation and pooled detection* (**Figs. 3–4**). Repeat samples from one host share more taxa than samples from different hosts. They can therefore only slow the accumulation of newly detected taxa, making curves in samples conservative relative to curves in hosts. Repeat sampling does not change VANISH$^+$/BloSSUM$^+$ membership (**Supplementary Text 5**).

*Health associations* (**Fig. 5**). Here repeat samples would inflate a study's weight in the pooled estimate and its cell counts in the per-taxon CMH, so every case–control comparison retains only one sample per host (**Supplementary Text 6B**). This removed 2,382 samples from the pooled Unhealthy comparison, 1,353 from IBD and 989 from obesity; the pooled figure exceeds the sum of the two condition-specific ones because the pooled contrast draws on all health conditions, not only these two.

*Unrecorded repeats.* Host identity comes from archived subject identifiers. 5,042 samples have none and are treated as separate hosts, so some repeat sampling may be unrecorded.

*Technical replicates*. Public metagenomic repositories contain a small number of nominal samples represented by multiple sequencing runs. By default, multiple runs for the same nominal sample were collapsed to a single sample, prioritizing paired-end data where available. This follows the logic used in the metadata curation pipeline, in which older studies sometimes included multiple accessions for the same sample as technical replicates across different sequencing runs. Treating these as a single sample avoids artificially inflating sample counts and prevents run-level duplication from biasing population-level accumulation or prevalence estimates.

The technical replicate checks suggest that this choice is conservative and does not drive the main conclusions. Technical replicates show high richness reproducibility, and within-sample Bray-Curtis dissimilarity between technical replicates is low and largely independent of sequencing depth for the large majority of pairs. A small subset of nominal replicate pairs show elevated dissimilarity, consistent with compositionally distinct samples or metadata misassignment rather than ordinary run-level noise. Because this only affected a small number of samples and was study-dependent, we chose not to include these pairs as separate samples. Replicate handling is unlikely to explain the principal richness, persistence, or health-association results, although imperfect public metadata remain a residual limitation of large-scale meta-analysis.

**Table S13. Industrial-to-non-industrial richness ratio under repeat sampling of hosts.** The ratio is GMW-corrected mean richness (SGBs detected at ≥10% breadth coverage, combined reference) in industrial samples divided by that in non-industrial samples; the first row is the estimate reported in **Fig. 2** and the main text. 95% confidence intervals are 2.5$^{th}$–97.5$^{th}$ percentiles of 1,000 bootstrap replicates drawn within each group, resampling either samples or hosts (all samples of a drawn host kept together). One sample per host uses each host's representative sample, the sample retained in the health analyses, with GMW weights refitted on that cohort.

| Analysis | Ratio | 95% CI | Industrial samples (hosts) | Non-industrial samples (hosts) |
|---|---|---|---|---|
| All samples; bootstrap over samples (as in **Fig. 2**) | 0.614 | 0.601–0.627 | 13,977 (10,847) | 4,098 (3,777) |
| All samples; bootstrap over hosts | 0.614 | 0.599–0.629 | 13,977 (10,847) | 4,098 (3,777) |
| One sample per host | 0.643 | 0.630–0.658 | 10,847 (10,847) | 3,777 (3,777) |

#### 7D. Sensitivity to detection thresholds

The distinction between a rare taxon and a spurious low-breadth signal depends on the detection threshold used. For this reason, the analyses were not based on a single arbitrary cutoff alone. Across the manuscript and supplementary analyses, we examined both a genome-relative breadth threshold and a flat mapped-base threshold, and we also explored a wider range of thresholds in sensitivity analyses. These thresholds embody the same biological intuition: genuine representation should produce breadth and depth patterns consistent with reads accumulating across a substantial fraction of the genome, whereas short contaminant fragments or plasmid-like signals should saturate at much smaller effective represented lengths.

The principal conclusions are robust across these choices. Across a wide range of thresholds, the GMW-corrected richness gap between industrial and non-industrial samples remained substantially negative even though the raw gap was consistently more negative than the corrected one, confirming that lower sequencing depth inflates the apparent richness deficit in industrial samples but does not fully explain it. Likewise, the population-level persistence results were not driven by a single permissive threshold: examination of pooled samples showed that all VANISH$^{+}$ taxa were detectable in industrial populations and all BloSSUM$^{+}$ taxa were detectable in non-industrial populations under both breadth-based specifications tested. Threshold choice therefore influences exact effect sizes and the status of a small number of edge-case taxa more than it influences the direction of the study's main conclusions. Any claims about nominally undetected taxa should accordingly be interpreted alongside the relevant threshold- and reference-specific analyses.

#### 7E. Persistence at low prevalence does not necessarily imply functional importance

The present study shows that many taxa proposed to be absent from industrial populations remain detectable when microbiomes are pooled across sufficiently many industrial samples. However, detectability at the population level does not by itself establish ecological importance or host benefit. Although the health associations reported here are consistent with a possible role for depleted taxa in health and disease, it remains unclear whether taxa present at such low prevalence retain functional significance for the host or instead represent transient colonisers that lack stable ecological niches in the industrial gut environment.

This distinction matters for interpretation. A taxon may persist at low prevalence in a population without being abundant within individuals, stably transmitted, or functionally consequential under industrial ecological conditions. Conversely, even low-prevalence reservoirs may still matter if they preserve ecologically important functions or enable recolonization under favorable conditions. Further work to delineate the functions and physiological effects of these depleted taxa — for example, via spike-in experiments and studies of mock communities of bacterial isolates in which the presence and abundance of these taxa can be manipulated — will be important for discriminating between these alternatives.

##### 7F. Distinguishing prevalence from abundance

A recurring interpretive issue in this study is the distinction between prevalence and abundance. In this work, prevalence refers to the fraction of samples in which a taxon is detected, whereas abundance refers to the within-sample quantity of that taxon conditional on detection. The VANISH/BloSSUM and VANISH$^+$/BloSSUM$^+$ results are fundamentally prevalence-based: these taxa are defined by differential prevalence across lifestyles, and the persistence/extinction analyses are likewise statements about detection and prevalence at the population level. They are therefore not direct claims about within-host abundance.

This distinction helps clarify several points that might otherwise sound stronger than intended. The VANISH taxa are not a measure of bacterial "missingness" in the absolute sense; rather, they are a measure of differential prevalence. For the same reason, when this study describes taxa as more or less common, the intended meaning is more or less prevalent across samples, not necessarily more or less abundant within individual hosts. Abundance enters separately in the compositional analyses and in the prevalence-versus-CLR-abundance contrasts used to refine taxon lists, but all persistence and attenuation claims should be read as claims about population-level detection and prevalence.

##### 7G. Attenuation versus extinction

The central interpretive conclusion of this study is that industrialization is more consistently associated with attenuation than with widespread extinction of represented gut taxa. Individuals in industrial populations do harbor lower gut microbial richness than individuals in non-industrial populations, but this does not mean that industrial populations have suffered collective microbial extinctions. Rather, many microbes are less prevalent than in non-industrial populations. In this study, attenuation refers to a reduction in population-level prevalence or detectability relative to non-industrial populations, whereas extinction would require failure of detection after broad pooling under the stated reference and threshold framework.

Under that operational definition, the results argue against widespread local extinction among currently represented gut SGBs and instead support a model in which many taxa persist at reduced prevalence across industrial populations. This interpretation is consistent with the observation that nearly all previously highlighted VANISH taxa, and all of the empirically derived VANISH$^+$ taxa, remained detectable when industrial samples were pooled broadly enough. The conclusions therefore reframe the effects of industrialization on the gut microbiome from matters of extinction to matters of attenuation. At the same time, they do not rule out the loss of unrepresented lineages, ecologically important reductions below functional thresholds, or loss of microbial traits not resolved at the level of represented SGBs.

**Glossary**

**Sequencing depth:** The total number of DNA reads (100-150 bp fragments) that are sequenced per sample. Shotgun metagenomic samples typically have 0.5-30 million reads per sample, but some studies have employed much deeper sequencing of up to 100 million reads per sample (e.g., Carter et al. 2023 (*7*)).

**Reference metagenome:** A set of high-quality (low contamination, high breadth coverage) microbial genomes binned into distinct Species Genome Bins (SGBs). This article considers biases related to the curation of the reference metagenomes as well as potential difficulties related to distinguishing SGBs from each other and incorrect assignment of bacterial reads to SGBs.

**Breadth coverage:** The fractional coverage (typically expressed in %) of an SGB within a given sample. This article discusses issues with assigning microbial species detection based on a fixed breadth coverage threshold.

**Depth coverage:** The average coverage of an SGB, expressed as the total number of reads mapping to the SGB times the read length divided by the length of the SGB. Depth coverage is not by itself a useful measure of whether an SGB is in a sample, but it can be combined with breadth coverage to effectively correct for differences in sampling depth.

**Richness:** The total number of microbial species detected in an individual or population. The theoretical upper bound of richness is the number of species actually present in the microbiota of that individual or population. Sequencing a microbiota at limited depth does not provide sufficient evidence to find all of these species, so the observed richness is lower than this theoretical upper bound; for instance, rarefaction analyses by Carter et al. 2023 (*7*) (their Figure 3A) find increasing numbers of SGBs even beyond 50Gbp/sample. Modeling is therefore required to estimate the true number of species in a sample.

**Diversity:** There are numerous methods to quantify microbiota diversity (*101*). In the context of this study, we use diversity to refer to α-diversity quantified as richness.

**Prevalence:** Fraction of samples in which a particular microbe is detected.

**Composition:** The identities and prevalences of the detected microbial species in an individual or population. The compositions of any two independent microbiota samples, even those sampled from within the same population, can be very different. Two randomly chosen individuals sampled from the same population share around 20% and at most 40% of their detected microbial species. For this reason, to characterize population-level microbiota composition, it is necessary to broadly sample the microbiota of many independent individuals within this population. Microbial population-level diversity thus increases along with sampling breadth.

**Hunter-gatherer, agro-pastoralist, industrial:** The three lifestyle categories used throughout. These were either seeded from existing lifestyle analyses (e.g. (*7*)), from the SRA, or curated from the titles of studies. Some hunter-gatherer curations (e.g. in (*73*)) were used to subdivide studies by lifestyle. Most studies of industrial populations focused on disease comparisons among urban cohorts and were easy to classify. The intermediate agro-pastoralist category is broad and spanned a large portion of the lifestyle gradient from nomadic pastoralists in Africa and Asia to rural communities in Latin America (**Fig. 2F**, which shows that the 3-tier lifestyle classification is a little fuzzy in borderline populations).

**Case, control, and "unhealthy":** A case is a sample from a host recorded as having the condition under test; a control is a sample from the same study without it. The pooled unhealthy group is any of 26 recorded conditions (**Supplementary Text 6B**); it is a convenience grouping used to examine microbial richness in healthy vs unhealthy samples, not a clinical category, and a reader should not read it as a disease.

**Glucose intolerance:** Type 2 diabetes together with impaired glucose tolerance. The broader term is used because the eligible cohorts contain both, and separating them would drop the comparison below the three-study minimum.

**Species-level genome bin (SGB):** A cluster of assembled genomes sharing at least 95% average nucleotide identity, used here as the working unit of "a bacterial species". Detection is always detection of an SGB in a reference catalogue, never of an organism directly, so every statement about presence is relative to what the catalogue contains.

**Multimapping and MAPQ:** A 100–150 bp read is often short enough to align equally well to several closely related genomes; such a read is multimapping, and the aligner reports its ambiguity as a low MAPQ score. We discard reads below MAPQ 30, which removes a substantial fraction of aligned reads but prevents one species' reads being credited to its relatives, which is the failure mode that would most inflate apparent richness in a large catalogue.

**Pooling, and population-level detection:** A taxon is detected at the population level if it appears in at least one sample once a set of samples is combined. This is the sense in which the paper asks whether a taxon is absent: absence from every sample of a large pool is evidence of absence from the population, whereas absence from one sample is mostly evidence about sequencing depth.

**Accumulation curve and asymptote:** The expected number of distinct SGBs detected as a function of how many samples or studies are pooled. Its fitted asymptote estimates how many SGBs the population would yield under unlimited sampling; comparing that asymptote with the catalogue size is how the paper tests for population-level loss.

**VANISH and BloSSUM; the + and H+ lists:** VANISH and BloSSUM are the published lists of taxa depleted and enriched in industrial populations, defined in a Hadza-versus-industrial comparison. $VANISH^{+}$ and $BloSSUM^{+}$ are redefined here on the full industrial-versus-non-industrial contrast, and $VANISH^{H+}$ and $BloSSUM^{H+}$ on an industrial-versus-healthy-industrial contrast. All are lists of taxa with large prevalence differences; none is a list of rare taxa, which is the distinction the paper turns on.

**Geometric-mean-target weighting (GMW):** A reweighting that makes two groups comparable in sequencing depth without discarding samples. Each group is weighted toward a shared target, namely the geometric mean of the two depth distributions, so that samples sequenced where both groups are well represented count more than samples from a depth range only one group occupies. It replaces rarefaction, which throws away reads and is therefore statistically problematic, particularly for an analysis of extinction where it is essential to maximize the available data.

**Effective sample size (ESS):** How many equally weighted samples would give the same precision as the weighted set (Kish's formula). A group of 1,000 samples whose weights are very uneven may carry the precision of only a few hundred, so ESS is reported alongside raw counts wherever weighting is used.

**Prevalence-matched background, and mean $z$:** How much a taxon can shift between cases and controls is bounded by how common it already is: a taxon in 95% of controls can fall but barely rise. Each taxon is therefore scored against the shift expected for non-list taxa of the same baseline prevalence, in units of the local spread ($z$), and a list is summarized by the mean of those scores. A positive mean $z$ means the list is less depleted in cases than comparable taxa, not that it is more prevalent in cases.

**Sample, host, run, and technical replicate:** A run is one sequencing experiment; a sample is one biosample, which may combine several runs; a host is one person, who may contribute several samples over time. Repeated runs of one biosample are technical replicates and are used only for reproducibility checks. Health analyses keep one sample per host, because treating repeat samples as independent would inflate the evidence.

96. X. Zhou, R. Baumann, X. Gao, M. Mendoza, S. Singh, I. Katz Sand, Z. Xia, L. M. Cox, T. Chitnis, H. Yoon, L. Moles, S. J. Caillier, A. Santaniello, G. Ackermann, A. Harroud, R. Lincoln, R. Gomez, A. González Peña, E. Digga, D. J. Hakim, Y. Vazquez-Baeza, K. Soman, S. Warto, G. Humphrey, M. Farez, L. A. Gerdes, J. R. Oksenberg, S. S. Zamvil, S. Chandran, P. Connick, D. Otaegui, T. Castillo-Triviño, S.

L. Hauser, J. M. Gelfand, H. L. Weiner, R. Hohlfeld, H. Wekerle, J. Graves, A. Bar-Or, B. A. C. Cree, J. Correale, R. Knight, S. E. Baranzini, Gut microbiome of multiple sclerosis patients and paired household healthy controls reveal associations with disease risk and course. *Cell* **185**, 3467-3486.e16 (2022).

102. B. W. Silverman, *Density Estimation for Statistics and Data Analysis* (Routledge, New York, 2018).

103. J. G. Sanders, D. D. Sprockett, Y. Li, D. Mjungu, E. V. Lonsdorf, J.-B. N. Ndjango, A. V. Georgiev, J. A. Hart, C. M. Sanz, D. B. Morgan, M. Peeters, B. H. Hahn, A. H. Moeller, Widespread extinctions of co-diversified primate gut bacterial symbionts from humans. *Nat Microbiol* **8**, 1039–1050 (2023).

104. S. Greenland, J. M. Robins, Estimation of a Common Effect Parameter from Sparse Follow-Up Data. *Biometrics* **41**, 55–68 (1985).

105. T. Sato, On the variance estimator for the Mantel-Haenszel risk difference. *Biometrics* (1989).

106. E. H. Simpson, The Interpretation of Interaction in Contingency Tables. *Royal Statistical Society. Journal. Series B: Methodological* **13**, 238–241 (1951).

107. Y. Mundlak, On the Pooling of Time Series and Cross Section Data. *Econometrica* **46**, 69–85 (1978).

108. E. B. Wilson, Probable Inference, the Law of Succession, and Statistical Inference. *Journal of the American Statistical Association* **22**, 209–212 (1927).

109. J. Fan, *Local Polynomial Modelling and Its Applications: Monographs on Statistics and Applied Probability 66* (Routledge, New York, 2018).
